# uPTI: uniaxial permittivity tensor imaging of intrinsic density and anisotropy

**DOI:** 10.1101/2020.12.15.422951

**Authors:** Li-Hao Yeh, Ivan E. Ivanov, Janie R. Byrum, Bryant B. Chhun, Syuan-Ming Guo, Cameron Foltz, Ezzat Hashemi, Juan A. Pérez-Bermejo, Huijun Wang, Yanhao Yu, Peter G. Kazansky, Bruce R. Conklin, May H. Han, Shalin B. Mehta

**Affiliations:** Chan Zuckerberg Biohub, San Francisco, CA, USA; Stanford University, Palo Alto, CA, USA; Gladstone Institutes, San Francisco, CA, USA; University of Southampton, Southampton, UK; University of California San Francisco, CA, USA

**Keywords:** Label-free Microscopy, Inverse algorithms, Density, Anisotropy, 3D orientation, Myelination, Organelle architecture, Respiratory infection

## Abstract

Biological architecture is intrinsically tensorial. The permittivity tensor (PT) of biological material reports the density, angular anisotropy, symmetry, and 3D orientation of biomolecules. High-resolution measurement of PT can enable quantitative and label-free analysis of organelle, cell, and tissue architecture, but remains challenging. We report uniaxial permittivity tensor imaging (uPTI), a label-free computational imaging method for volumetric measurement of PT with diffraction-limited resolution. uPTI encodes the components of PT into intensity modulations using oblique illumination and polarization-resolved imaging. The high-dimensional data is decoded with a vectorial image formation model and a multi-channel convex optimization, assuming that the molecular distribution in each voxel has uniaxial symmetry. We describe a modular implementation of uPTI that can be multiplexed with complementary imaging modalities. We report volumes of uPT in mouse brain tissue, SARS-CoV-2 infected cardiomyocytes, RSV infected A549 cells, H&E stained tissue sections, isotropic beads, and anisotropic glass targets. uPTI enabled volumetric imaging of the 3D orientation and symmetry of organelles, cells, and tissue components with higher spatio-angular resolution than current vectorial tomography, ptychography, and light-field microscopy methods. We provide an open source implementation of the image formation model and reconstruction algorithms.

## Introduction

Label-free imaging of the distribution of biomolecules with electrons, light, and radio waves has enabled multiple biological discoveries. The distribution of biomolecules varies not just in spatial, but also in angular dimensions, i.e., they are tensor quantities. Electron microscopy (EM) reports the distribution of charge density in fixed specimens and provides structural insights with spatial resolution of around a nanometer, but is currently limited to 1 mm^3^ sized tissue (1) despite time and labor-intensive effort. Magnetic resonance imaging (MRI) reports the distribution of hydrogen density and can image dynamic architecture of organs deep into the body, but is currently limited to resolution of 100 μm (2). Label-free light microscopy reports the density and anisotropy of bound electrons described by the permittivity tensor and bridges the resolution gap between EM and MRI. Measurement of permittivity tensor of biological material at spatial scales of 250 nm – 1 cm can reveal architecture of organelles, cells, and tissues from their intrinsic density and anisotropy. These physical properties are complementary to measurement of molecular distribution with fluorescent or histology labels. These measurements can enable new investigations, for example, analysis of myelination and mesoscale connectivity in brain tissue, cytopathic effects of infections, mechanobiology of cytoskeleton and extracellular matrix, image-based fingerprinting of cell types and cell states, and more accurate detection of pathology in histological sections.

Macromolecular components of biological systems (nucleic acids, proteins, lipids, and carbohydrates) form asymmetric assemblies, which underpin the anisotropic functions of organelles, cells, and tissues. The biomolecules are dielectric at visible wavelengths, i.e., the electrons bound to biomolecules oscillate in response to applied electric field, but do not flow. The spatio-angular architecture of biological material is described by the spatial distribution of the permittivity tensor (3), which is the capacity of the material to store electrical energy as a function of the direction and polarization of the incident light. Relative permittivity, whose square root is the refractive index, of a dielectric material quantifies how much the bound electrons are polarized by an applied electric field. Note that the polarization of material is its capacity to store electrical energy, whereas the polarization of light is the plane of oscillation of the electric field. The more easily a material is polarized, the more it delays the electromagnetic wave travelling through it, and higher the permittivity of the material. The polarization of anisotropic material depends on the direction and the polarization of incident light, i.e., the refractive index of the anisotropic material is a function of the direction and the polarization of the electric field. The anisotropy of the material is measured in terms of the polarization-dependent refractive index (birefringence) or the polarization-dependent phase delay (retardance). If the bound electrons resonate with the incident optical frequency, the material absorbs the light. At visible wavelengths, biological specimens exhibit substantial phase and retardance, but negligible absorption, which makes them transparent.

The angular distribution of macromolecules over each voxel can result in three distinct refractive indices along the principal axes of anisotropy. However, many biological structures, such as axon bundles, collagen fibers, filaments of cytoskeletal and motor proteins, plasma membrane, nuclear envelope, mitochondria, etc., are assembled with a single symmetry axis that results in two distinct refractive indices, ordinary index perpendicular to the axis of symmetry and extraordinary index parallel to the axis of symmetry. Such structures can be described by a uniaxial permittivity tensor (uPT), which is a rank-3 tensor with two of three eigenvalues being the same. Throughout this paper, we assume that the biological material has uniaxial symmetry, which is a correct assumption for a large range of structures with the isotropic, linear or planar symmetries. When the permittivity tensor is biaxial, i.e., has 3 distinct eigenvalues, our method provides a reliable measurement of its uniaxial component.

The permittivity tensor (PT) of a complex specimen can be decomposed into the isotropic component that reports the local density of macromolecules and the anisotropic component that reports the local angular distribution of macromolecules. The PT is mathematically analogous to, but physically distinct from, the diffusion tensor (DT) that is commonly measured with diffusion tensor imaging (DTI). PT reports the architectural symmetries at the scale of cells and tissues, just as DT reports the symmetries of axon distribution in the brain. The isotropic component of PT is similar to the mean diffusivity component of DT, the anisotropy of PT is similar to the fractional anisotropy component of DT, and the 3D orientation of anisotropy of PT is similar to the 3D orientation of the axial diffusivity of DT (4). The imaging resolution and depth of DTI and PTI, however, are quite different. DTI is suitable for organ to tissue level of imaging deep inside the specimen’s body, while PTI is suitable for organelle, cell, and tissue level of imaging up to the depth of few tens of microns.

Majority of the quantitative label-free light microscopy methods either image the isotropic component or the anisotropic component of uPT projected on the focus plane. Quantitative phase microscopy images the isotropic component of uPT by measuring the density (optical path length) and has been used for imaging of organelles (5, 6). Optical diffraction tomography (7, 8) and shearing interferometry methods (9) that account for diffraction effects measure density with spatial resolution of 0.2 × 0.2 × 0.4 μm^3^ and 0.3 × 0.3 × 1.1 μm^3^, respectively. On the other hand, quantitative polarization microscopy images the anisotropic component of permittivity tensor (the retardance and the orientation) and has been used to study microtubule spindle (10, 11), white matter in human brain tissue slices (12–15), collagen architecture in the eye tissues (16), etc. Phase and polarization imaging are also used to quantify the optical properties of fabricated materials (17, 18). The isotropic and anisotropic components of uPT are induced simultaneously when light interacts with the matter, but they are typically not measured simultaneously.

Development of methods that provide more complete measurement of uPT to analyze a larger set of biological structures is an active area of research. A complete description of uPT consists of 3D anisotropy, isotropic density, and the material symmetry (optic sign) in 3D space. Since we are reporting measurements in spatio-angular coordinates, the terms 2D/3D anisotropy and orientation are used to imply angular dimensions, whereas 2D plane/3D volume are used to imply spatial dimensions. Existing approaches measure uPT with various degrees of completeness. Among these, several methods employ geometric image formation models and do not achieve the diffraction-limited resolution. Shribak et al. reported a method combining orientation-independent differential interference contrast microscopy (OI-DIC) and quantitative polarization microscopy that provides joint density and 2D anisotropy (retardance and orientation projected on the image plane) measurements in a 2D plane, without considering diffraction effects (19). Ptychography-based phase retrieval method has been extended with polarization sensitive components and Jones calculus for joint imaging of density and 3D anisotropy (principal retardance and 3D orientation) in a 2D plane considering diffraction (20, 21). Our recent work, quantitative label-free imaging with phase and polarization (QLIPP), combines phase from defocus (22–24) with quantitative polarization microscopy (25–27) to measure density and 2D anisotropy (28) in 3D volume. A recent work has extended this idea to provide the measurement analogous to QLIPP, but with 3D diffraction-limited resolution (29). These approaches have not achieved a complete measurement of uPT with 2D and 3D diffraction-limited resolution or demonstrated measurement of uPT in diverse biological systems. The complete measurement of uPT in diverse biological systems would enable more quantitative, specific, and reproducible analysis of biological structure.

We report uniaxial permittivity tensor imaging (uPTI), a computational microscopy technique, which provides diffraction-aware measurements of complete uPT including absorption, phase, principal retardance, 3D orientation, and symmetry in 2D and 3D space for the first time. uPTI captures these properties of the specimen by combining oblique illumination (30–32) with polarization-sensitive detection (12, 26, 33–36). We implement this design on a research microscope with inexpensive addon modules. We develop a vectorial imaging model and the corresponding multi-channel deconvolution algorithm to extract spatial distribution of the components of the uniaxial permittivity tensor. We validate the accuracy and resolution with isotropic polystyrene beads and anisotropic laser-written glass targets. We demonstrate that uPTI allows analysis of the architecture of the mouse brain at scales of whole slice, axon bundles, and single axons. We show that uPTI measurements can be multiplexed with fluorescence deconvolution microscopy to enable analysis of the physical and molecular architecture of organelles in SARS-CoV-2 infected iPSC-derived cardiomyocytes and RSV-infected A549 cells. Finally, we show that uPTI can add new information when multiplexed with H&E stained histological sections. These data establish a novel label-free measurement technology for comprehensive high-resolution imaging of biological architecture. With our inexpensive optical design and open source software, we aim to enable rapid adoption and refinement.

## Results

### A: Computational imaging concept

#### Light path

Figure 1A and Figure 1-supplement 1 show the optical setup of uPTI. Video 1 shows the building process of the corresponding setup. We implemented uPTI on a Leica DMi8 inverted microscope with two add-on modules, an oblique illuminator and a polarization imaging module. The oblique illuminator is composed of a green color filter, a linear polarizer, a programmable amplitude modulator, a right-hand circular polarizer, and a condenser lens. The light from an LED source is first filtered by a green filter and the linear polarizer before passing through the amplitude modulator placed in the front focal plane of the condenser lens. The amplitude modulator is constructed from a low-cost liquid crystal display (LCD, Adafruit, ST7735R). The right-hand circular polarizer (RCP, Thorlabs, CP1R532) is placed after the amplitude modulator. This module enables computer-controlled oblique illumination with circularly polarized light. It is compact enough to be placed near the front focal plane of a high-NA condenser and allows illumination of the specimen with the numerical aperture (NA) as high as 1.4 with high light coupling efficiency. The oblique circularly polarized light then impinges on the specimen, generates scattered light, and is collected by the polarization imaging module. The polarization imaging module is formed by the microscope objective, tube lens, and a four-channel polarization sensitive camera (FLIR, BFS-U3-51S5P-C). The polarization sensitive camera has a patterned grid of linear polarizers on top of pixels, with transmission axes along 0°, 45°, 90°, and 135°. The camera images four linearly polarized light states with a single exposure. A microscope with this camera can capture the projected retardance and 2D orientation of material’s slow axis similar to other polarized light microscopes (12, 27, 28, 33, 36). The module that enables acquisition of 3D anisotropy is the high-quality oblique illuminator. Overall, our modular design enables tomographic polarization imaging of specimens with diverse oblique illumination patterns.

**Fig. 1.**
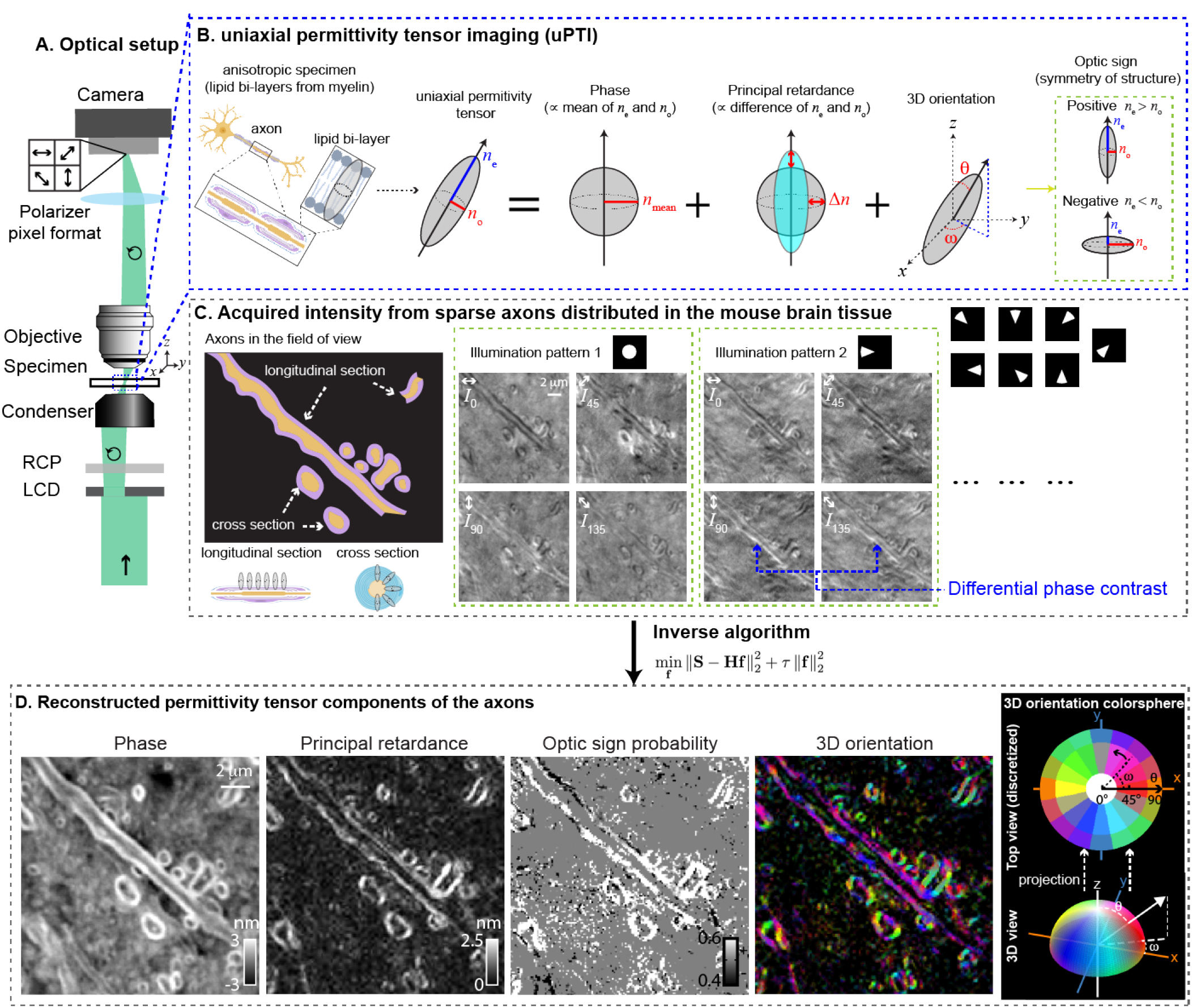
Measurements with the uniaxial permittivity tensor imaging (uPTI): **(A)** Light path of the microscope. **(B)** Optical anisotropy arises from the angular anisotropy of biological architecture, e.g., ordered lipid bilayers in the myelinated axons give rise to a higher refractive index perpendicular to the bilayer (created with BioRender.com). The optical anisotropy can be represented by a uniaxial permittivity tensor (uPT) or an indicatrix, which is an ellipsoidal surface that represents angular distribution of the optical path length over the spatial resolution of the imaging system. The uPT is parameterized by its 3D orientation (*ω* and *θ*) and two principal axes (*n_e_* and *n_o_*). The uPT can be decomposed and interpreted in terms of four physical quantities: the average delay in the light induced by the specimen (phase), the degree of anisotropy (principal retardance), 3D orientation, and the symmetry of the structure (optic sign, *n_e_* ≷ *n_o_*). **(C)** An example field of view with longitudinal sections and cross sections of axons (wrapped with myelin sheaths) illustrates the dependence of intensity contrast on the polarization and the angle of illumination. We observe intensity modulations due to properties of axons in the acquired intensities. Anisotropy variations are observed in different polarization channels, while phase variations modulate intensities across illumination patterns (illumination pattern 1 vs 2). We also observe contrast variations in different polarization channels across on-axis and off-axis illumination due to variations in the out-of-plane orientation and the optic sign. **(D)** Using an inverse algorithm based on convex optimization, we reconstruct phase, principal retardance, optic sign probability, and 3D orientation of the axons in the example field of view from intensities. The 3D orientation image shows false-color images in which the 3D orientations is shown by the color and the principal retardance is shown by the brightness of the color. We assign a color to the 3D orientation using a spherical colormap adopted from (37) (e.g. *ω* = 0° and *θ* = 45° for red, *ω* = 180° and *θ* = 45° for yellow). The corresponding color sphere is visualized in its 3D view and its projected top view with discretized color patches for ease of reference. We report the 3D orientation of the symmetry axis of the permittivity tensor, independent of its optic sign.

#### Interpretation of uPT

Figure 1B illustrates how 3D anisotropy of a lipid bi-layer structures is described by a uniaxial permittivity tensor (uPT) or an indicatrix, an ellipsoidal surface parameterized by: 1. 3D orientation (*ω* and *θ*) of the symmetry axis, 2. the ordinary refractive index (*n_o_*) experienced by the waves with electric field in the directions perpendicular to the symmetry axis, and 3. the extraordinary refractive index (*n_e_*) experienced by the waves with electric field in the direction parallel to the symmetry axis. The uPT can be interpreted through four physical components: 1. average delay in the light induced by the specimen (phase proportional to the mean of *n*_e_ and *n*_o_), 2. degree of anisotropy (principal retardance proportional to the difference of *n*_e_ and *n*_o_), 3. 3D orientation of the symmetry axis or optic axis of the material (azimuthal angle *ω* and inclination *θ*), and 4. symmetry of the structure around the optic axis (optic sign, *n*_e_ ≷ *n*_o_). Materials are referred to as positive or negative uniaxial depending on their optic sign (3). For example, the spatially resolved lipid bi-layer is a positive uniaxial material, whereas the anisotropy of the test target (fig. 2) and whole axons reported later in fig. 3 are negative uniaxial.

**Fig. 2.**
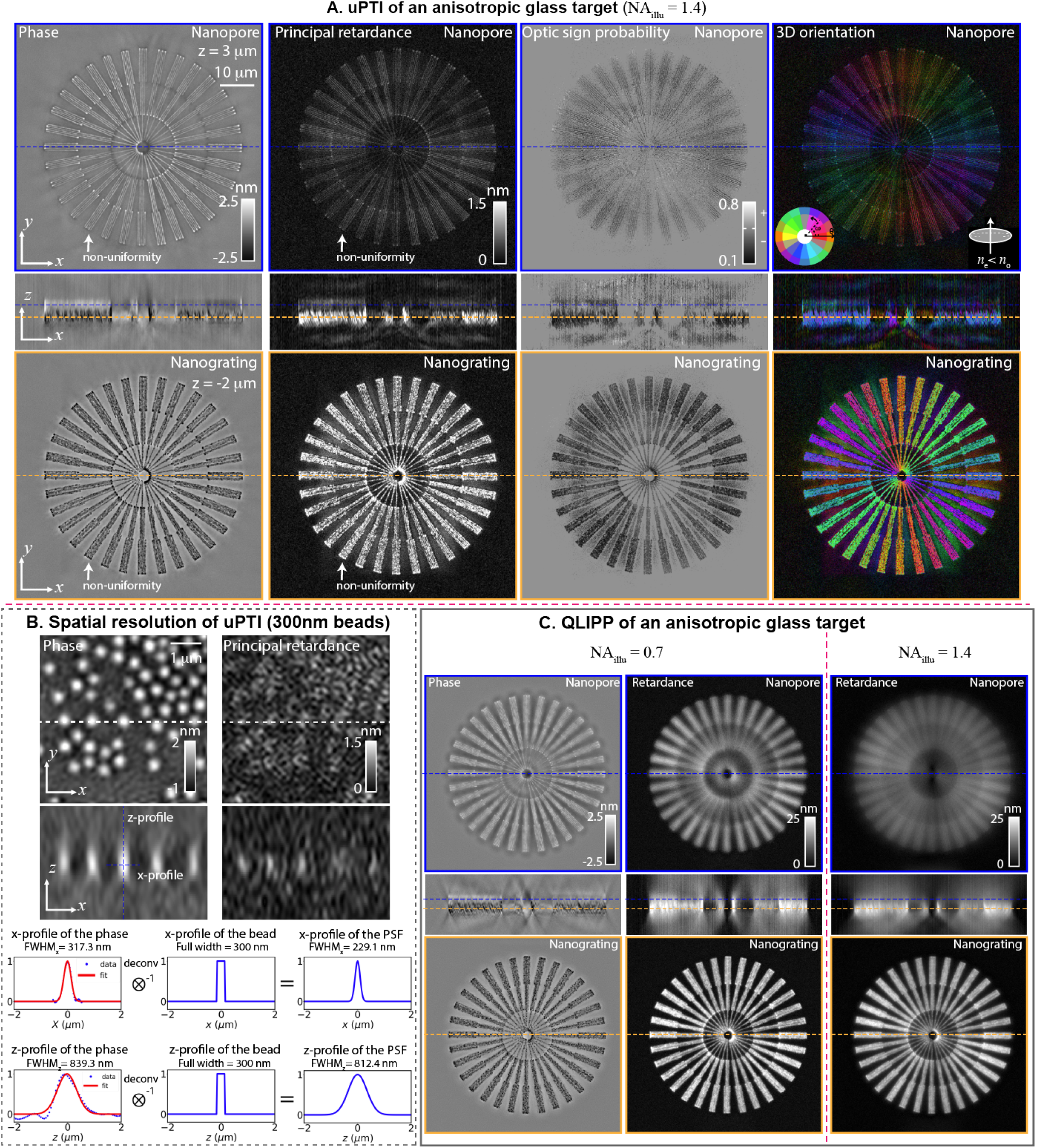
3D spatial resolution of uPTI: **(A)** *x-y* and *x-z* sections of phase, principal retardance, optic sign probability, and 3D orientation volumes of a laser written anisotropic glass enable identification of nanograting and nanopore structures at different depths. Nanopore layer is shown at the top and indicated by blue dashed line in *x-z* sections. Nanograting layer is shown at the bottom and indicated by orange dashed line in *x-z* sections. **(B)** Characterization of 3D spatial resolution by imaging 300 nm polystyrene beads with refractive index of 1.5956 immersed in oil with refractive index of 1.5536. The phase images of beads show dense center and principal retardance images resolve edges of the beads. The phase of the center bead is selected for Gaussian fits in *x* and *z* directions. The Gaussian fits are deconvolved with the physical size of the bead to measure the full width at half maximum (FWHM) of the point spread function (PSF) in *x* and *z*. **(C)** *x-y* and *x-z* sections of the phase and retardance of the same target measured using QLIPP with two different illumination NAs (0.7 and 1.4) show spatial resolution and contrast lower than in uPTI measurements. 3D orientation and optic sign are not accessible with QLIPP.

**Fig. 3.**
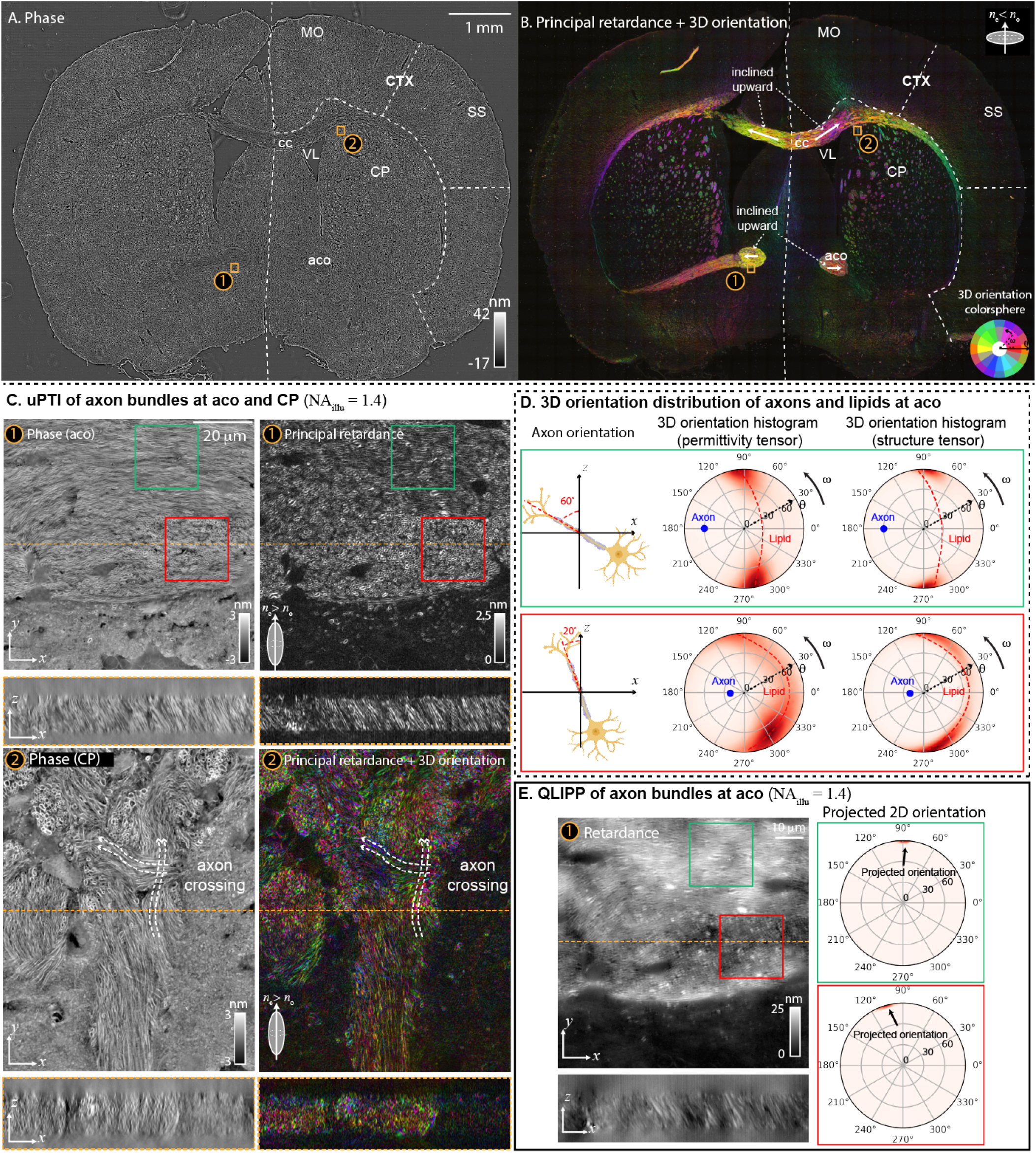
Multi-scale imaging of the architecture of an adult mouse brain section with uPTI: **(A)** phase, **(B)** principal retardance and 3D orientation images of the whole brain section show key anatomical landmarks. At the imaging and illumination NA of 0.55 (spatial resolution of ~ 0.5 × 0.5 × 3.2 μm), axons behave like negative uniaxial material (see Results). Therefore, we assume the negative uniaxial material when analyzing 3D orientation over the whole slide. Multiple anatomical landmarks are labeled according to the coronal section at level 51 of the Allen brain reference atlas, aco: anterior commissure (olfactory limb), cc: corpus callosum, CP: caudoputamen, CTX: cortex, MO: motor cortex, SS: somatosensory area, VL: ventricle. **(C)** aco and CP areas marked with orange boxes (labeled with ① and ②) in (A) and (B) are selected for 3D high-resolution imaging with imaging NA of 1.47 and illumination NA of 1.4 (spatial resolution of ~ 0.23 × 0.23 × 0.8 μm). The orthogonal sections of phase, principal retardance, and 3D orientation indicate complex axon networks in the mouse brain tissue. With high resolution, we resolve the boundaries of individual axons, which behave as the positive uniaxial material with 3D orientation perpendicular to the surface of the membrane. **(D)** We compare 3D orientation distribution of the measured permittivity tensor and computed structure tensor (from phase) of two volumes identified with green (axons tilted 60° to the left from *z*-axis) and red (axons tilted 20° to the left from *z*-axis) boxes. The histograms show that the distributions of the 3D orientation of lipids from permittivity tensor and structure tensor match well in the two cropped volumes. The blue dot in each histogram indicates the corresponding axon orientation in the selected volume. **(E)** Orthogonal sections of the retardance of the same volume at aco measured using QLIPP with the illumination NA of 1.4 show spatial resolution and contrast worse than uPTI measurements. Only projected 2D orientation is accessible with QLIPP, i.e., inclination component of 3D orientation is not accessible.

Since the 3D orientation reports the symmetry axis of the material in each voxel, it aligns with the slow axis of the material, i.e,. axis with higher refractive index, when the material is positive uniaxial. 3D orientation aligns with the fast axis of the material, i.e., axis with lower refractive index, when the material is negative uniaxial.

#### Image acquisition and simulation

To measure the uPT distribution, we image our specimen with diverse oblique illumination and four linear polarization states. The choice of illumination patterns projected on the LCD is discussed in Supplementary Note 3 and Figure 1-supplement 2. Following fig. 1B, we illustrate how the material properties of axons are transferred to intensities in fig. 1C. Figure 1C shows raw images from an example field of view containing longitudinal sections and cross sections of axons in the mouse brain tissue section. The sample is illuminated with 1.4 NA, and imaged with objective with 1.47 NA. Under both the circular (illumination pattern 1) and the sector (illumination pattern 2) illumination patterns on the LCD, we see strong intensity modulations due to the anisotropic myelin sheaths (multi-layer lipid bi-layers) in four polarization channels. When the illumination pattern is a sector (illumination pattern 2), edges of the middle longitudinal axon (indicated by blue arrows) show an intensity gradient perpendicular to the axon in addition to the polarization intensity modulation, demonstrating the multiplexing of differential phase contrast (30–32) and the polarization contrast. We show a similar illustration with a wave optical simulation of images Figure 1-supplement 3 of a target that consists of an isotropic spoke pattern and two anisotropic spoke patterns with defined 3D orientation, uniaxial symmetry, and opposite optic signs. In the simulation, the differential phase contrast is more visible in both the isotropic material and the anisotropic materials under a sector illumination. We also observe that the anisotropic spoke patterns cause differential intensity modulations in the four polarization channels when the on-axis (brightfield) and the off-axis (sector) illuminations are used. This is due to the difference in the optic sign and the 3D orientation of optic axis of the material (34).

For a 2D specimen, we use 36 2D images (9 oblique illuminations with 4 polarization channels) for data reconstruction. If the specimen is 3D in nature, i.e., the thickness of the specimen is larger than the depth of field of the microscope, instead of collecting intensities in a 2D plane, we collect 36 3D *z*-stacks (9 oblique illuminations with 4 polarization channels at each plane). To account for background polarization effects introduced by components in the optical path other than the specimen, we also collect a dataset (36 2D images from 9 oblique illuminations with 4 polarization channels) at an empty field of view, which is used in the reconstruction of the physical properties of the specimen (see Supplementary Note 2).

#### Reconstruction of uPT

Extraction of components of uPT from measured intensities considering diffraction requires a vectorial partially coherent imaging model that expresses intensities in terms of the components of uPT. We develop and summarize this model in Method A which expresses distribution of uPT as a scattering potential tensor and maps the scattering potential tensor to the Stokes parameters measured in the image plane. Scalar scattering potential is a scaled difference of the refractive indices of the specimen and the surrounding medium, which is a key concept employed in the diffraction tomography of 3D refractive index (density of bound electrons) (38). An extension of scattering potential, 2 × 2 scattering potential tensor, has been developed (29, 39) for volumetric reconstruction of projected anisotropy. Our work generalizes the formulation to measure a more complete 3 × 3 scattering potential tensor, which allows reconstruction of volumetric distribution of density, 3D anisotropy, and material symmetry. The mapping between scattering potential tensor and intensities recorded by uPTI is nonlinear. The non-linearity of the mapping makes the inverse problem challenging, because it can create ambiguities when searching for a unique solution. Fortunately, this mapping can be linearized when the scattering is weak enough such that the single-scattered photons contribute to most of the intensity variations. With this approximation, which corresponds to the first Born and weak object approximation (3), we build a forward model. Our forward model describes a linear relationship between the Stokes parameters and the scattering potential tensor via the transfer functions that depend on the parameters of the microscope. This forward model enables the development of a deconvolution algorithm that retrieves the uPT from high-dimensional acquisition. Before reconstructing the components of the uPT, we first calibrate the mapping between Stokes parameters and the measured intensities, convert the measured intensities into the Stokes parameters, and perform a background correction. The deconvolution algorithm (Method B) converts the Stokes parameters into physical properties summarized in fig. 1B. This computational framework allows us to transform the input intensities from fig. 1C into diffraction limited measurements of phase, principal retardance, optic sign probability, and 3D orientation of the specimen as shown in fig. 1D. The 3D orientation is represented by mapping the azimuthal angle and inclination to a color sphere introduced in fig. 1D. Since anisotropy measurement is usually noisier than the phase channel, we developed various denoising methods for uPT as described in Method D and applied to data reported in this paper.

The python software that implements the forward model, the reconstruction algorithm, the simulations, and illustration of reconstruction is available at https://github.com/mehta-lab/waveorder.

### B: Characterization of spatial resolution and accuracy

As illustrated in fig. 1, uPTI provides four distinct volumetric measurements: phase, principal retardance, 3D orientation, and the optic sign. uPTI is designed to use high NA partially coherent illumination and high NA imaging, which enables confocal-like depth sectioning in measured channels. To validate the accuracy and diffraction-limited volumetric resolution achievable by uPTI for these channels of information, we image three types of isotropic and anisotropic test targets. All uPT measurements reported in this section are acquired with an 1.4 NA (NA_c_) oil immersion condenser and 1.47NA (NA_o_) oil immersion objective.

#### 3D imaging of anisotropic glass target

First, we image a laser-written anisotropic glass target shown in fig. 2A (through-focus video is shown in Video 2) to characterize the 3D orientation, verify estimation of optic sign, and demonstrate the utility of uPTI for metrology. The anisotropic target is made of fused silica modified with a polarized femtosecond laser focused with 0.55 NA lens (Method G) (40). With uPTI, we identify two distinct laser-induced modifications: nanograting (41) and nanopore (18) at different axial layers of the material. Reading these two types of modifications along the depth has been challenging with current methods, including QLIPP (fig. 2C). According to (18), nanograting modification of the material generates negative phase and stronger retardance, while nanopore modification generates positive phase and weaker retardance. uPTI estimates the target to have high probability of being negative uniaxial material, which agrees with past observations (17, 18). The *x-y* and *x-z* sections through phase, principal retardance, and optic sign volumes match with these expected optical properties. Since this is a negative uniaxial material, the 3D orientation of the optic axis reports the fast axis of the material. We show 3D orientation in fig. 2A using the color sphere shown in fig. 1D. The orientation of the optic axis in each spoke aligns with the *x-y* plane and is orthogonal to the direction of the spokes, which matches with the axis of symmetry expected from the state of laser polarization used in the writing process. We clarify the same data with a RGB color sphere, which encodes 3D orientation by assigning RGB colors proportional to the projection of 3D orientation on *x*, *y*, and *z* axes in Figure 2-supplement 1. In addition, we measure subtle non-uniformity in phase and principal retardance at the ends of the line features along each spoke (shown with arrows).

Another anisotropic target fabricated with different writing parameters is shown in Figure 2-supplement 2 (through-focus video is shown in Video 3). This target has only one layer of nanograting modification. The double line-scan process used for writing this target eliminated the non-uniformity. Collectively, these measurements show that uPTI can be a valuable technology for metrology and for reading novel optical storage devices, in addition to its use in biological imaging.

#### Spatial resolution

Next, we characterize the spatial resolution of uPTI by imaging 300 nm polystyrene beads with refractive index (RI) of 1.5956 embedded in oil with the RI of 1.5536 (fig. 2B). In the phase image, we can resolve individual beads. In the principal retardance image, we can resolve edge retardance of the beads in the form of small rings. To quantify the resolution, we select the center bead in the phase image for Gaussian fits in *x* and *z* directions. Deconvolving the physical size of the bead from the fitted Gaussians, we obtain the shape of the point spread functions (PSFs) in *x* and *z* directions. The full width at half maximum (FWHM) of this PSF in *x* and *z* directions show that we achieve a transverse FWHM of 230 nm and an axial FWHM of 810 nm. We use FWHM of a theoretical image of a point (3) with a lens of 1.4 NA to benchmark the transverse and axial resolutions. The theoretical transverse FWHM is 190 nm (0.51λ/NA) and the axial FWHM is 543 nm (2λ/NA^2^). The theoretical axial resolution is a function of transverse spatial frequency of the specimen (22). For a spherical bead of finite spatial frequency, the FWHM can be worse than an infinitesimal point. Our measured transverse FWHM and axial FWHM compare well with the theoretical quantities. These results also illustrate that our deconvolution algorithm and parameters do not introduce artifacts. As illustrated by images in fig. 3-6, our measurements provide confocal-like 3D resolution that allows us to resolve crosssections of single axons, bands of sarcomeres and intracelluar features.

We compare the resolution of uPTI with our previous method, QLIPP (28). Phase and projected retardance of the anisotropic glass target measured with QLIPP are shown in fig. 2C with illumination NA of 0.7 and 1.4. Since phase information is not accessible through defocused intensity stack acquired with QLIPP when the illumination NA is close to the imaging NA, Figure 2C only shows QLIPP phase image of the target with illumination NA of 0.7. uPTI phase image has higher *xyz* resolution than QLIPP phase image as evident from lines within each spoke of the target and the sharper features in the *x-z* section.

The *x-z* sections of the principal retardance measured with uPTI show higher resolution than projected retardance measured with QLIPP retardance at both illumination NAs. uPTI provides optical sectioning needed to distinguish two layers of material modifications separated by 1 μm. The fine spacing (300 nm) inside individual spokes are better resolved in uPTI than in QLIPP. Thus, anisotropy measurements with uPTI approach the diffraction-limit.

#### Accuracy

Finally, we characterize the accuracy of phase and principal retardance by imaging isotropic 3 μm polystyrene beads embedded in oils of varied RIs as shown in Figure 2-supplement 3. Embedding the beads (*n*_beads_=1.5956) in the media of varying RI (*n*_media_ ranges from 1.5536 to 1.5826) changes the accumulated optical path length (theoretical phase) of the light as well as the amount of edge retardance (42) linearly. When the RI of the surrounding media is the same as the RI of the beads, there will be no accumulated phase and edge retardance. Such an embedding series allows us to characterize the linearity of phase and principal retardance measured with uPTI. In the phase images of the beads, we see that phase values drop as the RI of the immersion oil approaches the RI of the bead. We see similar trend in the edge retardance signal from the principal retardance measurements. Plotting the theoretical phase and the measured phase versus the difference of the RI between beads and oils, we find the measured phase matches well with the theoretical one. We also see that edge retardance varies linearly with the difference of the refractive index between beads and oils, which is in agreement of the measurement from (42).

### C: uPTI enables multi-scale analysis of the architecture of mouse brain tissue

A key limitation of current polarized light microscopy (28, 29, 39) approaches has been that their light paths are not sensitive to the inclination of the 3D anisotropy. As a result, they report anisotropy projected on the microscope’s image plane. Polarization microscopy with scanned illumination aperture (43) and light-field (44) detection are sensitive to the inclination, but do not have diffraction-limited resolution, because they don’t account for diffraction effects. So far, measurement of principal retardance and 3D orientation of anisotropic structures with diffraction-limited resolution has been an unsolved problem. With the diffraction-aware measurement of permittivity tensor, we reasoned that uPTI can measure the architecture of the brain tissue and enable high-resolution imaging of principal retardance and 3D orientation of axons.

The architectural connectivity of mammalian brains can be inferred from the spatio-angular distribution of myelinated axons. Myelin sheath is composed of multiple lipid bi-layers and wraps around axons. As fig. 1B illustrates, the electrons bound to lipid molecule are polarized more (have higher RI) along the orientation of molecules and less (have lower RI) in the plane of bi-layer, which results in an angular refractive index distribution of a positive uniaxial material. When the light scattered by the myelin sheath is integrated around the axon cross-section, the bound electrons are seen to have higher RI perpendicular to the axon axis, resulting in an angular RI distribution of a negative uniaxial material. The myelination in brain tissue can be measured in terms of the principal retardance and the optic sign of the uPT. Further, the 3D orientation measured at the resolution of diameter of single axons (~ 1 μm) can enable analysis of the complex connectivity within brain regions.

MRI can provide measurement of spatio-angular distribution of axon bundles (4) and myelin fraction (45) with millimeter resolution. However, inference of the connectivity or pathology frequently requires micro-architectural ground truth (45, 46). Polarization microscopy is emerging as a label-free method for analyzing mesoscale connectivity and the architecture of brain tissue (12–15, 28, 47) due to the following reasons: 1) High intrinsic anisotropy of the myelin sheath enables sensitive detection of distribution and orientation of axon fibers (48, 49), 2) Light microscopy can achieve sub-micron, single-axon resolution across large brains. Quantitative phase microscopy has also enabled imaging of brain architecture (50, 51). Here, we report measurements of phase, principal retardance, and 3D orientation at spatial scales ranging from 1 cm–1 μm in 12 μm thick sections of brain slices. At high resolutions, we acquire volumetric measurements and at low resolutions, we acquire planar measurements.

#### 2D imaging of whole section

First, we report planar (2D) measurements of phase, principal retardance, and 3D orientation of a section of adult mouse brain tissue. Figure 3A and 3B show the phase, principal retardance, and 3D orientation of an adult mouse brain located at level 51 of the Allen brain reference atlas (https://mouse.brain-map.org/static/atlas). With the imaging and illumination NA of 0.55 (corresponding to spatial resolution of ~ 0.5 × 0.5 × 3.2 μm), the imaging system measures anisotropy of myelin sheath averaged over whole axons. As a result, axons behave like a negative uniaxial material (Figure 3-supplement 1B) with 3D orientation (fig. 3B) is co-linear with axon axis (48, 49). Therefore, we assume all the axons are negative uniaxial material when computing 3D orientation at this resolution. Principal retardance and 3D orientation are rendered together with the brightness encoding the principal retardance and color encoding the 3D orientation as indicated by the color sphere. Phase shows overall morphology of the mouse brain, while principal retardance highlights the distribution of myelinated axons. As in other work (28, 47), important anatomical regions such as anterior commissure olfactory limb (aco), corpus callosum (cc), caudoputamen (CP), cortex (CTX), and ventricle (VL) are visible in both phase and principal retardance. In the 3D orientation image, we not only see the in-plane orientation aligned with the axon bundle, but also see a smooth transition in inclination from left to right over corpus callosum (yellow to pink colored region indicated by the top two white arrows in fig. 3B). We also notice that the left and right anterior commissure olfactory limb are inclined relative to the microscope axis (yellow and red colored stretches indicated by bottom two white arrows in fig. 3B). The same out-of-plane inclinations are also visible in bluish hue when the 3D orientation is encoded using RGB colored sphere Figure 3-supplement 1A.

#### Volumetric, high resolution imaging of brain regions

Next, we report high-resolution analysis of the brain tissue. Figure 3C shows *x-y* and *x-z* sections of the aco and CP regions imaged at high-resolution (1.47 NA, spatial resolution of ~ 0.23 × 0.23 × 0.8 μm) in the section described above. Corresponding scans through *x-y*, *x-z*, *y-z* sections are shown in Video 5-Video 6. At high resolution, uPTI measurements resolve myelin sheath around individual axons, which behaves like a positive uniaxial material with 3D orientation normal to the membrane. We provide another visualization of the 3D orientation with the RGB colored sphere and the optic sign probability in Figure 3-supplement 1. In these fields of view, longitudinal sections and cross sections of axons are visible in both phase and principal retardance channels, suggesting axons at these regions are partially parallel with the focal plane and partially parallel with the microscope imaging axis. We also observe multiple axon crossings in the field of view of fig. 3-②.

To further validate the measurements of the 3D orientation of lipids from the anisotropic part of the permittivity tensor, we compare it with the 3D orientation of lipids computed from the structure tensor of the isotropic part, the phase image (Method F). Figure 3D shows the histogram of the 3D orientation of lipids in two sub-volumes (with different axon inclinations) from the permittivity tensor and the structure tensor of the phase image. The azimuthal dimension of this histogram shows the in-plane orientation, *ω,* and the radial dimension shows the inclination relative to the imaging axis, *θ.* The green box contains axons mostly inclined at 60° to the left of the *z*-axis and the red box contains axons mostly inclined at 20° to the left of the *z*-axis. We indicate the axon orientation with blue dots in the histograms of 3D orientation. If the inclination of the axon is 0° from the *z*-axis, we expect 3D orientation of lipids to be in the focal plane evenly distributed in the azimuthal dimension, which will be a distribution around a circle with radius *θ* = 90°. With 20° and 60° inclinations of axons, we expect gradual rotation of this circle (collective 3D orientation of lipids) to the left side of the histogram, which is what we observed in 3D orientation histograms from both the permittivity tensor and the structure tensor. At high inclinations of the axon, we notice a gradual reduction in the density of orientations of lipids as lipids align along the *z*-axis. This drop in sensitivity is due to the poor transfer of optical properties of anisotropic material to intensity modulations as the material aligns with *z*-axis.

uPTI provides high-resolution images of axon networks in diverse channels of physical properties owing to its diffraction-aware model that accounts for wave-optical effects from a complete uniaxial permittivity tensor. Imaging at the same field of view shown in fig. 3C-① with QLIPP, fig. 3E shows the *x-y* and *x-z* sections of a 3D retardance stack and the corresponding 2D orientation measurements on the histogram of 3D orientation in the green box and red box areas of axons. Axon boundaries are barely visible in the QLIPP measurements due to lower resolution and contrast. The inclination and optic sign are accessible with uPTI, but not accessible with QLIPP.

#### Multi-resolution analysis

Finally, we automate multi-scale imaging to acquire uPT measurements of mm-sized tissue sections with sub-micrometer 3D resolution. We automate tiled acquisition using Micro-Manager (https://github.com/micro-manager), a python bridge to Micro-Manager (https://github.com/czbiohub/mm2python), and GPU-accelerated computational pipeline implemented on a compute cluster (Method E). We designed the analysis pipeline to enable robust reconstruction of uPT at any scale spanned by the acquisition. Measurements at larger scales (lower resolution) are computed by a spatially filtering approach (Method C). Results of one such multi-scale analysis of the right corpus callosum region are shown in Video 4. At spatial scales larger than the typical size of axons, we compute the 3D orientation assuming a negative uniaxial material. When axon cross-sections are resolved, we visualize complex axon networks by displaying the phase and principal retardance through focus and at multiple locations.

We verify the quantitative correspondence between 3D orientation distributions measured with low-resolution (20x, 0.55NA) and high-resolution (63x, 1.47NA) acquisitions. We image the 3D orientation in fig. 3-① at high-resolution. We low-pass filter the high-resolution data (Method C) to have similar spatial resolution as the low-resolution data and compute the 3D orientation histogram within two sub-regions as shown in Figure 3-supplement 2. The histograms of 3D orientation of axon bundles in the low-resolution data and the smoothed 3D orientation computed from high-resolution data agree well, confirming that our pipeline provides physically meaningful measurements across spatial scales. These results also indicate that uPTI provides sensitive measurement of 3D anisotropy that cannot be resolved from the spatial architecture, and uPT measurements with 1 μm resolution can be used for rapid, quantiative analysis of the distribution of axons in different regions of brain.

### D: 3D imaging of cytopathic effects using uPTI multiplexed with fluorescence deconvolution microscopy

Quantitative label-free imaging provides unbiased and consistent readouts of physical architecture of diverse cell types, including human cells and tissues. Immunolabeling, on the other hand, provides complementary information about the distribution of specific molecules. To map both physical and molecular architecture of cells at confocal-like spatial resolution, we designed and implemented uPTI to enable multiplexing with a wide-field fluorescence deconvolution imaging. uPTI’s design permits use of the highest NA illumination and imaging lenses, which allows us to achieve diffraction-limited 3D resolution. We use uPTI multiplexed with fluorescence to analyze cytopathic effects in two cellular models of infection, and demonstrate that uPTI can reveal impacts of perturbations such as infection at cellular and organelle scales. Infection typically changes the architectural and functional states of host cells. Diffraction-limited measurements of uPT in models of infection is a promising tool for discovery of host cell states throughout the infection cycle.

#### Cytopathic effects of SARS-CoV-2 infection on iPSC-derived cardiomyocytes

Induced pluripotent stem cell (iPSC)-derived cardiomyocytes (CMs) have emerged as genetically editable models of cardiac diseases and drug screening (52). CMs are highly specialized contractile cells. Studying the architecture of the myofibril and its building blocks, the sarcomeres, is of critical importance to characterize their function (52). Polarized light imaging has played an important role in understanding architecture and activity of sarcomeres (53) (fig. 4B). In fact, the A- and I-bands of sarcomeres were named after the anisotropic and isotropic bands first observed in muscle tissue with polarized light microscopy (54).

**Fig. 4.**
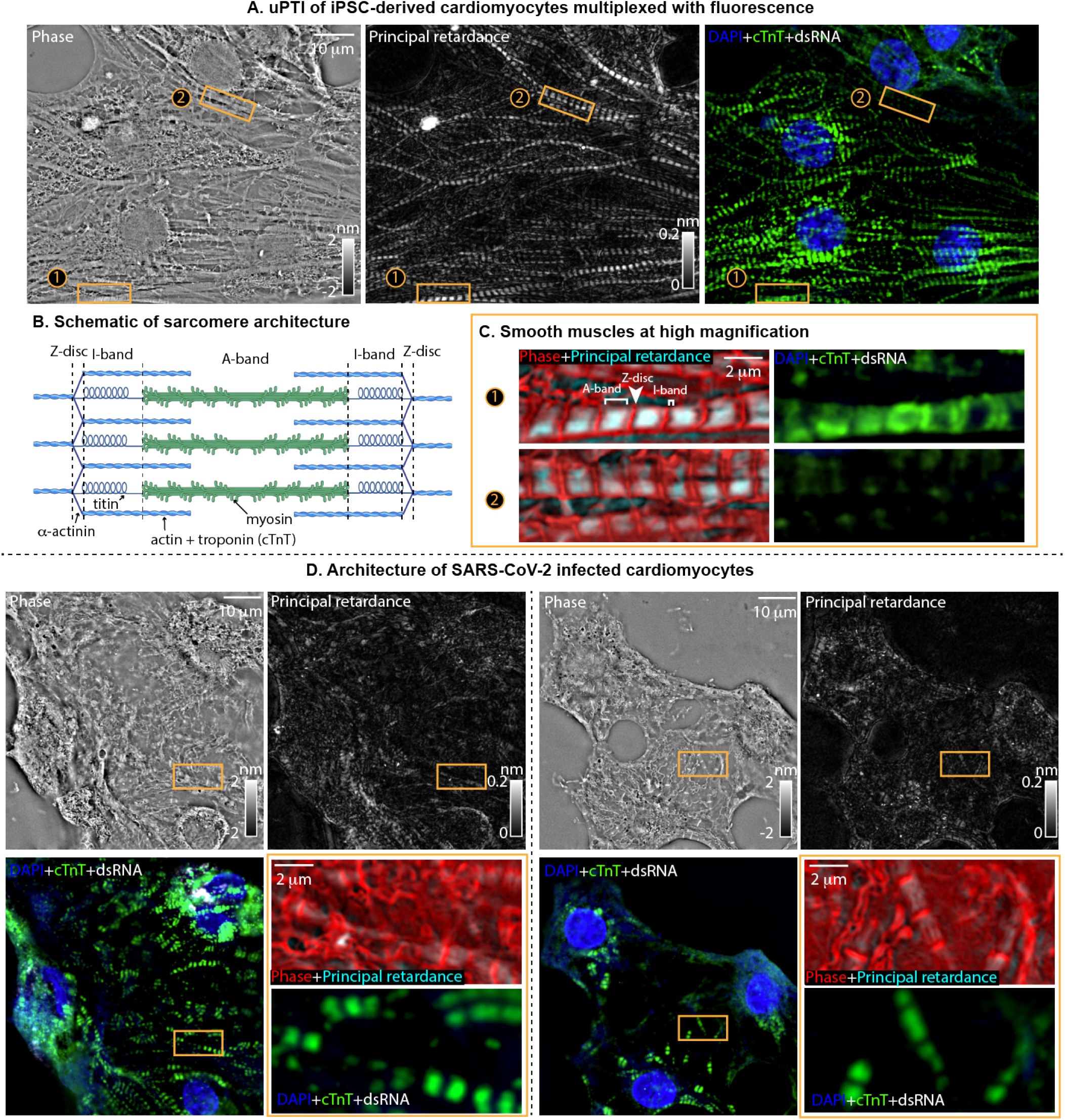
Imaging physical and molecular changes in architecture of iPSC-derived cardiomyocytes due to infection by SARS-CoV-2: **(A)** Phase, principal retardance, and fluorescence images (overlay shows DAPI stain in blue, cTnT stain in green, and dsRNA stain in white) of the uninfected iPSC-derived cardiomyocytes. **(B)** A schematic of the sarcomere architecture (created with BioRender.com) shows its key molecular components and their organization to enable interpretation of the images. **(C)** Two zoomed regions of the iPSC-derived cardiomyocytes are shown with label-free channel (overlay of phase in red and principal retardance in cyan) and the fluorescence channel. Z-disc, A-band, and I-band can be identified in the label-free overlay. A-band and I-band are visible from variations in phase and principal retardance, and Z-disc is visible due to high phase and low retardance. In zoom ①, the cTnT label shows troponin in the actin-rich regions of the sarcomere, overlapping with both I-band and A-band. The zoom ② shows weak fluorescence signal due to labeling stochasticity, but the sarcomeric architecture is visible in the label-free imaging. **(D)** Two fields of view showing the same information as in (A) and (C), but for cardiomyocytes infected with SARS-COV-2. The zooms of both fields of view show broken sarcomeres with label-free overlay and the fluorescence. Relative to mock infection, the phase and fluorescence images show explosion of nuclei and the reduction in the retardance in A-band indicates loss of myosin thick filaments.

Figure 4A and the corresponding through-focus video Video 7 show label-free (phase and principal retardance) and fluorescence images of fixed iPSC-derived CMs. uPT measurements reported in this section are acquired with 1.4 NA (NA_c_) oil immersion condenser and 1.47 NA (NA_o_) oil immersion objective. The phase image shows nuclei, myofibrils, and a crowded meshwork of membranous organelles surrounding sarcomeres. The principal retardance shows the distinct striated pattern of myofibrils. The CMs are stained with DAPI (blue) to label chromatin and fluorescent antibody (green) against thin filament marker cardiac Troponin T (cTnT) to label sarcomeres. We see agreement between the label-free channels and the fluorescence channels in terms of the locations of the nuclei and the sarcomere. However, retardance image shows consistent periodic sarcomere organization that is not always captured by cTnT labeling, especially for ROI-②.

The schematic in fig. 4B shows sarcomere architecture and its key molecular components to enable interpretation of the images. Each sarcomeric unit is bracketed by two Z-discs, composed of densely packed proteins including *α*-actinin. Between Z-discs, the sarcomere is organized in an I-band (isotropic band) and an A-band (anisotropic band). An I-band is mainly composed of thin actin filaments, while an A-band contains thick myosin filaments. The myosin filaments in the A-bands contain bound electrons more easily polarized (higher refractive index) along the filaments, resulting in an angular refractive index distribution of a positive uniaxial material. Figure 4-supplement 1A shows the 3D orientation and optic sign probability of the corresponding field of view. The 3D orientation aligns well along with the orientation of myofibrils. The optic sign probability suggests that thick filaments in the sarcomere behave as a positive uniaxial material, which matches their molecular structure.

We zoom in on two regions of fig. 4A to examine sarcomeric structures in fig. 4C. We display the label-free channels of these two ROIs with an overlay of phase in red and principal retardance in cyan, from which we can clearly resolve sarcomeric components. The phase channel emphasizes the electron-dense Z-disc region. In between Z-discs, we see both strong phase and principal retardance that come from the anisotropic thick myosin filaments, which defines the A-band. We further notice spacing between Z-discs and A-bands. This spacing has lower phase and almost no principal retardance, i.e., it is less dense and isotropic. Comparing its location and size with the transmission electron microscopy images of CMs (55), we identify it as the I-band structure. Figure 4C also shows corresponding fluorescence images for the same ROIs. Since the cTnT is localized in both I-band and A-band, we see most of the signal between two Z-discs in ROI ① of the fluorescence image. cTnT labeling in ROI-② does not detect sarcomeres, while label-free channels clearly detect sarcomeres. This data suggests that label is missing due to inaccessibility of cTnT to antibody or mis-localization of cTnT. Here, label-free imaging complements the inconsistent immunostaining by providing consistent physical measurements of sarcomeres without relying on the labeling of a specific factor.

Recently, iPSC-drived CMs have been shown (55) to recapitulate cytopathic effects of COVID-19 in the autopsy specimens, even though the virus was not detected in the autopsy sections. The most significant phenotypes discovered from these studies are fragmentation of myofibrils and loss of chromatin stain. We multiplex label-free and fluorescence measurements of the SARS-COV-2 infected CMs in fig. 4D and corresponding through-focus videos (Video 8 and Video 9). The infected cells are recognized by immunostaining double stranded RNA (dsRNA), a unique signature of replicating virus. Here we show two distinct fields of view (FOV) of the infected cells from the same coverslip. In the left FOV of fig. 4D, we see substantial reorganization of CMs around the nucleus in both label-free and fluorescence channels. In fluorescence image, double stranded RNA signal is visible in the perinuclear region of this cell, indicating replication of virus through the endoplasmic reticulum-Golgi system. Multiple fragmented myofibrils are visible in our data, especially from cTnT label, as also reported in (55). In both phase and retardance, myofibrils are much less visible, indicating the loss of integrity of sarcomere architecture. In particular, large reduction in retardance suggests loss of thick filaments in A-band. This is in agreement with a recent report that outlines myosin cleavage by a SARS-CoV-2 viral protease (56). Figure 4-supplement 1B shows the 3D orientation and optic sign probability of the corresponding fields of view. The reduced anisotropic signal leads to strong noise in reading 3D orientation and the optic sign prediction, but we can still detect pieces of broken sarcomeres with the parallel orientation and small patches of positive optic sign.

The ability to resolve Z-discs and small I-bands further illustrate high resolution and sensitivity of uPTI, and presents it as a feature-rich tool to evaluate sarcomeric structure, maturity, and the cytopathic effects. These results show that complementary information can be gained in the architecture of cardiac cells and tissues using uPTI multiplexed with fluorescence imaging. Furthermore, these results present uPTI as a powerful tool for the modeling of sarcomeric cardiomyopathies, for screening of cardiotoxic or cardioprotective drugs, or for the development of methods to improve iPS-CM maturity. It also can be applied to other valuable muscle specimens that are challenging to label, such as primary cardiac or skeletal myocytes, or for the non-disruptive imaging of sarcomeric architecture in live muscle cells without the need of engineering fluorescent reporter cell lines. Collectively, these results show that new insights can be gained in the architecture of cardiac cells and tissues using uPTI multiplexed with fluorescence imaging, which opens new opportunities for image-based rich phenotyping of the cellular architecture.

#### Cytopathic effects of RSV infection on A549 cells

Next, we use uPTI multiplexed with fluorescence deconvolution microscopy to analyze cytopathic effects of another model of infection: A549 cells infected by respiratory syncytial virus (RSV). RSV is a leading cause of serious lower respiratory tract infection in infants and elderly (57). Although it is known that RSV replicates in cytoplasm (58), many aspects of its impact on cellular architecture remain to be studied. As a lung cell line, A549 cells are commonly used to study RSV infection (59). Many studies have immunostained A549 cells to investigate the organelle rearrangement (60, 61) during RSV infection. Quantitative measurements of physical properties of RSV-infected cells can clarify the cytopathic effects and enable image-based staging of the infection cycle.

Figure 5A and the corresponding through-focus video Video 10 show images of fixed A549 cells with the label-free channels (phase and principal retardance color-coded with 3D orientation) and the fluorescence channels. uPT measurements reported in this result are acquired with an 1.2 NA (NA_c_) oil immersion condenser and 1.2NA (NA_o_) water immersion objective. The phase image shows nuclei, nucleoli, and a mesh of membranous structures surrounding nuclei. The principal retardance identifies anisotropic fibers around the nucleus and the membranous structures near the the cell-cell junctions. The A549 cells are stained with DAPI (blue) to label DNA and GFP to label RSV. We see co-localization of density, anisotropy, and fluorescence of nuclei. Since these are the uninfected A549 cells, we observe no GFP signal.

**Fig. 5.**
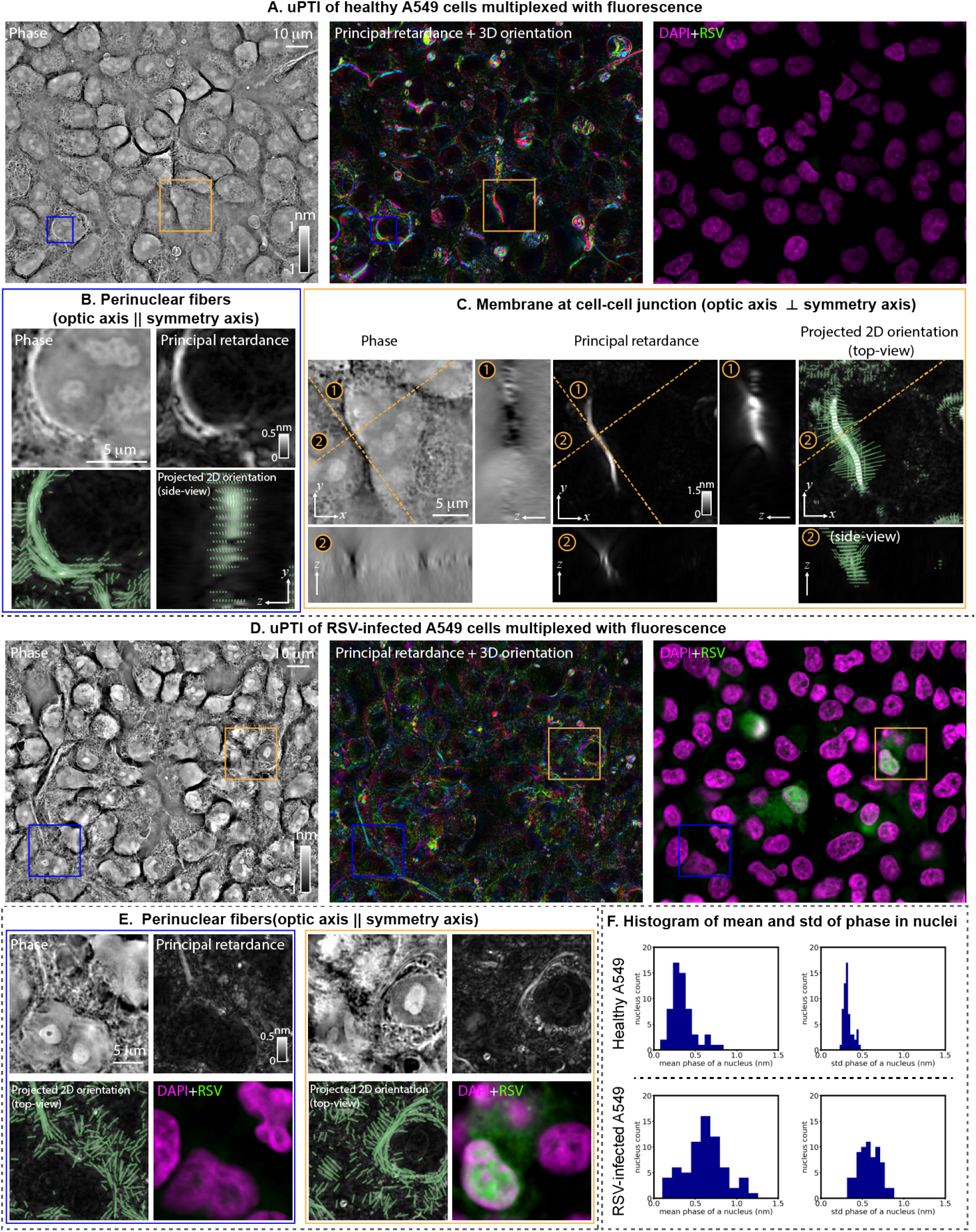
Imaging physical and molecular changes in A549 cells due to infection by respiratory syncytial virus (RSV): **(A)** Phase, principal retardance (color-coded with 3D orientation), and fluorescence images of the uninfected A549 cells. Fluorescence overlay shows DAPI stain in magenta and RSV-GFP signal in green. **(B)** We show an *xy* slice of phase and principal retardance from the volume in blue-square region in (A) to show dense, anisotropic perinuclear structures. We show orthogonal slices (*xy, yz*) of retardance and orientation projected onto the plane of viewing. The projected orientation indicates that the optic axes of these structures are parallel to the their symmetry axes, indicating they are fibers. **(C)** The orthogonal slices of phase, principal retardance, and projected orientation over the field of view indicated by the orange box in (A) show highly anisotropic structures at the cell-cell junction. The orange dashed lines cut at two axes (labeled with ① and ② tangent and normal to the structure to show its cross-sections in phase and principal retardance channels. From the projected 2D orientation, we find the optic axes of this structure is perpendicular to its symmetry axis, indicating these are membranous structures. **(D)** A field of view showing the same information as in (A), but for A549 cells infected with RSV. **(E)** Two fields of view chosen from (D) show the perinuclear fibers observed in (A) and (B) with label-free channels. We do not observe the highly anisotropic membranes at cell-cell junctions in the infected condition. **(F)** Histogram of mean and standard deviation of phase in nuclei for the uninfected and infected A549 cells show significant increase in density and variability in density of nuclei in infected cells. These statistics agree with the variations in phase shown in (B), (C), and (E).

Figure 5B and 5C zoom in on two anisotropic structures indicated by the blue box and the orange box regions in fig. 5A. Figure 5B shows the anisotropic structure with positive phase and 3D orientation of optic axis parallel with to symmetry axis of the structure throughout the volume. The parallel alignment of the optic axis and the symmetry axis of the structure suggests that this is a polymeric fiber assembly with the bound electrons more easily polarized in the fiber axis, analogous to myofibrils in sarcomeres. From the projected 2D orientation in the side view, we see this fiber is slightly inclined out of the focal plane. Figure 5C shows the orthogonal sections of the anisotropic structure at the cell-cell junction along its tangential and normal directions. The orthogonal sections suggest this structure has negative phase and almost 3× stronger principal retardance than the fibers around the nucleus. From the projected 2D orientation in both views, we see the anisotropy is perpendicular to the symmetry axis. This orthogonality between the optic axis and the symmetry axis of the structure suggests this structure is a membrane structure with the bound electrons more polarized perpendicular to the sheet, which is analogous to lipid bi-layers in the myelin sheaths. A549 cells are epithelial cells that form tight junctions between cells when the confluency of cells is high, which can align the membranes of neighboring cells to give rise to high anisotropy. Figure 5-supplement 1A-Figure 5-supplement 1C show the principal retardance and the optic sign probability of this field of view and the zoomed regions of fixed A549 cells, indicating both the perinuclear fibers and the membranes at the cell-cell junctions are positive uniaxial materials.

Now compare above phenotypes with the phenotypes of the RSV infected A549 cells in fig. 5D and corresponding through-focus videos (Video 11). In the fluorescence image, GFP distribution is visible in few cells, indicating they are infected. We observed three consistent phenotypes across multiple cells. First, the highly anisotropic membranes at the cell-cell junctions that are visible in the uninfected cells disappear in the infected cells, suggesting the reorganization of the cellular functions due to RSV infection suppresses formation of these membranes. Second, the perinuclear fibers seen in the uninfected cells are still visible in zoom-in fields of view shown in fig. 5E. Third, the phase or electron density of nuclei and nucleoli in the infected A549 cells are generally larger than the phase of the uninfected A549 cells. Using the DAPI images, we conduct a segmentation of the nuclei for both uninfected and infected A549 cells. Figure 5F shows the histogram of the average and standard deviation of phase value in the nuclei. In the infected condition, we observe noticeably higher average and standard deviation of phase in the nuclei. This suggests that the infection causes the cells to condense themselves. Figure 5-supplement 1D-Figure 5-supplement 1E show the principal retardance and the optic sign probability of this field of view and the zoomed regions of fixed A549 cells. The optic sign probability estimation of the infected cells is noisier than the estimation of the uninfected cells because the infected cells are thicker and may introduce more unwanted scattering artifacts.

Multiplexing uPTI with fluorescence imaging allowed us to identify the RSV infected cells and discover architectural changes due to the infection. Phase, principal retardance, 3D orientation, and the optic sign provide rich information to identify new phenotypes that complement existing immunostaining approaches. We demonstrate that uPTI is useful for disease phenotyping and think it will continue to be useful for studies of multiple infectious diseases.

### E: uPTI reports density and 3D anisotropy of H&E stained histological sections

Microscopic imaging of H&E stained histological sections has been the gold standard in diagnosis of many diseases including many kinds of cancers for a long time (62). Because of their utility, pipelines to generate these sections including proper tissue storage, clearing, staining, and diagnostic tools are well-established. Using uPTI to image already available H&E stained specimens has two immediate advantages: First, the complementary information from uPTI may help discover new phenotypes for better diagnosis of diseases. Second, the compatibility of uPTI with H&E stained sections enables quantitative mapping of the architecture of a wide range of tissues. Here we demonstrate this by multiplexing uPTI with H&E images on two off-the-shelf H&E specimens (Carolina Biological Supply).

For this experiment, we chose 770 nm wavelength for uPTI imaging and illuminated the specimens with red (635 nm), green (525 nm), blue (470 nm) light separately to synthesize H&E images with proper white balance (63) (NA_c_ = NA_o_ = 1.2). The forward model of uPTI assumes weak light-matter interactions in order to simplify recovery of the physical properties of the specimens. In the visible spectrum, H&E stained sections demonstrate strong absorption. We found that imaging at 770 nm avoided the strong absorption and thus avoided any model mismatch.

Figure 6A and the corresponding through-focus videos Video 12-Video 13 show images of mammal cardiac tissue and human uterus tissue (at myometrium) with uPTI and the H&E channels. Similar to fig. 4, in the cardiac tissue, we observe the electron-dense Z-discs of the sarcomeres and nuclei in the phase channel and the anisotropic A-bands of the sarcomeres in the principal retardance channel. Moreover, tissue-scale architecture, unlike cell-scale architecture, shows that bundles of sarcomeres are grouped with the electron-dense cardiac-muscle-specific intercalated discs that show strong phase but low principal retardance (arrows). In the uterus section, we observe electron-dense nuclei and the collagen fibers in the phase channel and, specifically, the anisotropic collagen fibers in the principal retardance channel. Collagen proteins polymerize to form triple helical fibers. The bound electrons of a collagen fiber are more easily polarized along the fiber direction than the radial direction, resulting in an angular refractive index distribution of a positive uniaxial material. All these structures are also visible in the H&E images with blue color referring to the nuclei and the red color referring to protein structures such as sarcomeres and the collagen fibers. Each imaging mode provides complementary views of the tissues with molecular and physical specificity. In addition to phase and the principal retardance, uPTI measure 3D orientation and the optic sign (Figure 6-supplement 1A). These measurements show that both the thick myosin filaments (A-band of the sarcomere) and the collagen fibers are positive uniaxial materials with the optic axes aligned with the long axes of the fibers, which match their molecular structures.

**Fig. 6.**
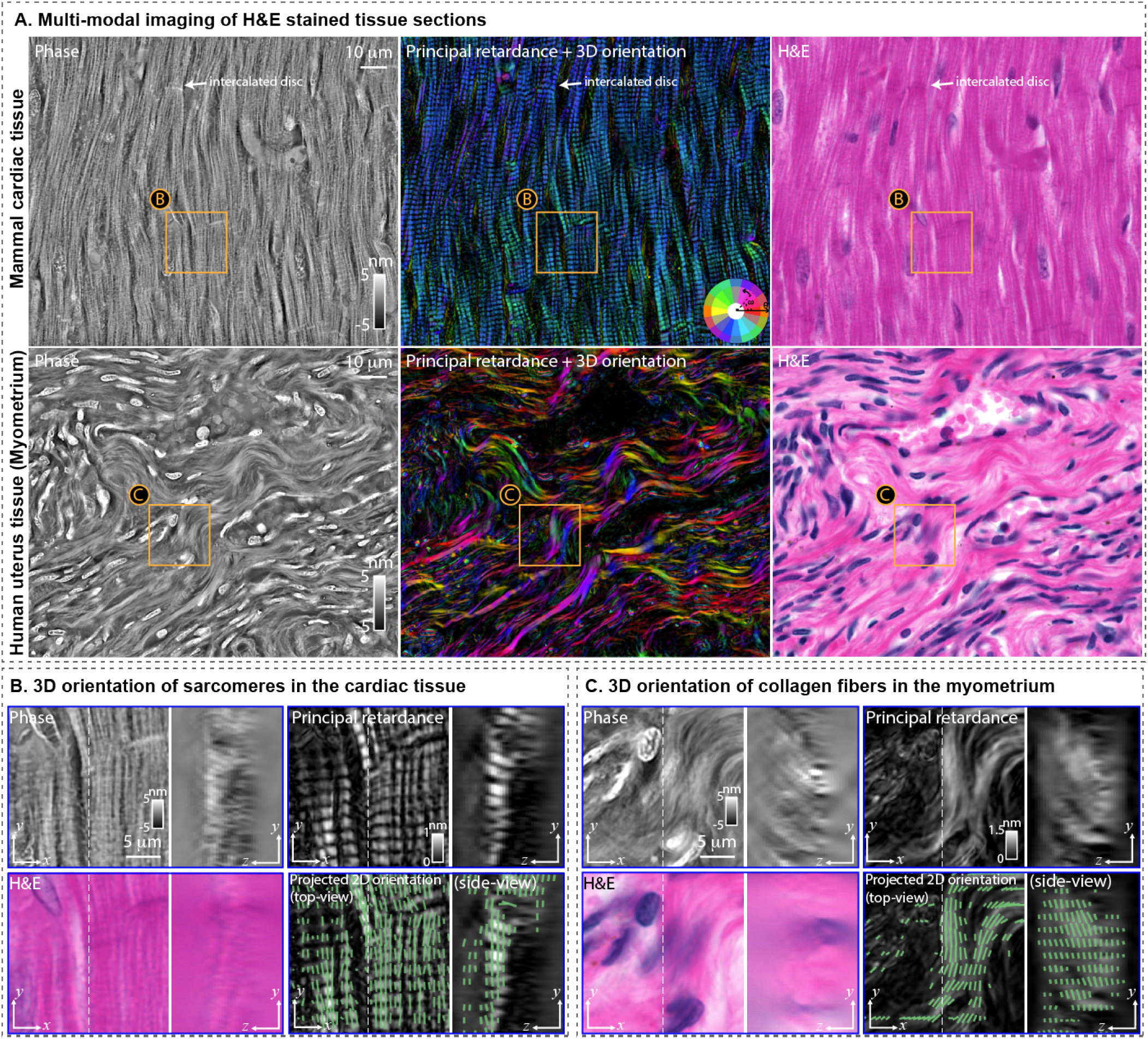
Imaging histological sections with uPTI and H&E stain: **(A)** Phase, principal retardance (color-coded with 3D orientation), and H&E images of (top) the mammal cardiac tissue and (bottom) the myometrium region of the human uterus tissue. We used 770nm wavelength for imaging H&E sections to avoid the strong absorption from the H&E stains. H&E images show histological structures such as nuclei in both tissues, collagen fibers in the uterus tissue, sarcomeres (z-discs, A-bands, and I-bands) and intercalated discs (arrows). All of these structures are visible in phase and principal retardance images at higher contrast and quantitative precision. The principal retardance images specifically highlight anisotropic structures such as A-band of the sarcomere in the cardiac tissue and the collagen fibers in the uterus tissue. The 3D orientation (encoded by the colors in the principal retardance images) clarify how collagen fibers and sarcomeres are arranged. **(B)** The orthogonal slices (*xy* and *yz*) of phase, principal retardance, H&E, and projected 2D orientation of the field of view indicated by the orange box in the cardiac tissue. The 3D orientation of the sarcomeres cannot be observed from the geometry of sarcomeres in H&E or phase images, but is visible with uPTI measurements of principal retardance and projected orientation. **(C)** Same information as in (B) is shown for the field of view indicated by the orange box in the uterus tissue. The 3D orientation of the collagen fibers cannot be observed through the geometry from the *y-z* sections but is measured and shown in the projected 2D orientation channel from uPTI. we employed 780nm illumination for uPTI imaging to limit the absorption from H&E stain.

To better visualize the 3D orientation, we zoom at the orange box regions of the cardiac tissue and the uterus tissue and show the *x-y* and *y-z* sections of phase, principal retardance, H&E, and 3D orientation in fig. 6B and 6C. Here, 3D orientation is projected on the plane of viewing and shown by lines. The *x-z* sections of phase and the principal retardance of both tissues show good sectioning that enables identification of the tissue layers. The principal retardance of fig. 6B shows that the sarcomeres are oriented north-south with small inclination from the focal plane. The 3D orientation visible from the shape of sarcomeres matches well with the two projected views of the 3D orientation measurements from the permittivity tensor. For the uterus section, the principal retardance of fig. 6C does not provide sufficient resolution to visualize 3D orientation purely from the shape of fibers. Fortunately, in this case, the 3D orientation of the permittivity tensor reports the collective orientation of sub-resolution collagen fibers. Figure 6-supplement 1B-Figure 6-supplement 1C show the optic sign probability maps for the same zooms on these tissues. The orientation of collagen fibers is an important prognostic indicator of the human breast cancer (64). We envision this information being potentially useful in cancer diagnosis.

We have demonstrated that uPTI is compatible with the H&E stained tissue sections and provides complementary information regarding the physical properties of the tissues. Researchers have shown that artificial intelligence (AI) can identify rich information from H&E tissue sections for better diagnosis (65, 66). We believe adding uPTI measurements to existing histological tissue sections and AI-platforms will advance the field of general pathology.

## Discussion

We have demonstrated that uPTI provides a more complete description of the uniaxial permittivity tensor of specimens compared to previously reported quantitative phase and polarization microscopy methods. We have illustrated broad utility of uPTI by analyzing the architecture of laser-written anisotropic glass, mouse brain tissue sections, cells infected with respiratory viruses, and H&E-stained histological sections. We also illustrated how the uniaxial permittivity tensor can be interpreted in terms of the physical properties of the specimen and as a function of its 3D orientation. We have implemented automated acquisition and analysis that enable multi-resolution analysis of tissue architecture. Next, we describe how we chose to balance the trade-offs among spatial resolution, temporal resolution, sensitivity, and complexity when designing and implementing uPTI. We also discuss future directions of research enabled by this work.

Volumetric analysis of the architecture of the mouse brain tissue, cardiomyocytes, A549 cells, and H&E tissue sections illustrates that a high-NA implementation of uPTI can measure density and 3D anisotropy with confocal-like resolution, which has been challenging so far. The physical architecture accessible with uPTI is complementary to the molecular architecture that can be imaged with multiplexed fluorescence. The sensitivity and resolution of our data indicate that the measurements provided by uPTI can enable new studies in demyelinating diseases, changes in organelle architectures of infected cells, in mechanobiology, pathology, and other fields. Measurements of 3D orientation and principal retardance at high spatial resolution can provide new quantitative insights in the mechanobiology of polymeric cellular assemblies, such as myofibrils. Our measurements provided one of the first 3D volumes of phase, retardance, and 3D orientation, which have required distinct instruments before. We have employed uPTI to characterize laser-written anisotropic glass, which is a rapidly emerging high density optical storage technology. uPTI can therefore provide a foundation for developing readers of such optical storage devices. While label-free imaging is particularly suitable for live cell imaging, the speed and sensitivity of uPTI need to be improved further to enable live uPTI as discussed next.

uPTI employs simpler hardware relative to the existing label-free microscopy methods that report both density and anisotropy (19–21, 29, 39), yet provides more complete and quantitative measurements. We achieved this simplicity by employing partially coherent illumination and modeling wave optical effects on the phase and polarization of light. The simpler optical design makes it easy to multiplex uPTI with other wide-field imaging modalities, such as fluorescence (as demonstrated), H&E staining (as demonstrated), and spatial transcriptomics. uPTI can be implemented on a commercial microscope by adding an LCD panel, a circular polarizer, and a machine vision polarization camera. This design eliminates the need to tilt or rotate the specimens as required by some of the existing methods. In the present implementation, uPTI is too slow to enable live imaging because of slow refresh rate and slow software communication of the Adafruit LCD. However, faster electronic control of phase and polarization diversity and a more advanced LCD panel (e.g., transmissive spatial light modulator) will enable rapid acquisition that enable imaging of live cells and tissue. uPTI detects polarization-sensitive modulations with a compact polarization camera. Our current choice of using a machine vision polarization camera enabled simple setup and robust calibration, but it is not as sensitive to small changes in retardance as polarization imaging based on elliptical states (26, 28). We are currently extending the design of uPTI to utilize elliptical states, which should lead to higher sensitivity to small changes in anisotropy of biological or fabricated specimens.

Multimodal label-free measurements provided by uPTI experience varying amounts of noise. In particular, the estimations of the optic sign and the inclination of the 3D orientation become unstable when the optic axis is aligned with the imaging axis. This occurs because imaging with a single lens cannot probe the specimen with polarization states that are aligned with the imaging axis. Nevertheless, we show that the high-NA illumination version of uPTI provides sufficient sensitivity to map inclination of axons in mouse brain tissue and enables estimation of optic sign in 2D or 3D space (Figure 3-supplement 1). As far as we know, the estimation of optic sign in 2D or 3D space has never been possible before. Further improvements in robustness of inclination and optic sign can be achieved using elliptical polarization states that reduce noise or using tilted sensors that enable probing of the axial electric field components.

uPTI achieves high transverse and axial resolution using high-NA partially coherent illumination. The partially coherent illumination can provide 2× resolution as compared to approaches that use coherent illumination. Synthetic aperture imaging with coherent illumination (20) can approach the same resolution, but has not yet been extended to measure 3D anisotropy. The partial coherence also makes the measurements robust to speckle noise that commonly affects label-free imaging methods. Relative to the interference-based optical designs (7, 67, 68), our non-interferometric design achieves better robustness to speckle noise by trading-off the sensitivity to large-scale variations in the specimen (23, 31) that correspond to low spatial frequencies.

Spatio-angular measurements of biological systems is a rapidly growing field. Spatio-angular measurements akin to uPTI are being developed with fluorescence polarization imaging (35, 69, 70). We anticipate that the models and deconvolution approaches we have reported will be of value to spatio-angular fluorescence imaging as well. Similarly, modeling and deconvolution of permittivity tensor can be further refined by adopting models and deconvolution algorithms developed for spatio-angular fluorescence imaging and diffusion tensor imaging. Key aspects of our current deconvolution algorithms that we aim to improve: 1) using a calibrated imaging pupil, akin to using calibrated polarization response for accurate imaging in the presence of aberrations, 2) reducing the number of regularization parameters to make the deconvolution algorithm more user-friendly and make it easier to obtain reproducible reconstructions, and 3) extending the model to enable imaging of thicker biological specimens that multiply scatter the light. All of the reported and proposed improvements in image formation and deconvolution are of value in multiple areas of biological microscopy and clinical imaging.

Taken together, we report the unique capability of measuring the uniaxial permittivity tensor of diverse specimens using simple add-on modules on a commercial microscope and open-source diffraction-aware deconvolution algorithm. uPTI has allowed us to image myelination and 3D orientation of axons in mouse brain tissue, the organelle architecture of SARS-CoV-2 infected cardiomyocytes and RSV-infected A549 cells, and 3D anisotropy of the H&E stained tissues. The comprehensive analysis of architecture enabled by uPTI can address open questions of fundamental importance and lead to markers of clinical relevance. Similarly, it can enable quantitative analysis and discovery of new material properties in the material science community.

## Methods

### A: Forward model

In order to deconvolve components of the uniaxial permittivity tensor (uPT) from the intensity measurement, we first develop a forward model that relates the uPT to measured intensities. Our model integrates vector wave equation (71, 72) and partially coherent scalar imaging model (73–75). We summarize the model in this section and provide a detailed derivation in Supplementary Note 1.

We make a key assumption, in addition to the uniaxial symmetry of molecular distribution discussed before, that the recorded intensity modulations are dominated by the interference of the light scattered by a weakly scattering specimen, i.e., the Born approximation (3) is valid. The assumption of weak scattering is typically valid for cells and tissues up to ~ 100 μm thick, which limits the depth of imaging of all single photon imaging methods. We express the distribution of the scattered electric field vector in terms of the distribution of the Stokes parameters, which are linearly related to recorded intensities and, therefore, can be calibrated accurately ((28) and Supplementary Note 2). The assumption of weak scattering leads to a linear relationship between measured Stokes parameters and the unknown components of the scattering potential tensor.

This partially-coherent vectorial model establishes a linear relationship between the measured volumes of Stokes parameters in the image space with the unknown 3D distribution of the uniaxial permittivity tensor in the specimen space, through a set of transfer functions that depend on the size, pattern, and polarization of the illumination and detection apertures of the microscope. This multi-channel transfer function model enables the recovery of specimen properties via multi-channel deconvolution.

#### A.1: Permittivity tensor and scattering potential tensor

The permittivity tensor of an uniaxial material oriented with in-plane orientation of *ω* and inclination of *θ* as shown in fig. 1B is expressed as

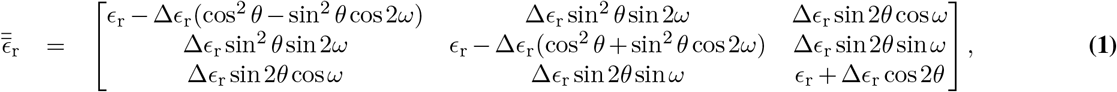

where

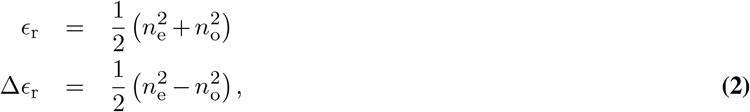

*n_o_* and *n_e_* are refractive indices (RI) experienced by the ordinary and extraordinary wave, respectively. Diffraction tomography approaches have relied on the scattering potential (38) and 2 × 2 scattering potential tensor (29, 39) models to reconstruct volumetric distribution of density and projected anisotropy, respectively. We extend this concept and model 3 × 3 scattering potential tensor to reconstruct volumetric distribution of density, 3D anisotropy, and material symmetry. The scattering potential tensor is defined as,

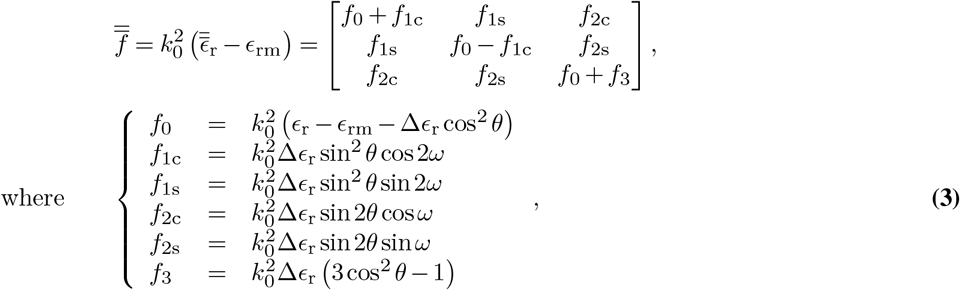

*k*_0_ = 2π/λ_0_ is the free-space wavenumber, *λ*_0_ is the free-space wavelength of the light, and *ϵ*_rm_ is the isotropic relative permittivity of the surrounding media. The scattering potential tensor contains the same information as the permittivity tensor of the specimen, except that it is relative to the permittivity of the surrounding medium.

#### A.2: Scattered electric field due to the scattering potential tensor

The Stokes parameters are measurements of the scattered electric field. The scattered electric field comes from the interaction between the incident light and the scattering potential tensor, and it is derived based on the vector wave equation (71, 72) and Born approximation (3) as

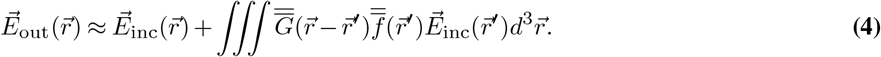

where 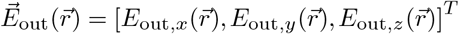 is the scattered output electric field in 3D space 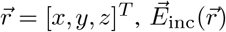 is the incident electric field, and 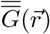 is the dyadic Green’s tensor. This equation describes a single scattering event from a coherent incident wave.

#### A.3: Stokes parameters under partially coherent illumination

In our experiments, we use partially coherent illumination from large illumination NA, which causes the recorded intensities to be the sum of intensities due to coherent scattering of light at each angle of illumination.

Each angle of illumination modulates the specimen with electric field of spatial frequency 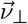. We first define the scattered electric field with incident light of spatial frequency 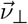 to be 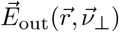. Then, the Stokes parameters under the *α*-th partially coherent source pattern are sums of the contribution from individual coherent scattering events and are expressed as in (73, 74)

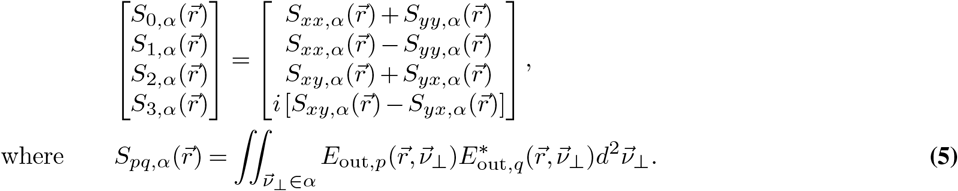

From here, we neglect the nonlinear contribution to the Stokes parameters from the scattering potential tensor, which is usually small for weakly scattering specimens, and arrive at our linearized forward model expressed in the Fourier space as

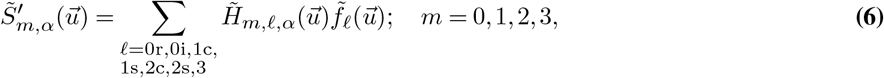

where 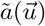 denotes the Fourier transform of a function 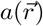 at the 3D spatial frequency 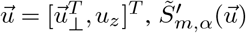 is the DC-subtracted Stokes parameter, 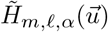 is the transfer function mapping from each scattering potential tensor component *ℓ* to the *m*-th Stokes parameter under illumination pattern *α*. In a 2D imaging case, the forward model only requires a simple modification of *u_z_*-integration and is expressed as

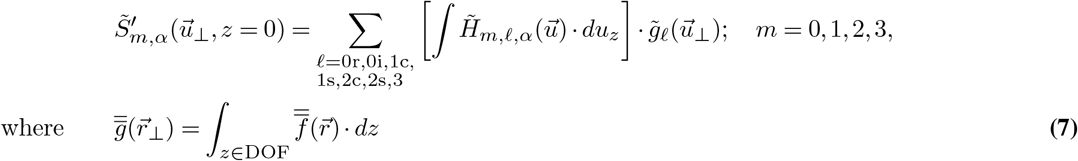

### B: Deconvolution algorithm

Our deconvolution algorithm takes the Stokes parameters of the scattered light under different illuminations (converted from measured intensity images through calibration and background correction, Supplementary Note 2) as inputs to estimate the physical properties that are encoded in the uPT. Since some parts of this algorithm are nonlinear, we decompose this algorithm into three parts to achieve robust estimation and computational efficiency. The first part of the algorithm is a least square optimization solver that estimates the entries of the scattering potential tensor in 3D space. Second, we compute the phase, principal retardance, and 3D orientation from the entries of the scattering potential tensor assuming that each voxel is a positive and a negative uniaxial material. The last part of the algorithm fits these two solutions to the recorded Stokes volumes via the forward model to estimate the optic sign in 3D.

#### B.1: Solving for scattering potential tensor

According to the forward model Eq. (6), the DC-subtracted (background subtracted) Stokes parameters of the scattered light, 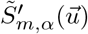, under different illuminations are linearly related to the entries of the scattering potential tensors. Thus, we can write down an inverse problem to retrieve the entries of the scattering potential tensor as

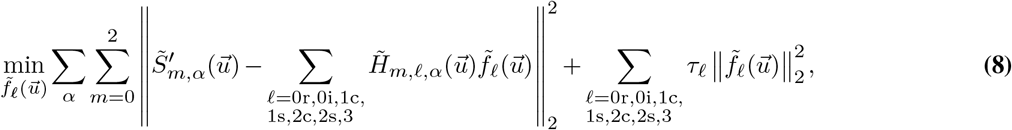

where *τ_ℓ_* is the Tikhonov regularization parameter for each independent term in the scattering potential tensor. In our implementation, this optimization is done in the Fourier space, where each voxel (or pixel) in Fourier space is independent from other voxels (or pixels).

We speed up computation of Eq. (8) by solving the point-wise optimization problems at all spatial frequencies 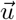 in parallel. The least square solution of this one-point optimization can be obtained by solving the fully determined inverse problem of

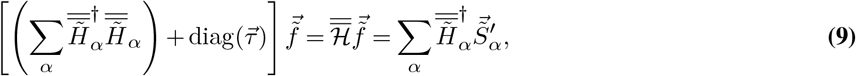

where

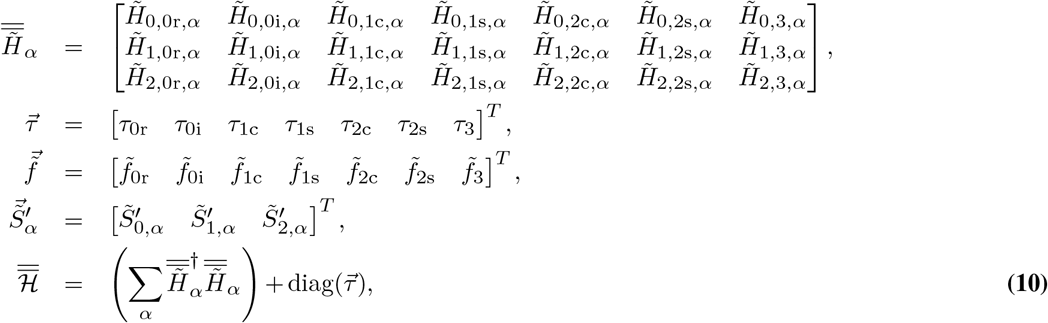

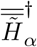 denotes the Hermitian of the matrix 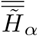. With this fully determined inverse problem, we apply the Cramer’s rule on it to obtain the analytical solution as

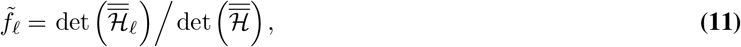

where 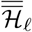 denotes that the *ℓ*-th column of the 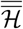 matrix is replaced with 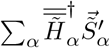.

We set the regularization *τ_ℓ_* as a multiple of the mean absolute value of the *ℓ*-th diagonal entry of 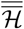 over all spatial frequency 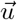 to reduce its dependency on different imaging parameters. Usually the multiple factor of *τ_ℓ_* is ranging from 1 to 100 (smaller for less noisy data to get sharper reconstruction and larger for more noisy data to smooth out the noise). It should also be noted that, for simplicity of the math, we only show the single point solution of this deconvolution problem. In the actual implementation, we vectorize the whole 3D array of the data and the transfer functions for fast parallel processing with GPU.

#### B.2: Phase, principal retardance and 3D orientation from scattering potential tensor

After solving Eq. (8), we obtain components of scattering potential tensor as shown in Eq. (3). 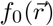 is related to the difference of average permittivity between the specimen and the environment, which corresponds to accumulated optical phase and absorption information. If the effect of phase and absorption is much stronger than the effect of optical anisotropy, 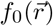 is roughly proportional to the phase and the absorption. If the permittivity of the environment is close to the average permittivity of the specimen, the phase and absorption are approximately related to 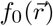 with the following scaling

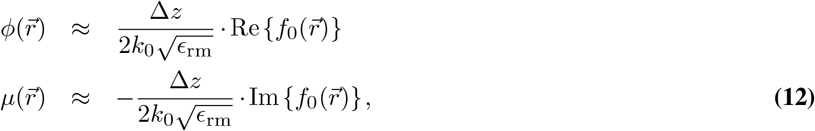

where 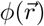 and 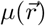 are phase and absorption of the specimen and Δ*z* is the sampling size of our data in the axial direction.

Estimating principal retardance and 3D orientation from four terms in the scattering potential tensor, 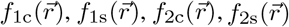, requires information of the optic sign (*n_e_* > *n_o_* or *n_e_* < *n_o_*). Here, we first compute two sets of analytical solutions for principal retardance and 3D orientation assuming two optic signs. These two sets of solutions will be used to estimate the probability of the optic signs in the next part of algorithm. Using trigonometry relations, we express these two solutions of principal retardance and 3D orientation as

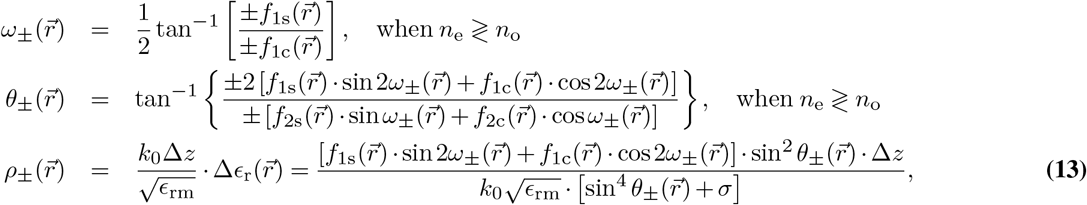

where *ω*_±_, *θ*_±_, and *ρ*_±_ are in-plane orientation, out-of-plane inclination and principal retardance of the positive or negative uniaxial material. *σ* is a small number to prevent noise amplification in the estimation of principal retardance. Typically, *σ* = 10^-2^ ~ 10^-3^ is a good choice to balance between accuracy and noise amplification.

At the end of this computation, the range of the azimuthal angle is *ω* ∈ [0,*π*) and of the inclination is *θ* ∈ [0,*π*). These ranges correspond to the front half of the hemisphere of unit sphere (*y* ≥ 0). For intuitive visualization of 3D orientation, we transform (*ω, θ*) coordinates to span the range of the top hemisphere (fig. 1D, *z* ≥ 0) by reflecting the 3D orientations around the origin.

#### B.3: Optic sign from scattering potential tensor

The optic sign of the anisotropy reports symmetry of underlying structure. If we know of the type of the material imaged, we pick one set of solution from *ω*_±_, *θ*_±_, and *ρ*_±_. More often, the optic sign of a biological structure is not known and can be spatially variable.

For cases where the optic sign is unknown, we employ an algorithm that models the scattering potential as a mixture of positive uniaxial and negative uniaxial material. The algorithm starts with constructing the scattering potential tensor components with positive (*ω*_+_, *θ*_+_, *ρ*_+_) and negative (*ω*_, *θ*_, *ρ*_) set of solutions according to Eq. (3). We then want to know which set of solution is more favorable in our data by solving the following optimization problem

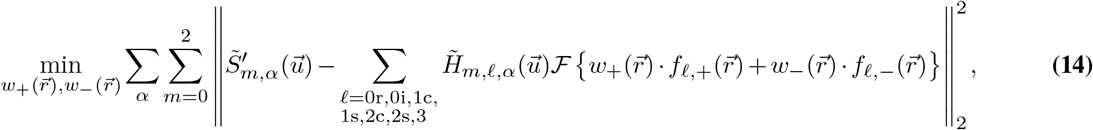

where 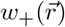 and 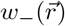 are the weights for positive and negative uniaxial solutions (we only consider positive values of the weight). When the positive material is more favorable (the structure within a voxel is denser along axis of symmetry), 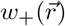 is larger than 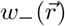. On the other hand, 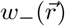 is larger than 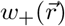 when negative material is more favorable (the structure is denser perpendicular to the symmetry axis). When the material is isotropic, 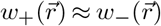. We implement a gradient descent iterative algorithm to solve this optimization. To identify material type with these two weights, we define the probability of being positive material to be

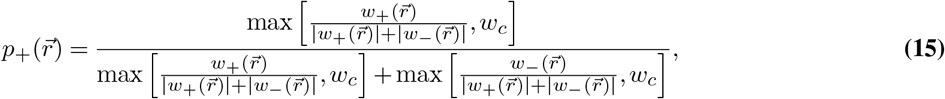

where *w_c_* is a cut-off weight to threshold out noisy weight estimate for smooth probability reconstruction. *w_c_* is usually set around 0.05 ~ 0.2. The higher the value, the stronger the thresholding effect.

### C: Multi-scale analysis of optical properties by averaging the permittivity tensor

In fig. 3, we describe measurements of optical properties of the mouse brain tissue at multiple scales. Computing physically meaningful measurements of density, retardance, 3D orientation, and optic sign at lower spatial scales from the high-resolution acquisition requires a specific filtering approach. The reconstruction steps that lead to components of the permittivity tensor (Eq. (11)) are linear. But, the computation of optical properties from the permittivity tensor components involves non-linear operations (Eq. (13) and Eq. (15)). Therefore, the low resolution measurements of optical properties are computed by linear filtering of the high resolution volumes of the scattering potential tensor, which are transformed into phase, retardance, 3D orientation, and optic sign according to Eq. (13) and Eq. (15). This approach ensures that the computed optical properties at lower resolution are representative of an acquisition with a lightpath of lower spatial resolution.

### D: Improving the sensitivity of uPTI through averaging and denoising

uPTI converts the intensity modulation from the density and anisotropy of the specimens into physical properties as shown in fig. 1C and 1D. These intensity modulations are built upon a constant intensity background that defines the transmission of the surrounding media. The ratio between the strength of the intensity modulation and the shot noise created by the constant background intensity defines the signal to noise ratio (SNR) of our measurements. Biological specimens that we work with generally have strong phase (average refractive index) and relatively weak anisotropy. Hence, SNR is typically lower for anisotropy measurements. In the following, we take two approaches to improve the SNR of our measurements.

The first approach is to reduce the shot noise in the raw intensity through averaging multiple frames. Since we cannot change the strength of anisotropy from a biological specimen, we average our raw intensity by *N_avg_* frames to improve our SNR by 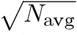 times. *N*_avg_ varies by experiments. For strongly anisotropic structures such as the anisotropic glass target in fig. 2 and myelin sheaths in fig. 3 (principal retardance > 2 nm with the high-NA objective), *N*_avg_ = 1 already gives satisfying results. For weaker anisotropic structures such as iPSC-derived cardiomyocytes in fig. 4 and A549 cells in fig. 5, we average 32 frames and 100 frames per intensity acquisition, respectively, to increase the SNR.

The second approach is to apply an additional wavelet denoising on the principal retardance after the deconvolution algorithm Eq. (8). Tikhonov regularizer in the existing deconvolution algorithm prevents over-amplification of noise in the reconstructed physical properties. Total-variation or wavelet-based denoising algorithms that leverage the continuity of the images could provide additional improvements in the SNR of the image. However, a proper implementation of these algorithms with the deconvolution requires an iterative approach to solve the optimization problem, further increasing the currently heavy computation load. We then adopt a compromised solution, directly performing wavelet denoising algorithm on the principal retardance images, to balance the need of SNR improvement and computation load. The denoising algorithm is a single-step soft-thresholding operation in the wavelet space on the images. This operation produces a solution of the following the optimization formalism (76)

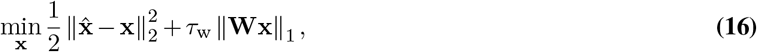

where 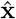 is the input image, *τ*_w_ is the wavelet regularization parameter, and **W** is the operator of the wavelet transform. The solution of this optimization problem is analytical and is expressed as

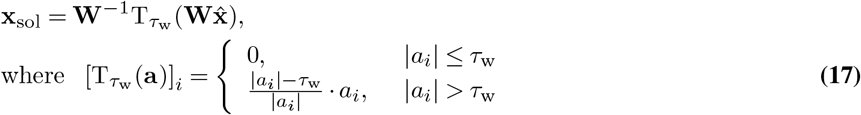

To denoise the principal retardance images, *τ*_w_ is generally chosen to be 10% to 20% of the average signal level.

In addition to low SNR of the principal retardance measurements, strong phase discontinuities can mask the true anisotropy by introducing edge retardance. A distinguishing feature of the edge retardance is shows two maxima (42) around the phase edge with orthogonal orientations. The retardance of a true anisotropic structure (not the edge retardance) shows single maxima of a consistent orientation. To emphasize true anisotropy in the principal retardance channel, we suppress edge retardance by computing orientation consistency map. Using this map, we reject features with fast-varying orientation. We apply this additional filtering on the cardiomyocyte dataset we show in fig. 4.

The orientation continuity map is computed through the following steps. First, we synthesize the scattering potential tensor components with constant retardance and filter these synthesized tensor components with a uniform filters, 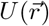, of kernel size *N_k_* to have the averaged scattering potential tensor components as

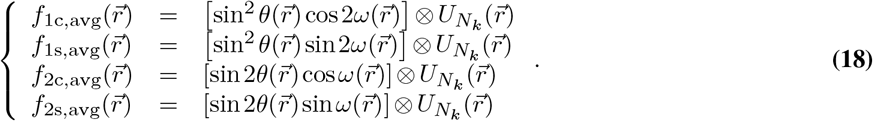

Second, we use these averaged scattering potential tensor components to compute the principal retardance according to Eq. (13). The averaged principal retardance shows high values when the orientation is more continuous along spatial dimensions and small values when the orientation is varying pixel by pixel. The orientation continuity map is derived from the average retardance normalized by its maximum. Multiplying this map with the original principal retardance from the measurements, we eliminate signals from the edge retardance and see the low-retardance structures better. This method is effective when the orientation of the structures is more continuous in space (fig. 4). Potential artifacts may happen when the orientation of the structures varies more strongly in space (lipids in fig. 3C).

### E: Automation for multi-scale imaging and analysis

We automate the multi-scale imaging shown in fig. 3 and Video 4 by controlling individual devices in python. This acquisition requires control of three main devices, the LCD panel for switching illumination patterns, the machine vision polarization camera for collecting images, and the microscope stages for scanning in *x*, *y,* and *z* directions. First, we control the LCD panel using the built-in APIs from Adafruit with Arduino board. Serial connection is established from the acquisition computer to the Arduino board for software triggering in python. Second, we control the polarization camera with a python package, PySpin, developed by camera manufacturer, FLIR. Last, the microscope stage is controlled by Micro-Manager (https://github.com/micro-manager). In order to build a bridge between the Java-based Micro-Manager and python, we leverage the mm2python library (https://github.com/czbiohub/mm2python). Collectively, these packages allow us to compose an acquisition script to control each device. For the 2D acquisition shown in fig. 3A-3B, we acquired 9 images under different illumination patterns per location for total of 609 FOVs in a 29 × 21 (*x* × *y*) rectangular grid. The overlap between each location is set to be ~ 15% in *x* direction and ~ 30% in *y* direction. For 3D acquistion shown in fig. 3C-3I and Video 4, we acquired 9 × 120 (pattern × *z*) images to form a *z*-stack per location for a total 153 FOVs in a 17 × 9 (*x* × *y*) rectangular grid. The overlap parameter is similar to the previous case.

We implement our algorithm to be GPU-compatible on an IBM Power9 server equipped with 4 GPUs (Tesla V100-SXM2-32GB, NVIDIA) per compute node. One FOV of 2D acquisition easily fits in the memory of one GPU, so we initiate 4 instances of computation with 4 different GPU to process the data. The algorithm takes about 200 seconds to process one FOV of 2D acquisition and about half an hour to stitch one channel of the reconstruction. Fitting a FOV of 3D acquisition in the memory of one GPU is infeasible, so we break one 3D acquisition into 30 (or smaller number) small patches for processing. We also initiate 4 to 8 instances of computation with 4 GPUs to process these small patches in parallel. Each FOV of 3D acquisition takes about 6 hours of the processing time and the stitching process takes about 3 hours for one channel of the reconstruction.

### F: Computation of 3D orientations of axons and their cross-sections from structure tensor of phase image

To obtain the structure tensor for analysis of axons’ morphological orientation, we consult and modify the vessel segmentation algorithm from the X-ray angiography community (77) for our data shown in fig. 3D. The algorithm first computes the structure tensor, the second-order gradient, of the phase image 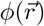, as

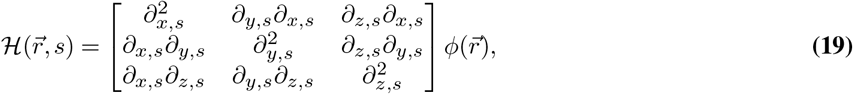

where *s* is the scale parameter and scale-dependent gradient is defined as a convolution between the scaled intensity and the derivative of a Gaussian function written as

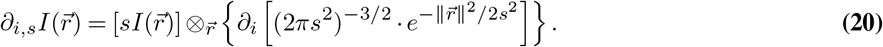

Depending on the geometry of the interested feature (a shell-like, tubular-like, or blob-like structures), the eigenvalues of the structure tensor may be used in different ways to construct the segmentation map. In our case, we selected shell-like structure and construct the segmentation map as

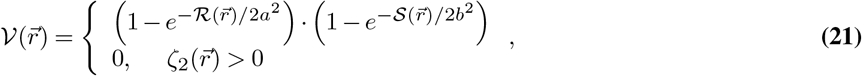

where

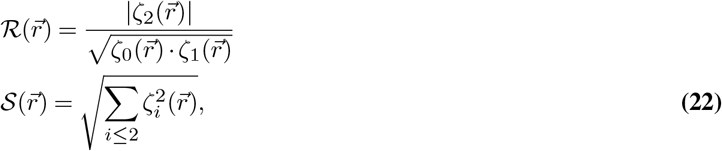

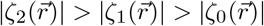 are eigenvalues of the structure tensor at position 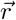, *a* and *b* are parameters to adjust the level of sharpness of the segmentation map. The eigenvectors corresponding to the largest eigenvalue of the structure tensor on every voxel within this map contain the normal orientation of the shell-like structures. We then use these eigenvectors as a different measurement of lipids’ 3D orientation to verify our results from 3D anisotropy.

### G: Specimen preparation

#### Femtosecond laser written anisotropic glass

The target used in fig. 2 was written into a fused silica cover glass that is about 0.25 mm thick and 22 mm by 22 mm on its sides using a polarized femtosecond laser. The star pattern consists of 32 equally spaced birefringent wedges that rotate in steps of 11.25°. The wedges consist of a single line near the center of the star, flanked by additional lines towards larger diameters (1 line between 3 and 20 μm diameter, 3 lines between 20 and 40 μm, and 5 lines between 40 and 60 μm). While the slow axis of a wedge rotated with the wedge, within a wedge, the slow axis was uniform and was parallel to the lines. Each line was written only once and the scanning direction was parallel to the slow axis. The parameters of laser fabrication were the following: pulse duration – 500 fs, repetition rate – 500 kHz, fabrication speed – 0.01 mm/s, wavelength – 515 nm, focused with 0.55 NA lens. More details on the fused silica modification through laser writing are documented in Supplementary Note 4.

#### Mouse brain section

The mice were anesthetized by inhalation of isoflurane in a chemical fume hood and then perfused with 25 ml phosphate-buffered saline (PBS) into the left cardiac ventricle and subsequently with 25 ml of 4% paraformaldehyde (PFA) in the PBS solution. Thereafter, the brains were post-fixed with 4% PFA for 12-16 hours and then transferred to 30% sucrose solution at the temperature of 4°C for 2-3 days until the tissue sank to the bottom of the container. Then, the brains were embedded in a tissue freezing medium (Tissue-Tek O.C.T compound 4583, Sakura) and kept at the temperature of –80°C. Cryostat-microtome (Leica CM 1850, Huston TX) was used for preparing the tissue sections (12 and 50 *μ*m) at the temperature of –20°C and the slides were stored at the temperature of −20°C until use. Upon experiment, the OCT on the slides were melted by keeping the slides at 37°C for 15-30 minutes. Then, the slides were washed in PBST (PBS+Tween-20 [0.1%]) for five minutes and then washed in PBS for five minutes and coversliped by mounting media (F4680, FluromountTM aqueousm sigma).

#### iPSC cardiomyocyte

Cardiomyocytes were differentiated from iPS cells (WTc cell line (78)) using a modified Wnt pathway modulation protocol (79). Briefly, cells were maintained in mTesr media (Stem Cell Technologies) and three days before differentiation, they were seeded on 12-well plates. During differentiation, basal media was RPMI supplemented with B-27 minus insulin (Gibco) for days 0-7 and RPMI with B-27 (Gibco) on days 7-onwards. Cells were treated with 6 μM CHIR99021 (Tocris) for 48 hours on day 0, and with 5 μM IWP2 (Tocris) for 48 hours on day 3. At day 15, cells were harvested and stored on cryovials. When ready for experiments, cell pools were thawed in RPMI with B-27 supplemented with 20% FBS (HyClone) and ROCK inhibitor Y-27632 (10 μM, Selleckchem). Cardiomyocytes were then selected in culture using a metabolic switch method (80) by treating the cells with 4 mM lactate media changes every other day for 6 days. Final cultures were >90% ACTN+.

On day 30, cardiomyocytes were replated into glass coverslips, and maintained on RPMI with B-27 for five more days. Then they were fixed using 4% paraformaldehyde for 20 minutes at room temperature, washed three times with PBS supplemented with Triton X-100 (PBS-T) and blocked and permeabilized with 5% bovine serum albumin in PBS-T. Cells were then stained with an anti-cTnT antibody (Abcam, ab45932) in PBS-T overnight, and with DAPI for 10 minutes. After three PBS-T washes, Alexa Fluor 488 anti-rabbit (ThermoFisher) in PBS-T was used as secondary antibody, followed by three more PBS-T washes. Then, a drop of Prolong Antifade (without DAPI) (ThermoFisher) was added to coverslips for mounting into a glass slide.

## Acknowledgements

We thank Rafael Gómez-Sjöberg (CZ Biohub) and Mirella Bucci (CZ Biohub) for critical reading of the manuscript; Andreas Puschnik (CZ Biohub), Joseph DeRisi (CZ Biohub/UCSF), and Sara Sunshine (UCSF) for providing A549 cell line and RSV. L-H.Y, I.E.I, B.C, S-M.G, J.R.B, and S.B.M are supported by the intramural program of the Chan Zuckerberg Biohub. B.R.C received support from the Gladstone Institutes, Innovative Genomics Institute and NIH grants R01-HL130533, R01-HL13535801, P01-HL146366. This research is supported by the Chan Zuckerberg Biohub.

## Supplementary Figures

**Figure 1-supplement 1.**
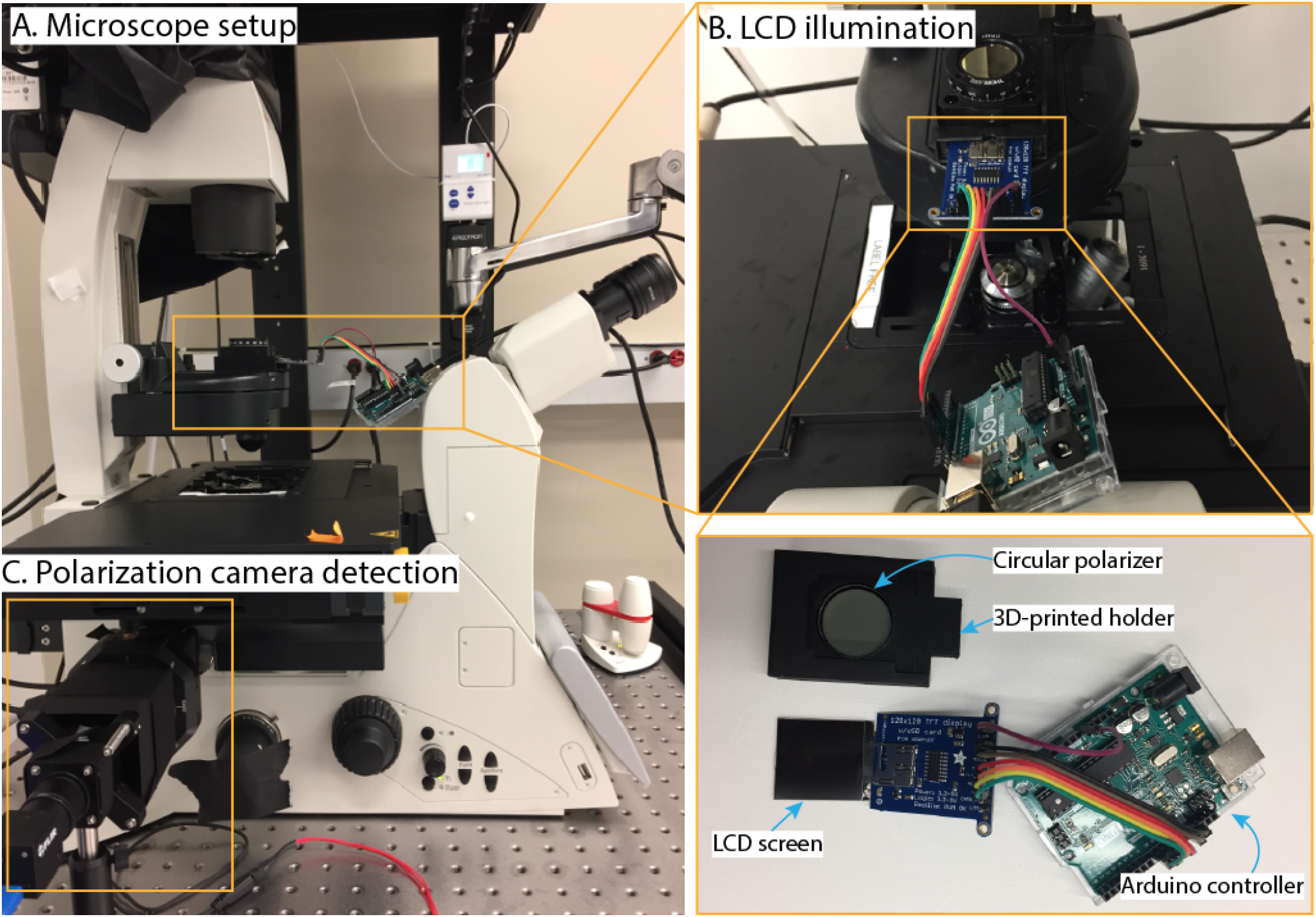
Photographs of the uPTI setup: **(A)** The microscope setup. **(B)** The add-on LCD illumination setup composed of adafruit LCD screen (Adafruit, ST7735R) and the circular polarizer (Thorlabs, CP1R532). **(C)** The add-on polarization camera (FLIR, BFS-U3-51S5P-C) for detection.

**Figure 1-supplement 2.**
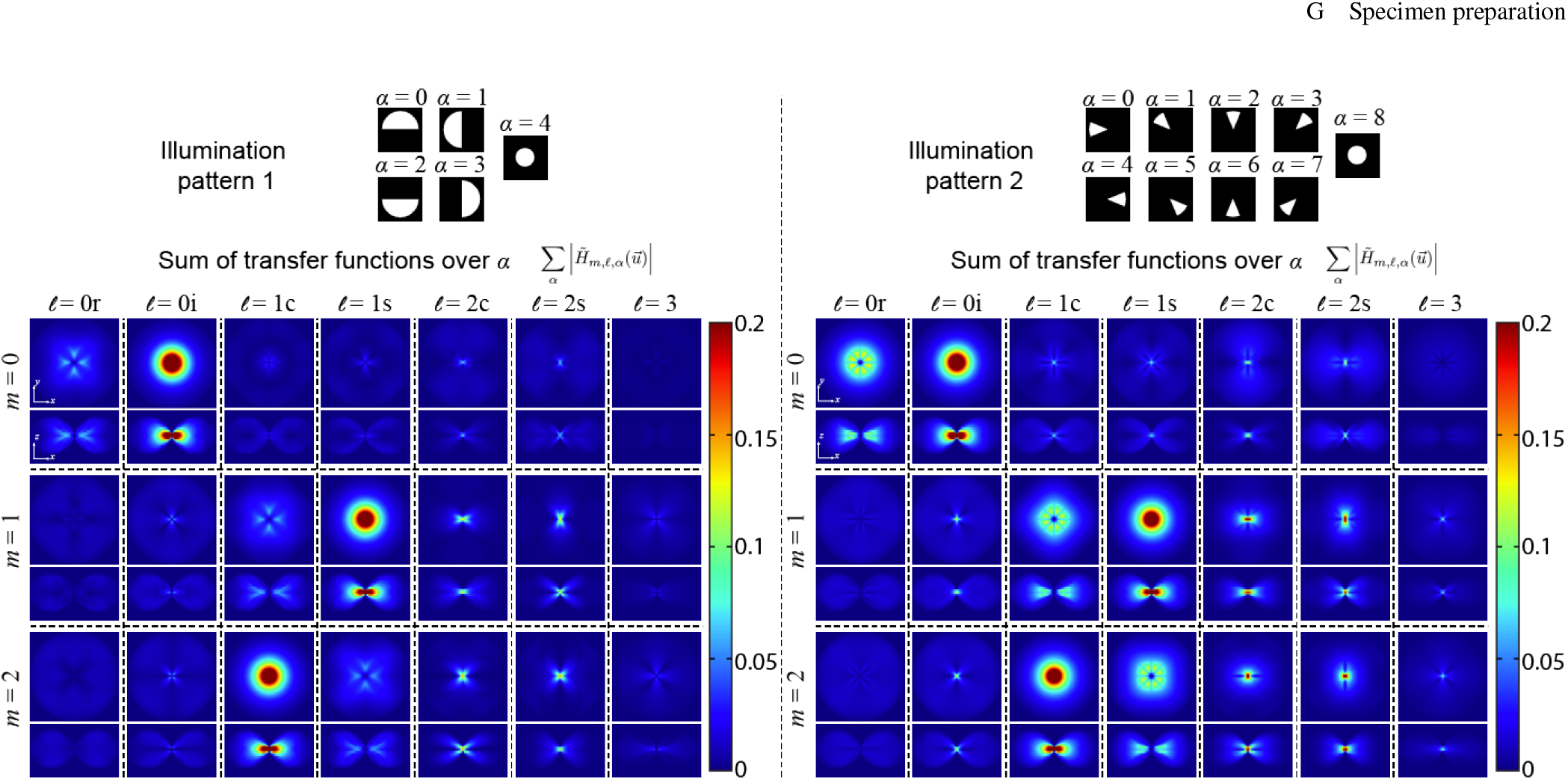
Comparison of transfer functions for two sets of illumination patterns: Sum of transfer functions (mapping from the *ℓ*-th individual scattering potential tensor component to the *m*-th Stokes parameres) over all illumination patterns (*α*) with the combination of semi-circular (typically used for differential phase contrast) and brightfield illumination (left) and the combination of sector and brightfield illumination (right). We adopt the right set of illumination pattern, which transfers more uniformly across 3D spatial frequencies in all tensor components.

**Figure 1-supplement 3.**
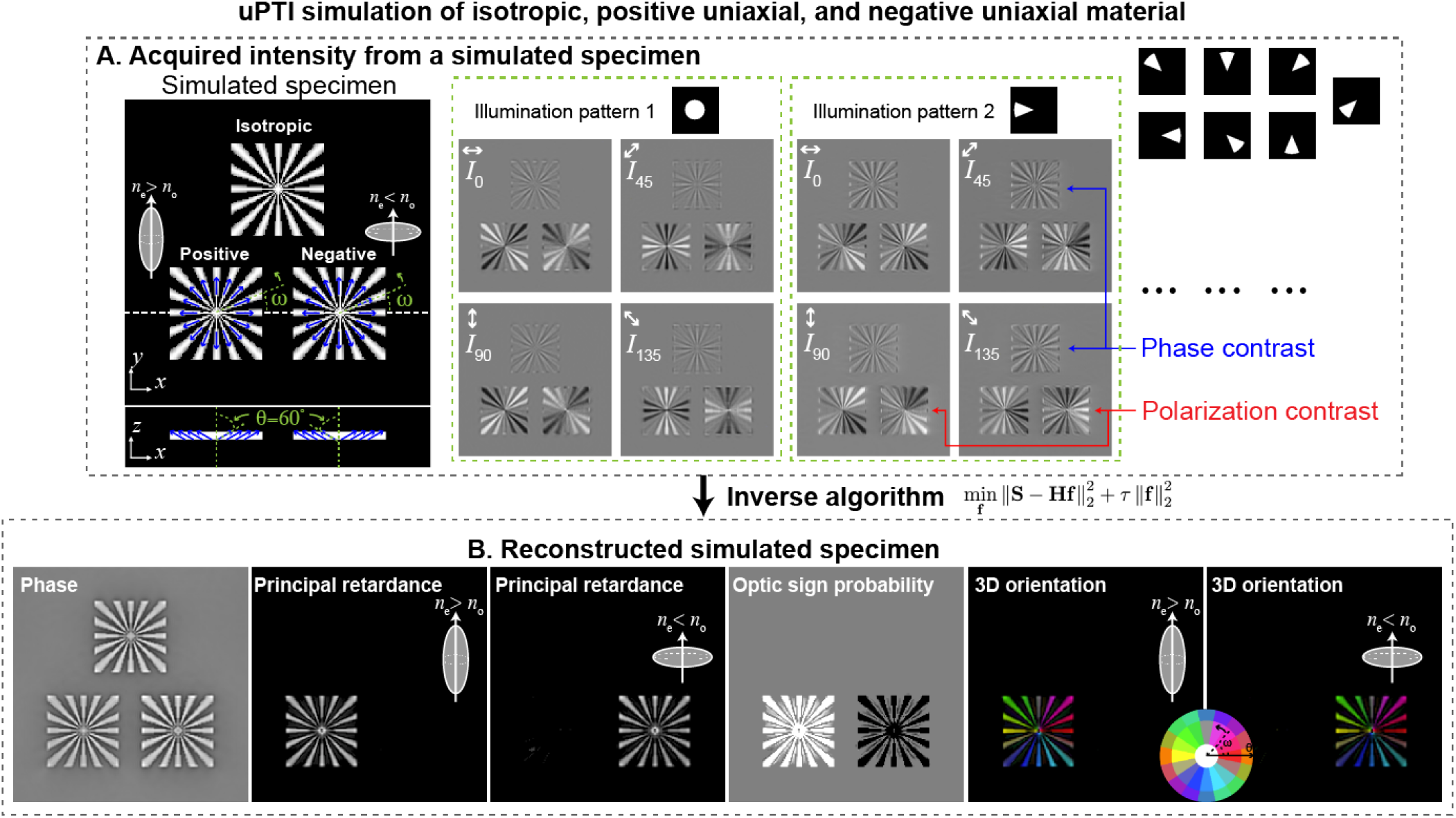
uPTI simulation of isotropic, positive uniaxial, and negative uniaxial test targets: **(A)** Simulated raw images of a structure consisting of isotropic material (*n_e_* = *n_o_* = 1.52), positive uniaxial material (*n_e_* = 1.521, *n_o_* = 1.519), and negative uniaxial material (*n_e_* = 1.519, *n_o_* = 1.521) oriented radially (along the spokes) and inclined with *θ* = 60° illustrate dependence of intensity contrast on the polarization and the angle of illumination. The refractive index of the surrounding medium is set to 1.518. Each type of material is 5.7 × 5.7 μm^2^ in size. We set the NA of illumination to be 1.4 and the NA of the objective to be 1.47 to mimic the experimental conditions. We observe intensity modulations due to specimen properties in the simulated intensities. Phase variations are evident in isotropic material with off-axis illumination (illumination pattern 2), while anisotropy variations are better observed in different polarization channels. We also observe contrast variations across on-axis and off-axis illumination due to variations in out-of-plane orientation and optic sign. **(B)** Using an inverse algorithm based on convex optimization, we reconstruct phase, principal retardance, optic sign probability, and 3D orientation of the simulated target from intensities.

**Figure 2-supplement 1.**
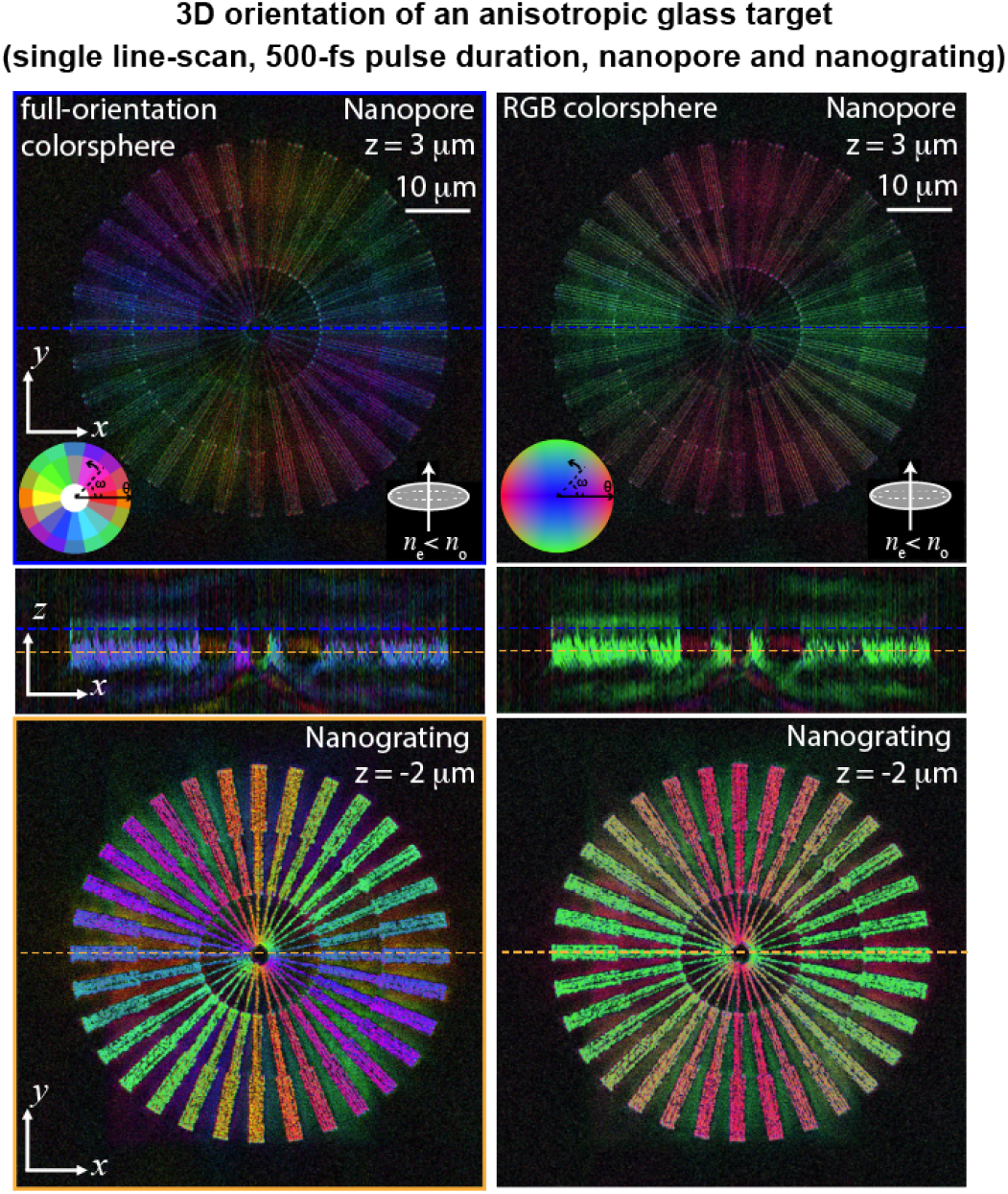
3D orientation of the anisotropic glass target: The 3D orientation of the anisotropic glass target shown in fig. 2C is rendered with the full-orientation colored sphere (left) and the RGB colored sphere (right). The full-orientation colored sphere encodes full range (upper hemisphere) of measurable 3D orientation, but is more complex to read. The RGB colored sphere maps the absolute *x, y,* and *z* components of the 3D orientation into red, green, and blue color, which is more intuitive to read, but maps only one octant of the 3D orientation, i.e., quarter of the measurable range. From both renderings, we see the 3D orientation of the target is perpendicular to the spoke feature and mostly in the *x-y* plane.

**Figure 2-supplement 2.**
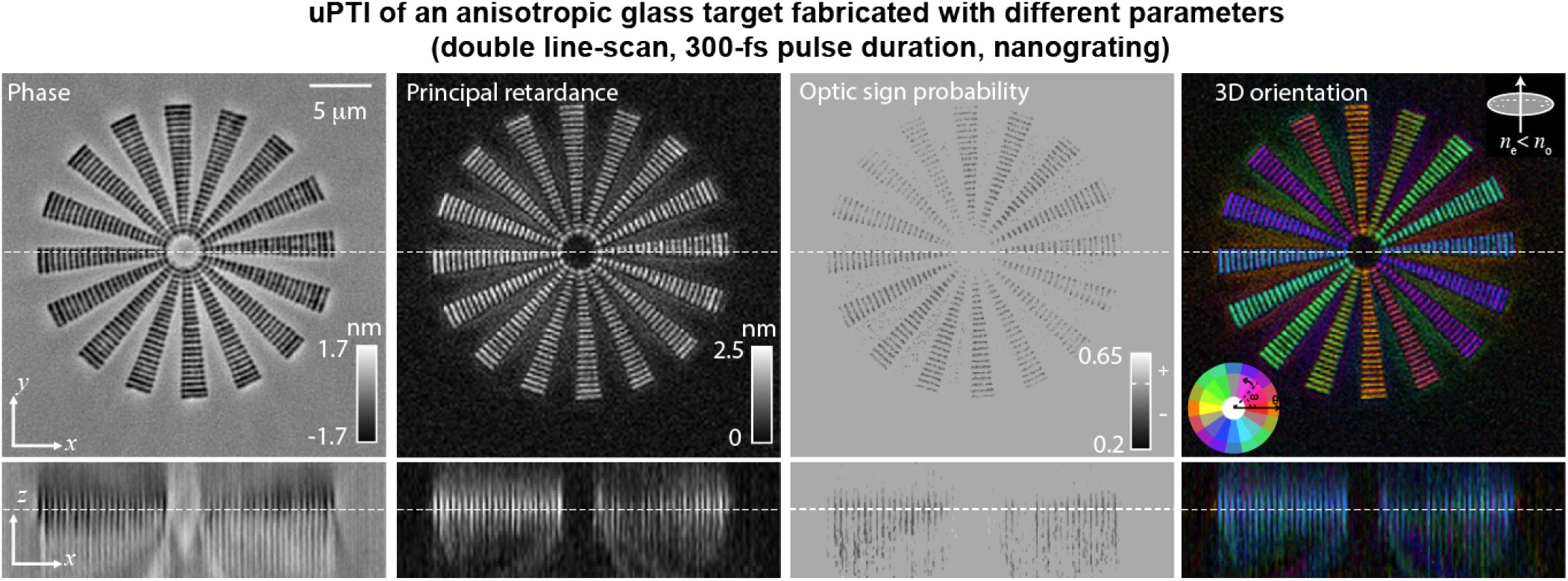
uPTI discriminates the properties of laser written anisotropic glass as writing parameters are tuned: Phase, principal retardance, optic sign probability and 3D orientation of a nanograting birefringent target that was written with 300-fs laser pulse duration, 515 nm wavelength, and illumination NA of 0.55 using a double line scan process, which writes a feature by scanning the laser two times in opposite directions. Differences in the laser parameters relative to the target shown in fig. 2 prevents the creation of two layers of modification as shown in fig. 2C. In addition, the double line scan creates more uniform line features compared to fig. 2C, where we see stronger phase and principal retardance at the end of each grating line within a spoke.

**Figure 2-supplement 3.**
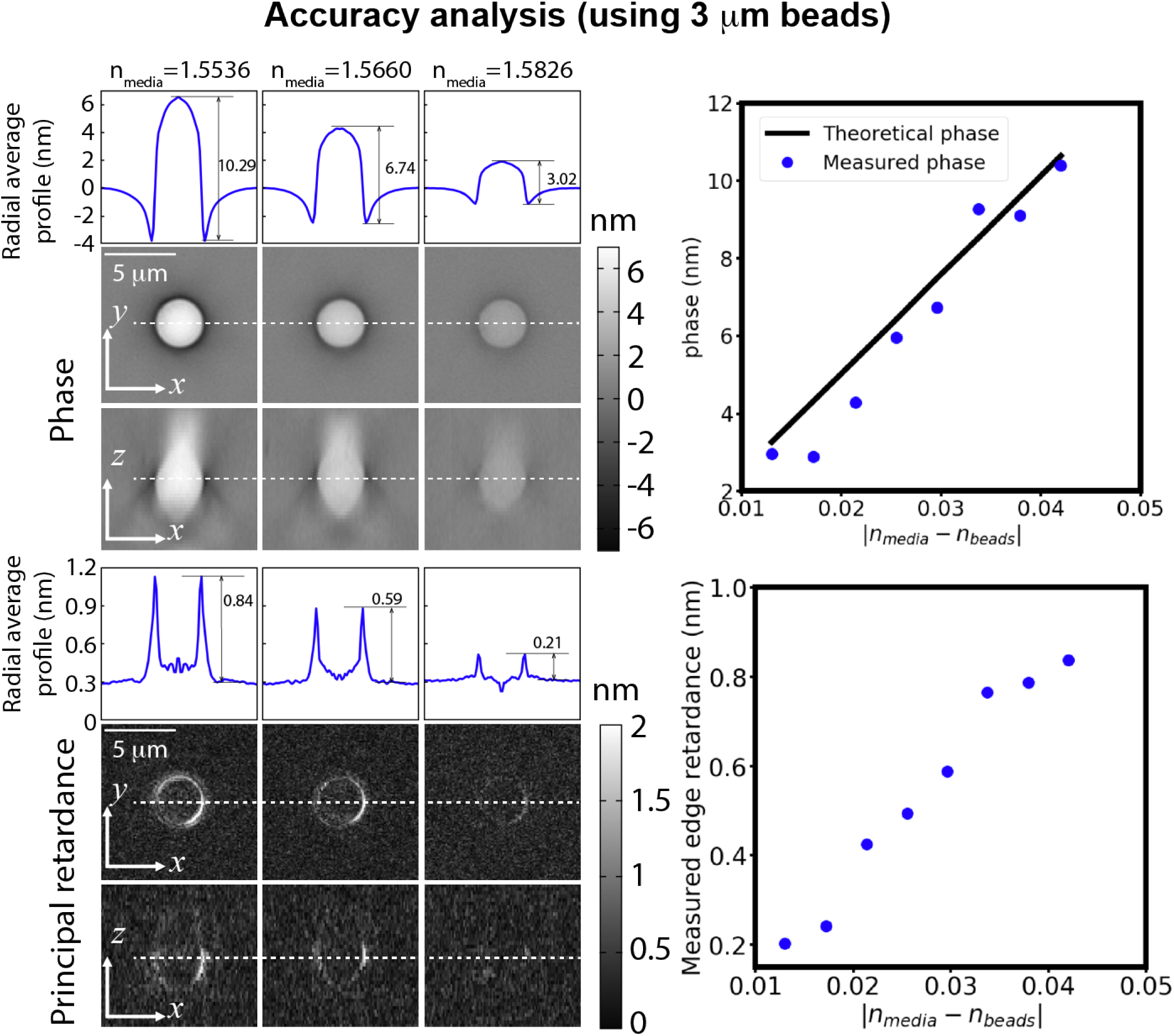
Accuracy of phase and principal retardance: Characterization of the accuracy of phase and principal retardance by imaging 3 μm polystyrene beads immersed in oils with varying refractive indices (RI). Phase and principal retardance images of beads, and their corresponding radial average profiles show that measured phase and retardance decrease as the RI of the immersion oil approaches the RI of the bead. Plots of measured phase, theoretical phase, and edge retardance versus the difference of the RI of the immersion oil and the beads (*n*_beads_ = 1.5956 at 532 nm) show that the measured values follow the expected trend.

**Figure 3-supplement 1.**
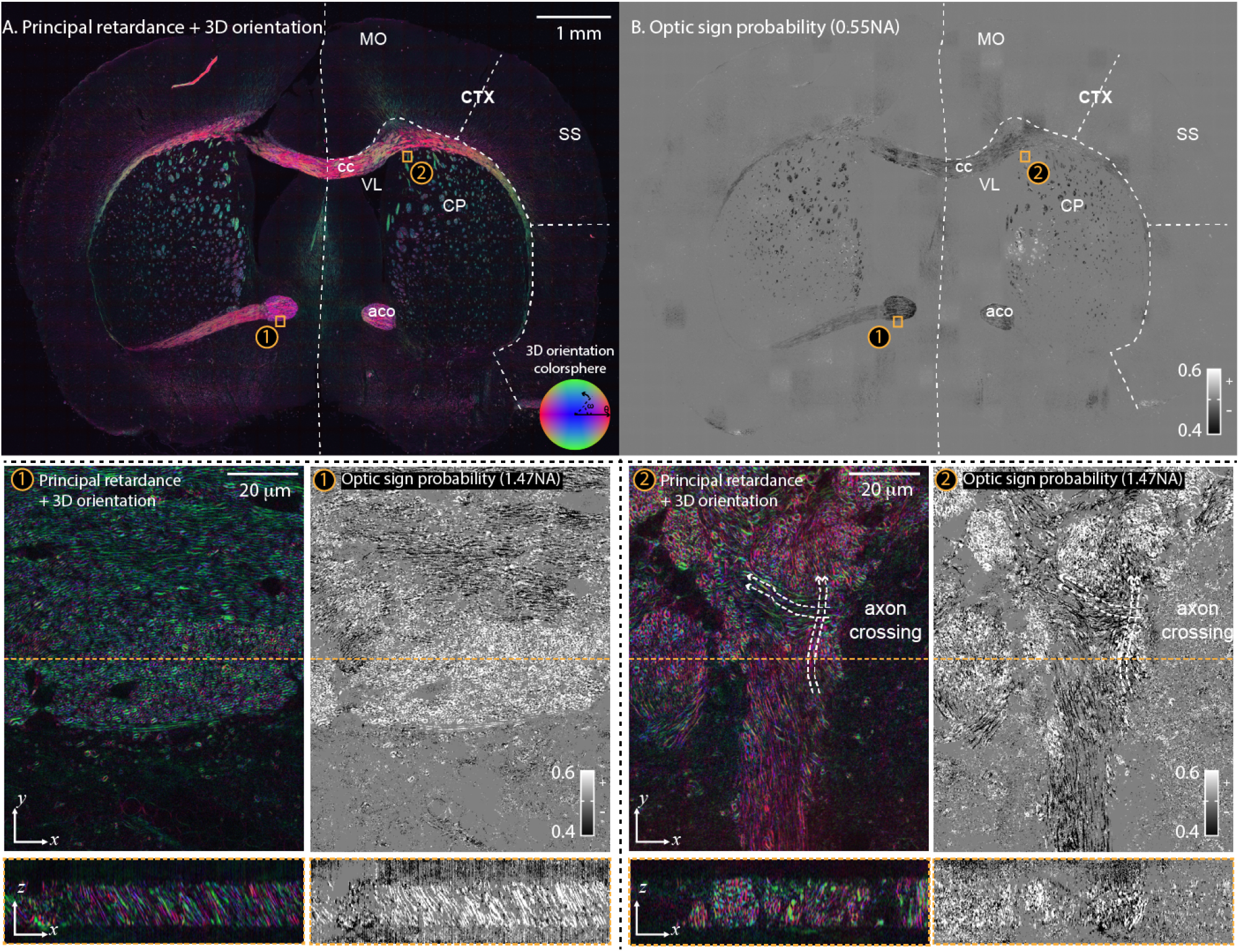
3D orientation with RGB colored sphere and optic sign probability of an adult mouse brain section: (top) **(A)** principal retardance, 3D orientation and **(B)** optic sign probability images of the whole brain section show key anatomical landmarks. At the imaging and illumination NA of 0.55 (spatial resolution of ~ 0.5 × 0.5 × 3.2 μm), axons behave like negative uniaxial material as seen in the optic sign probability. Therefore, we assume the negative uniaxial material when displaying 3D orientation over the whole slide. Multiple anatomical landmarks are labeled according to the coronal section at level 51 of the Allen brain reference atlas, aco: anterior commissure (olfactory limb), cc: corpus callosum, CP: caudoputamen, CTX: cortex, MO: motor cortex, SS: somatosensory area, VL: ventricle. (bottom) aco and CP areas marked with orange boxes (labeled with ① and ②) in (A) and (B) are selected for 3D high-resolution imaging with imaging NA of 1.47 and illumination NA of 1.4 (spatial resolution of ~ 0.23 × 0.23 × 0.8 μm). The orthogonal sections of principal retardance, 3D orientation with RGB colored sphere, and the optic sign probability indicates complex axon network in the mouse brain tissue. With high resolution, we resolve the boundaries of individual axons, which behave as the positive uniaxial material with 3D orientation perpendicular to the surface of the membrane. The RGB colored sphere provides an easier interpretation to see if the structure is (red and green) or is not (blue) in the *x-y* plane.

**Figure 3-supplement 2.**
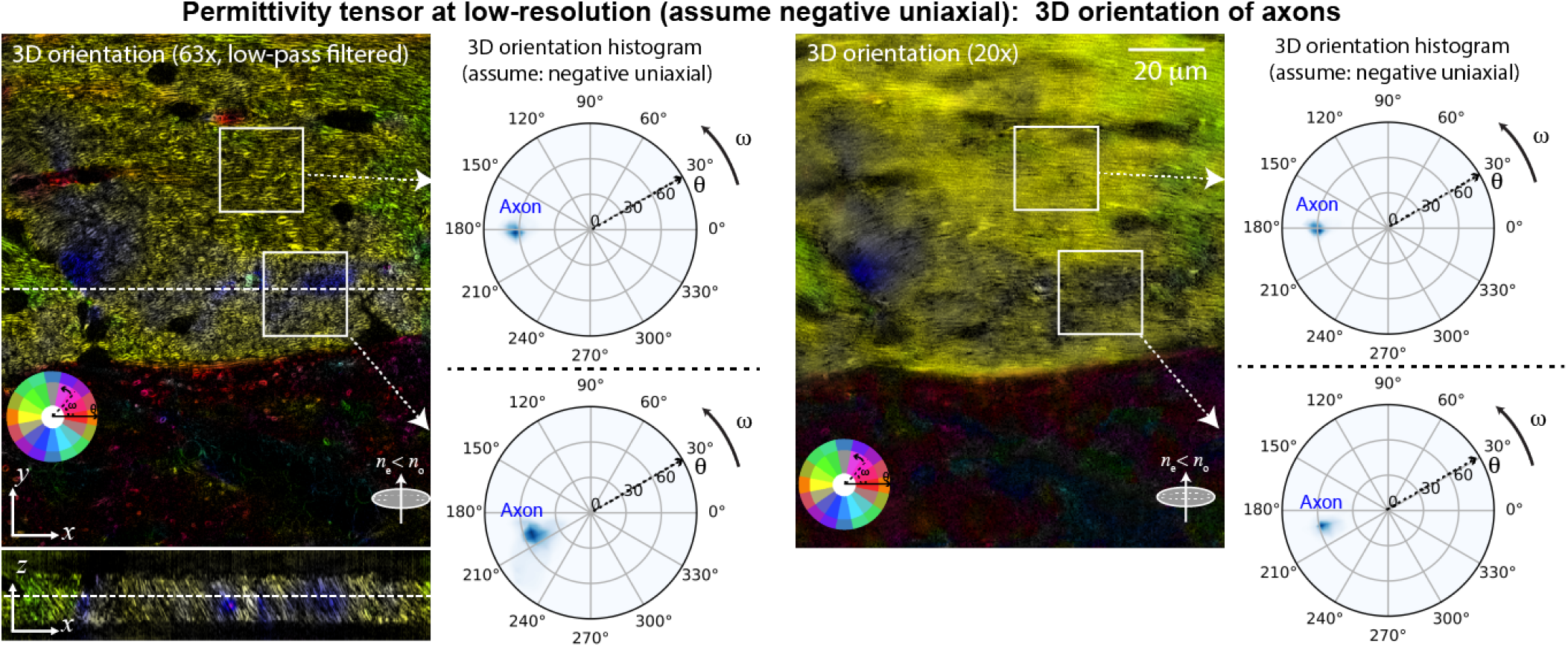
Interpretation of 3D orientation measurements at low resolution: The 3D orientation measurements of two axon bundles (the same region as shown in fig. 3C①) computed from (left) the low-pass filtered scattering potential tensor acquired at 63× objective (1.47 NA) and (right) the scattering potential tensor acquired at 20× objective (0.55NA) assuming the specimen is negative uniaxial. The averaged orientation measurements acquired at 63× show similar distribution of 3D orientation to the distribution measured at 20×, indicating that uPTI provides sensitive measurement of 3D anisotropy across different spatial scales.

**Figure 4-supplement 1.**
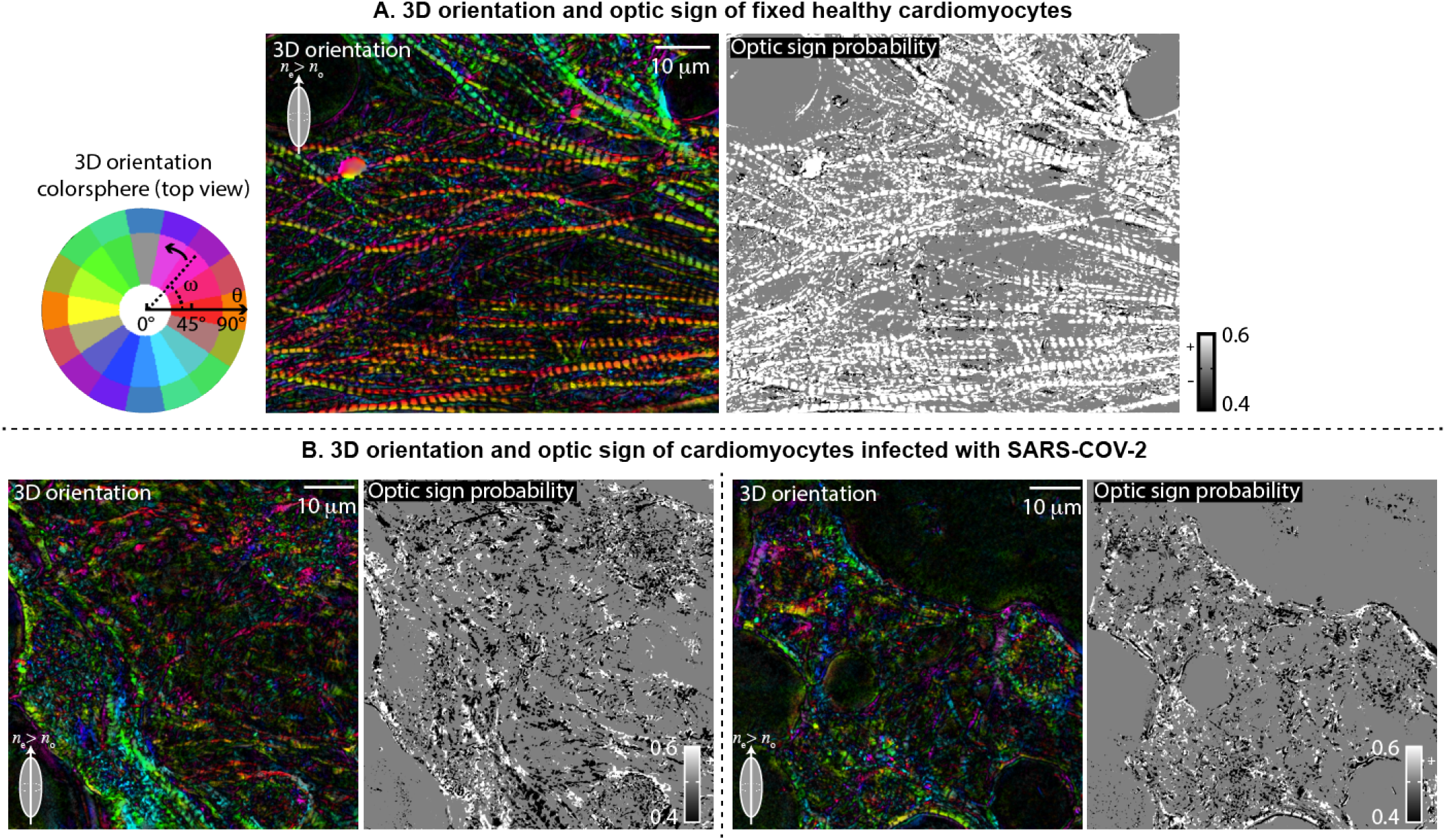
3D orientation and optic sign probability of the iPSC-derived cardiomyocytes: The 3D orientation and optic sign probability of the (A) uninfected and (B) SARS-COV-2 infected cardiomyocytes shown in fig. 4. In the uninfected cardiomyocytes, the optic sign of the sarcomere structure is found to be a positive and 3D orientation is found to point along the axis of the sarcomere. In the infected cardiomyocytes, the principal retardance drops as seen in fig. 4D, which makes the optic sign estimate noisier. The 3D orientation map in the infected cells shows small stretches of consistent orientation that suggests broken sarcomeres, but the orientation estimate is also noisier due to reduced retardance.

**Figure 5-supplement 1.**
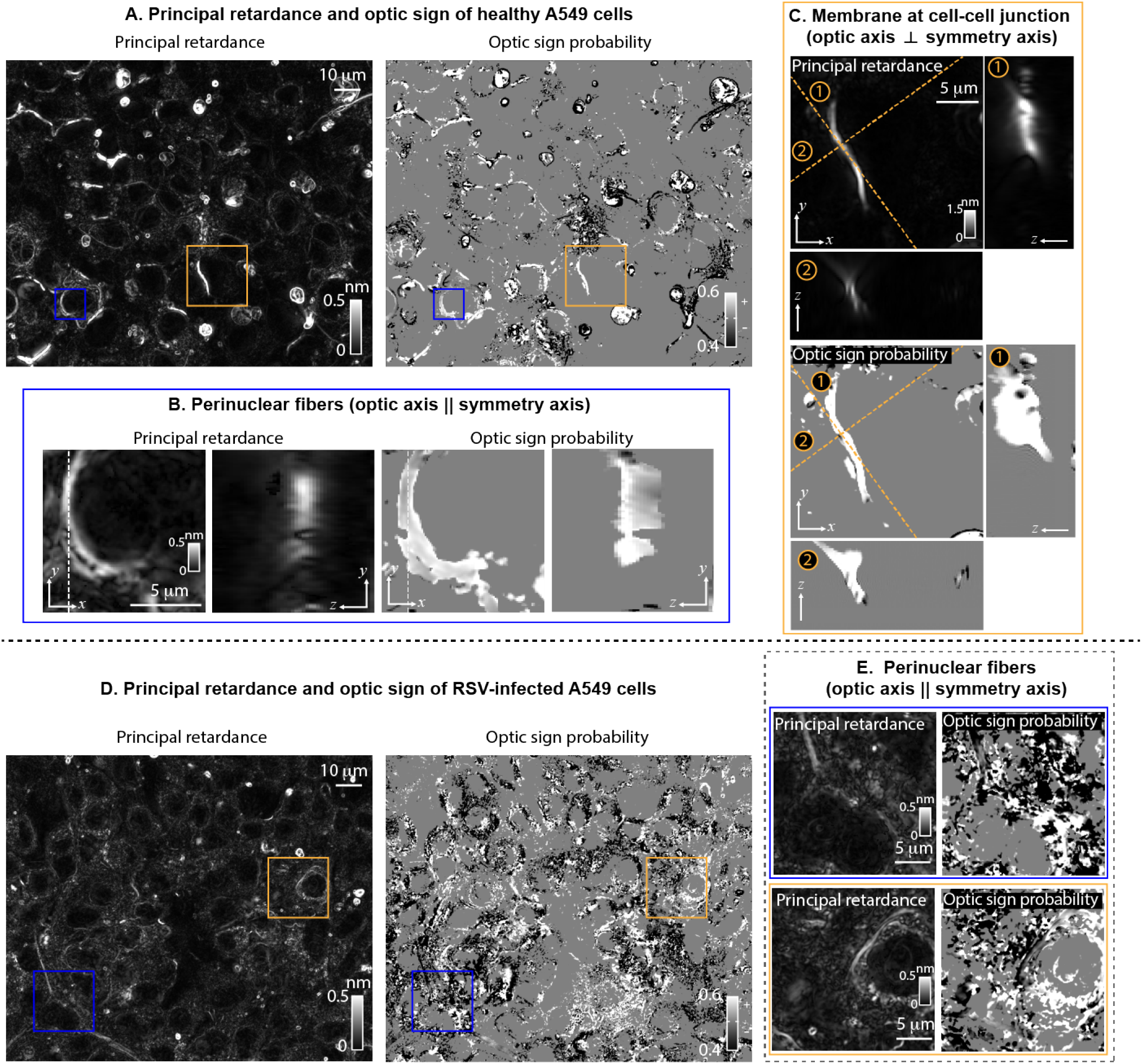
Principal retardance and optic sign probability of the A549 cells: **(A)** Principal retardance and optic sign probability images of the uninfected A549 cells. **(B)** We show an *xy* slice of principal retardance and optic sign probability from the volume in the blue-square region in (A) for the perinuclear structures. Both the *xy* slice and *yz* slice of the principal retardance and the optic sign probability channels indicate this perinuclear structure to be consistently positive uniaxial. **(C)** The orthogonal slices of principal retardance and optic sign probability over the field of view indicated by the orange box in (A) show highly anisotropic structures at the cell-cell junction with a positive optic sign. The orange dashed lines cut at two axes (labeled with ① and ②) tangent and normal to the structure to show its cross-sections in principal retardance and optic sign probability channels. **(D)** A field of view showing the same information as in (A), but for A549 cells infected with RSV. **(E)** Two fields of view chosen from (D) show the perinuclear fibers observed in (A) and (B) with principal retardance and optic sign probability channels.

**Figure 6-supplement 1.**
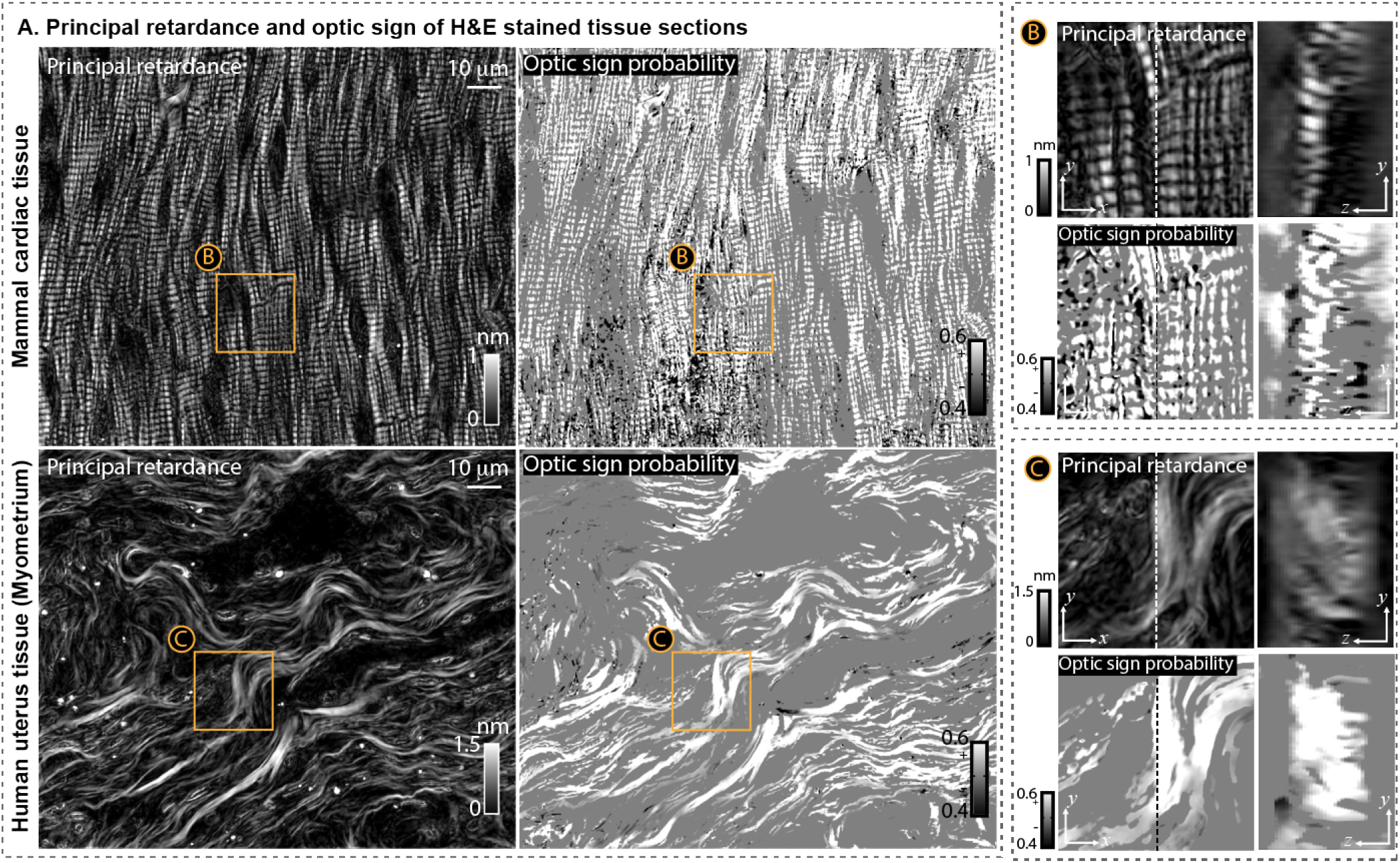
Principal retardance and optic sign probability of the histological sections: **(A)** Principal retardance and optic sign probability of (top) the mammal cardiac tissue and (bottom) the myometrium region of the human uterus tissue. **(B)** The orthogonal slices (*xy* and *yz*) of principal retardance and optic sign probability of the field of view indicated by the orange box in the cardiac tissue. **(C)** Same information as in (B) is shown for the field of view indicated by the orange box in the uterus tissue. Highly anisotropic structures such as A-bands of sarcomeres in the cardiac muscle tissue and collagen fibers in the uterus tissue show consistent positive optic sign estimation through out the volumes.

## Videos

**Video 1.**
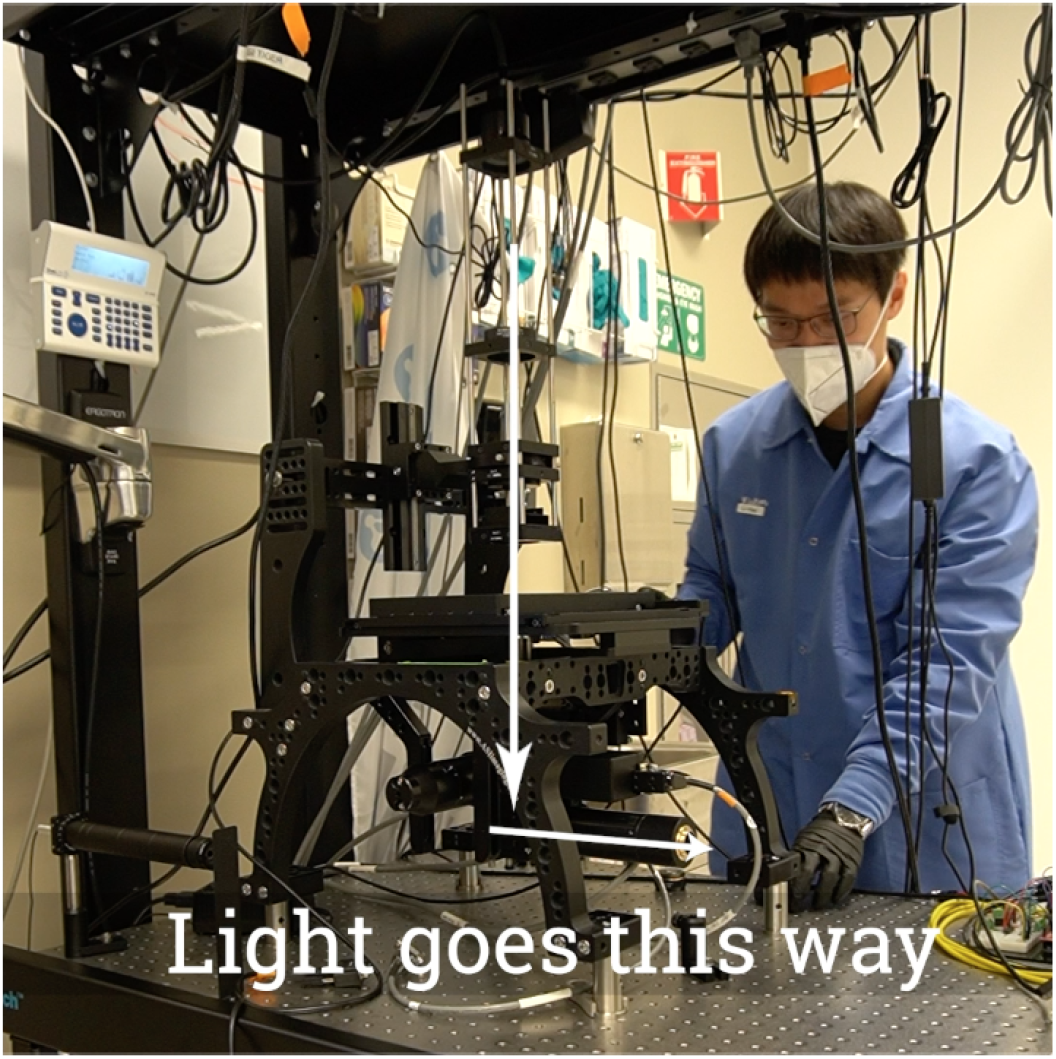
The build video demonstrating how to setup uPTI on a working microscope.

**Video 2.**
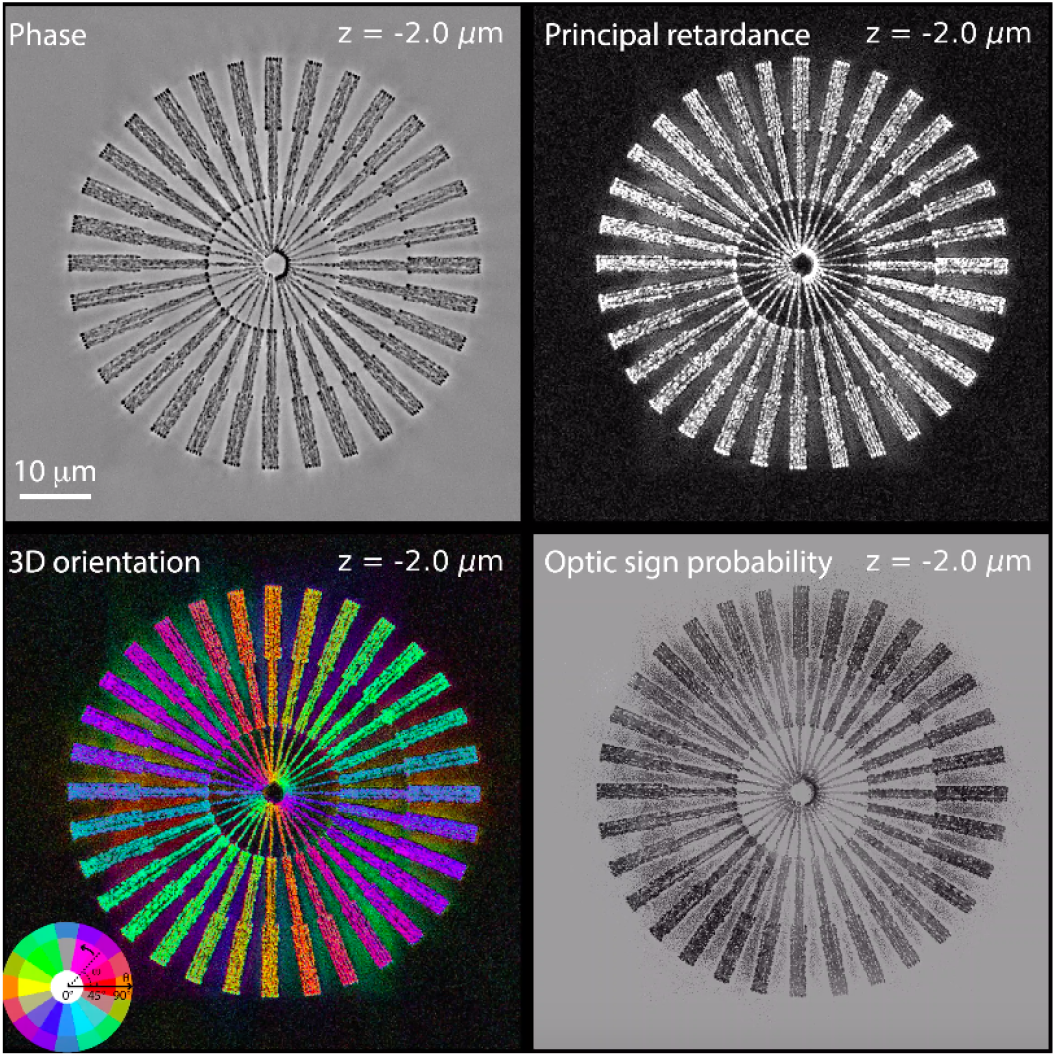
*z*-stacks of phase, principal retardance, 3D orientation and optic sign probability images of the laser written birefringent target shown in fig. 2.

**Video 3.**
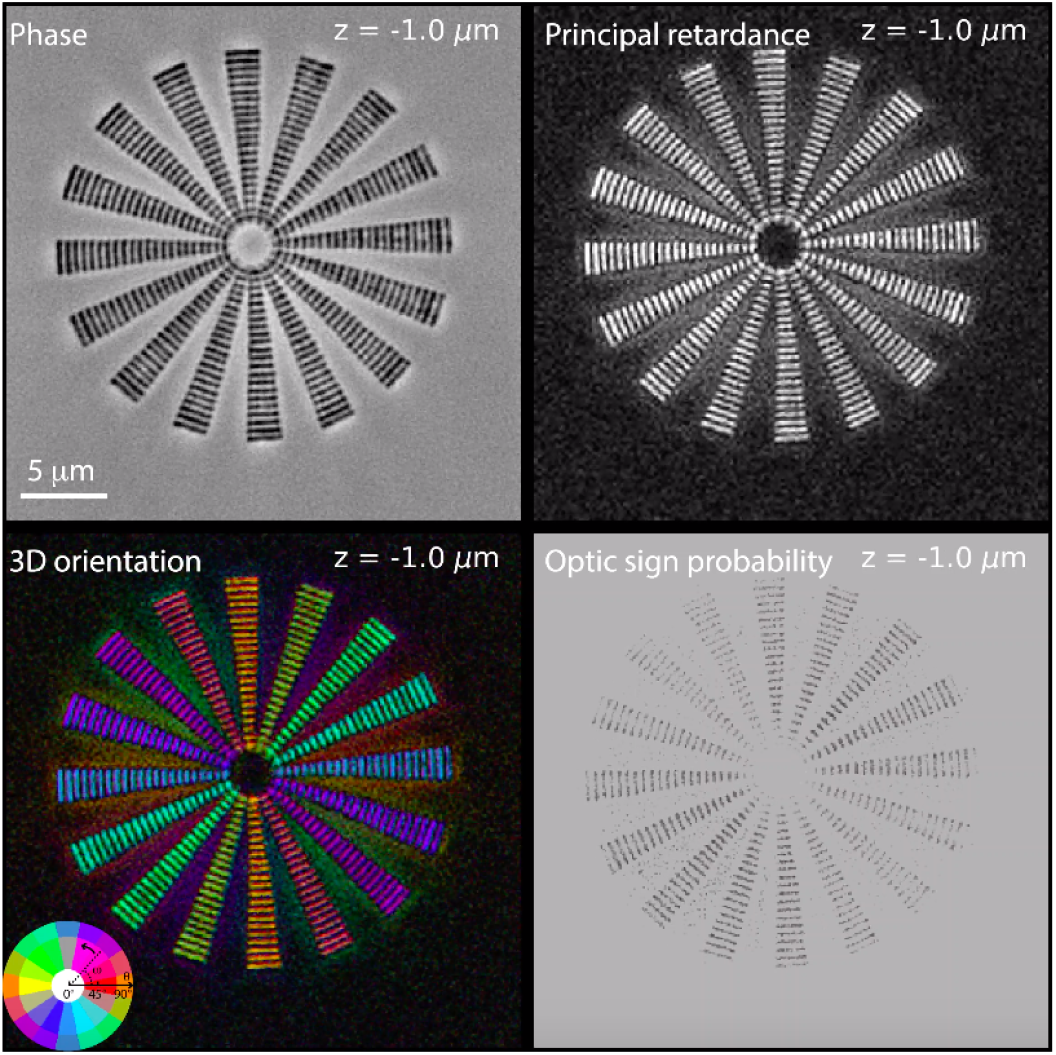
*z*-stacks of phase, principal retardance, 3D orientation and optic sign probability images of the laser written birefringent target shown in Figure 2-supplement 2.

**Video 4.**
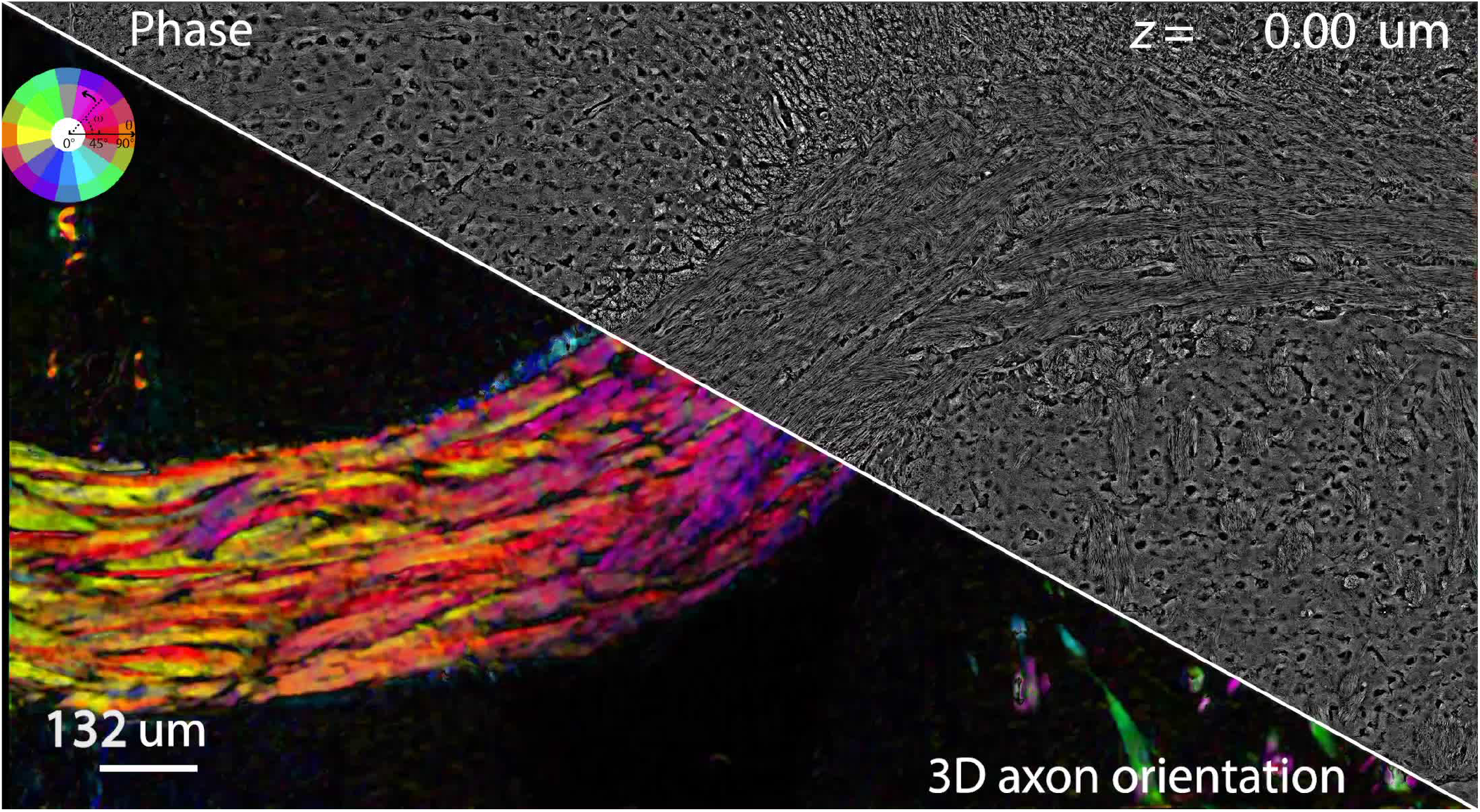
Visualization of 3D high-resolution (1.47NA) phase and principal retardance of the adult mouse brain section.

**Video 5.**
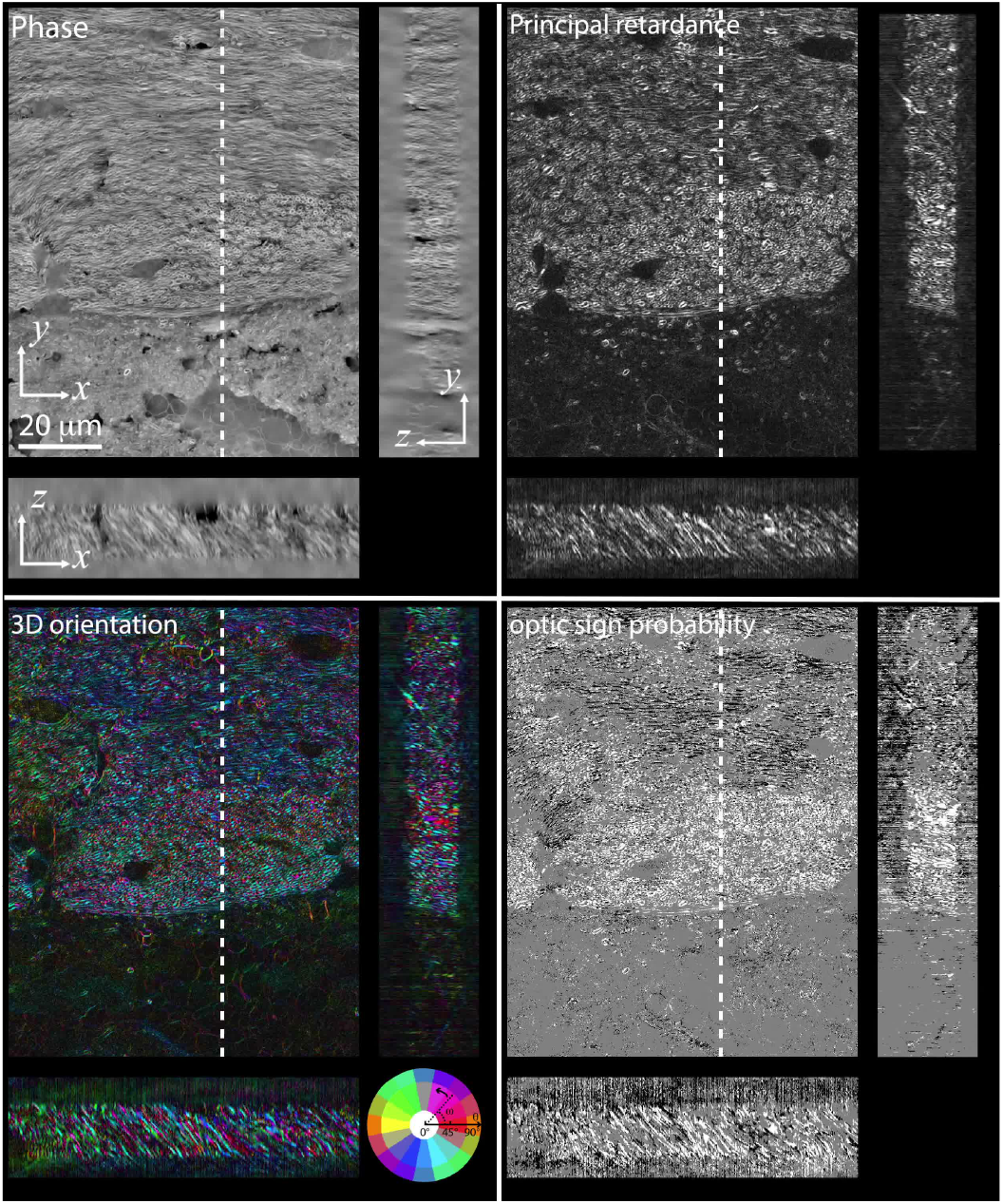
*z*-stacks, *x-z* section, and *y-z* section of phase, principal retardance, 3D orientation and optic sign probability images of field of view ① of the mouse brain section shown in fig. 3.

**Video 6.**
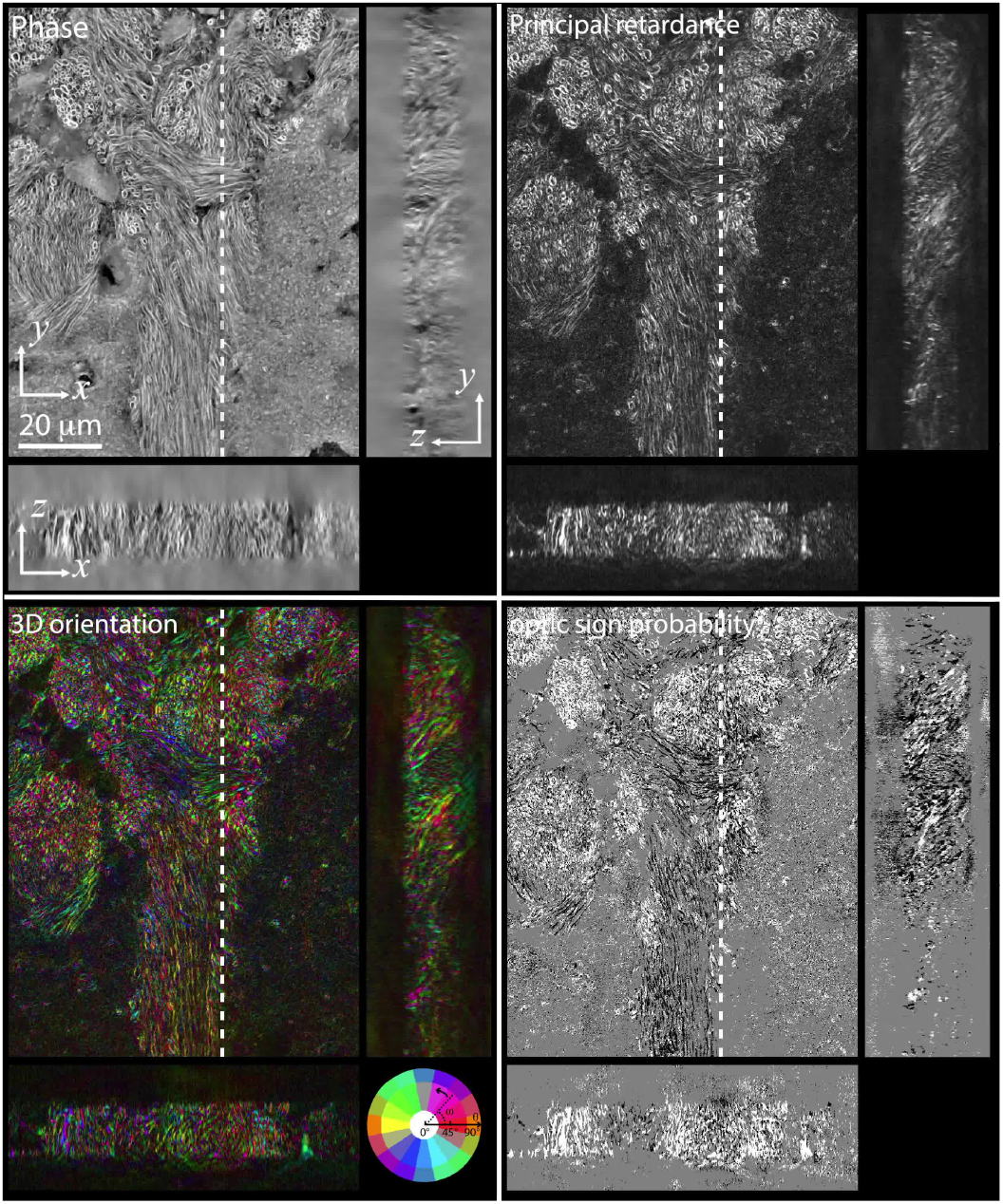
*z*-stacks, *x-z* section, and *y-z* section of phase, principal retardance, 3D orientation and optic sign probability images of field of view ② of the mouse brain section shown in fig. 3.

**Video 7.**
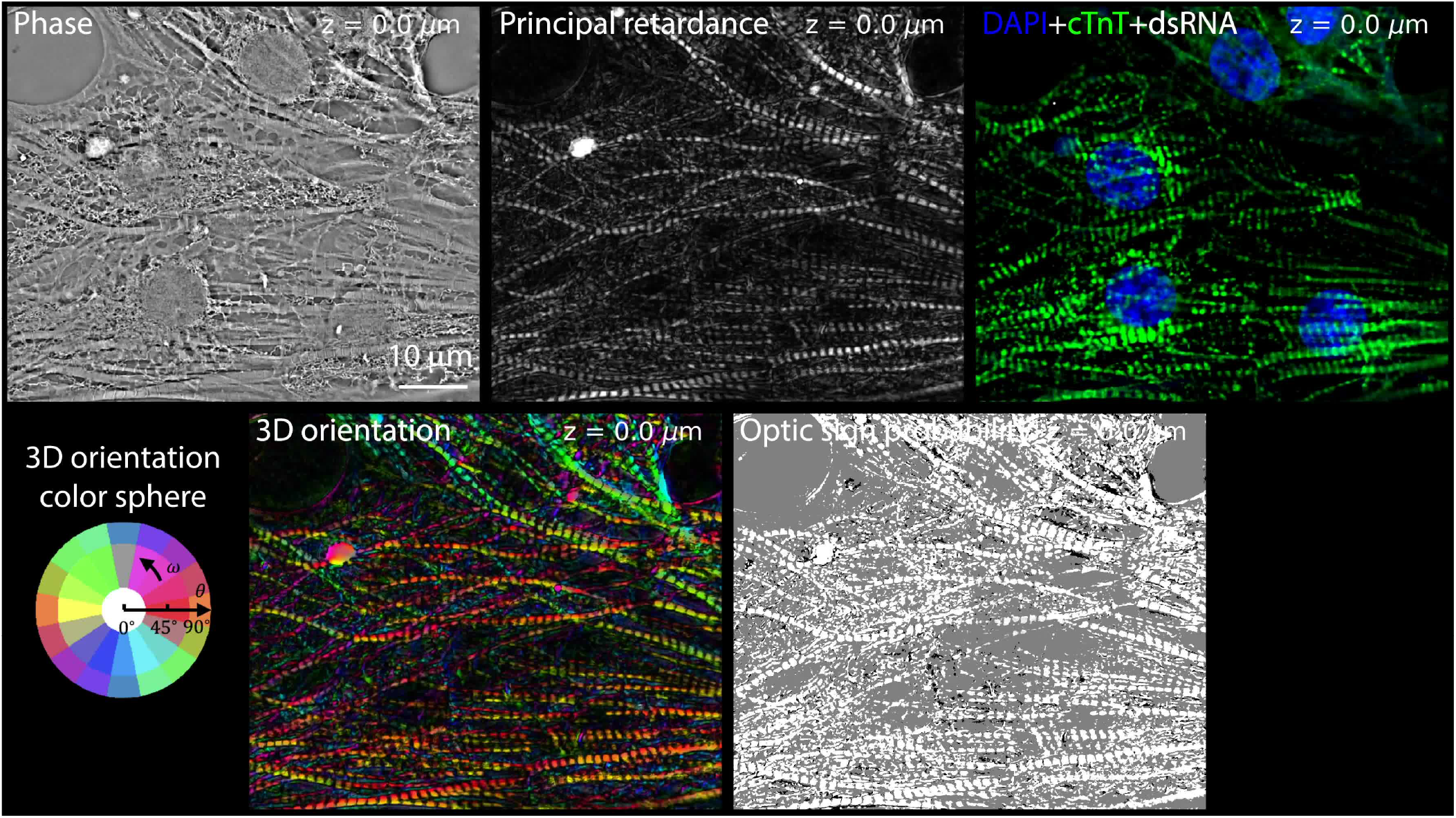
*z*-stacks of phase, principal retardance, 3D orientation, optic sign probability, and fluorescence images of fixed uninfected cardiomyocytes shown in fig. 4A.

**Video 8.**
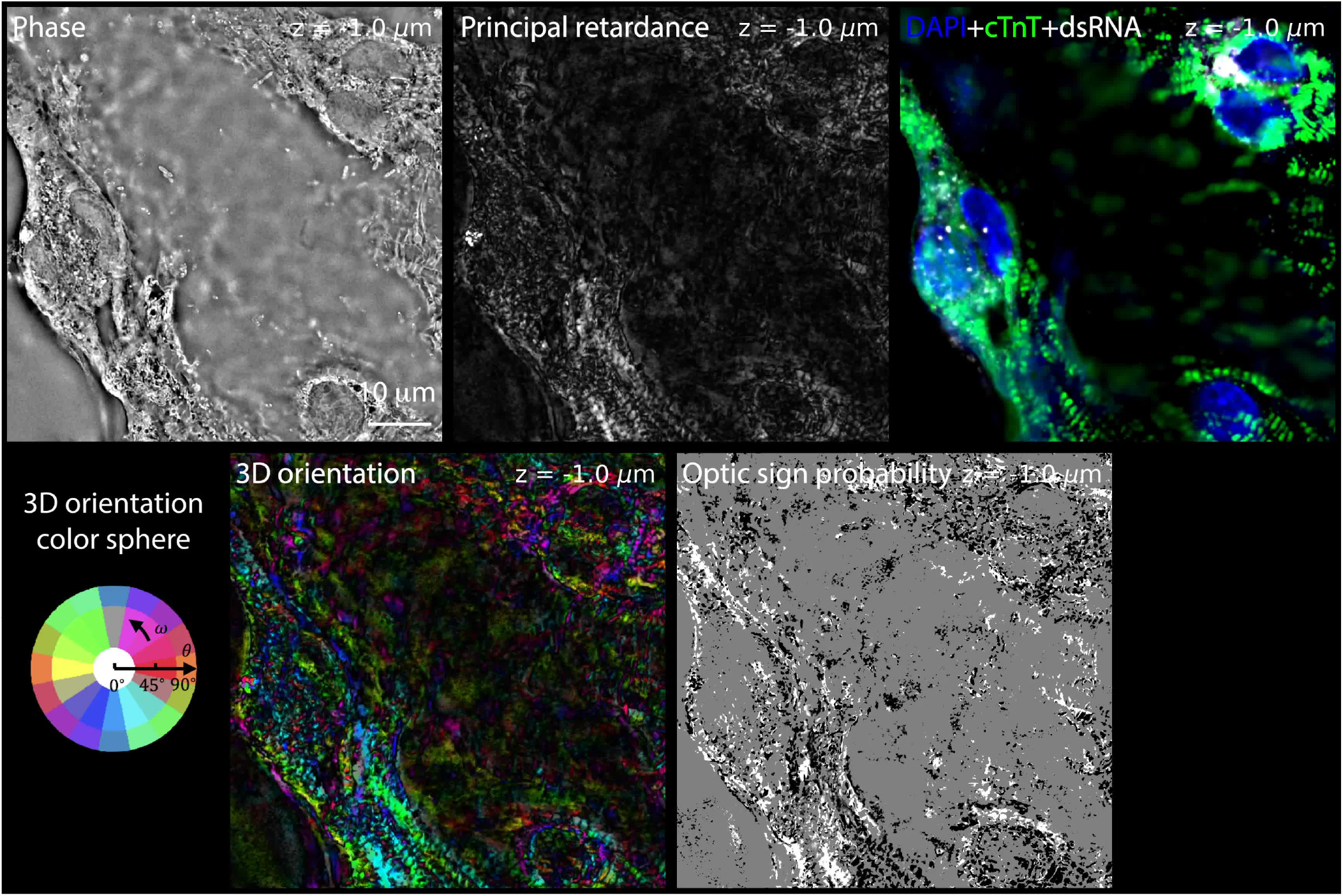
*z*-stacks of phase, principal retardance, 3D orientation, optic sign probability, and fluorescence images of SARS-COV-2 infected cardiomyocytes shown in fig. 4D (left).

**Video 9.**
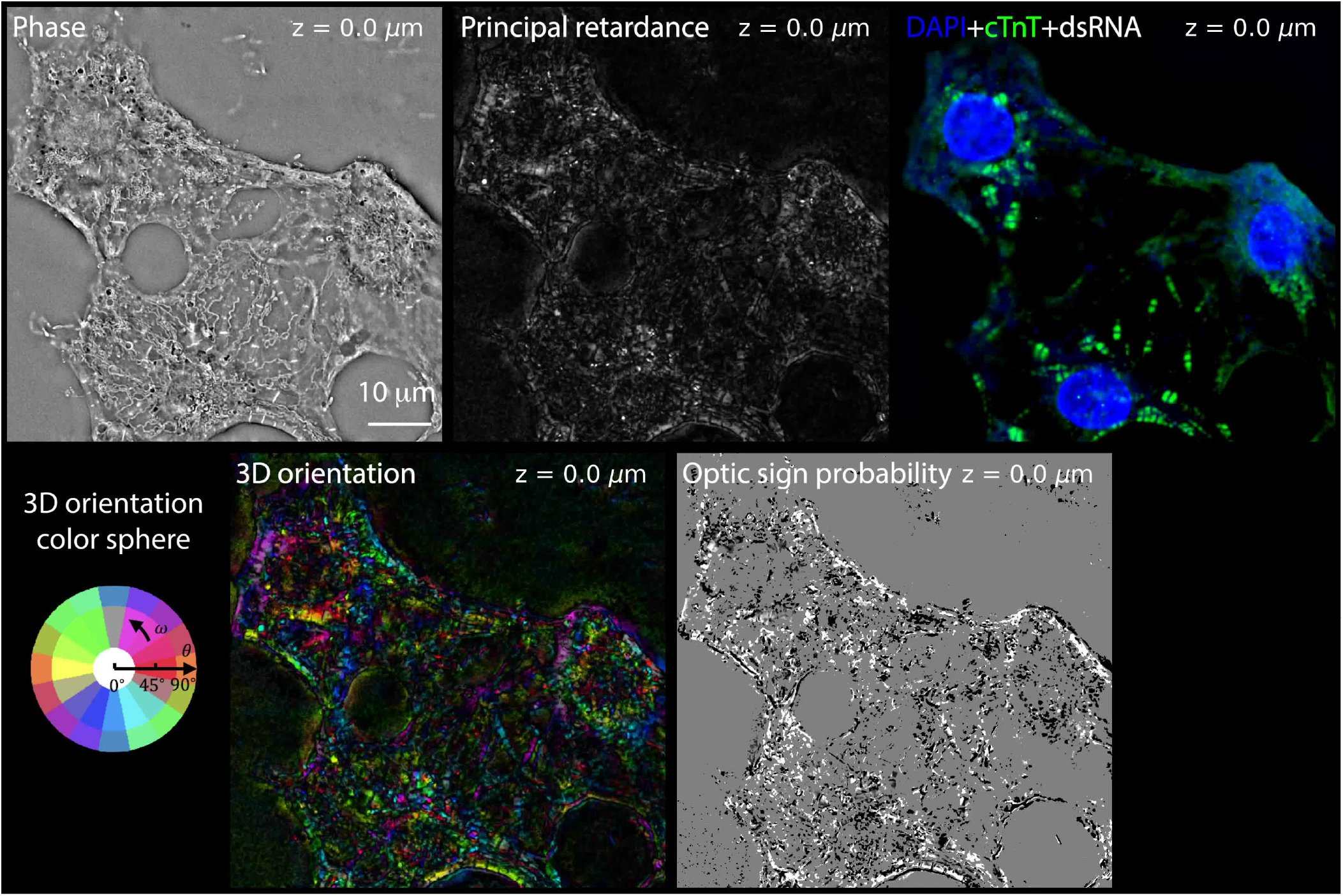
*z*-stacks of phase, principal retardance, 3D orientation, optic sign probability, and fluorescence images of SARS-COV-2 infected cardiomyocytes shown in fig. 4D (right).

**Video 10.**
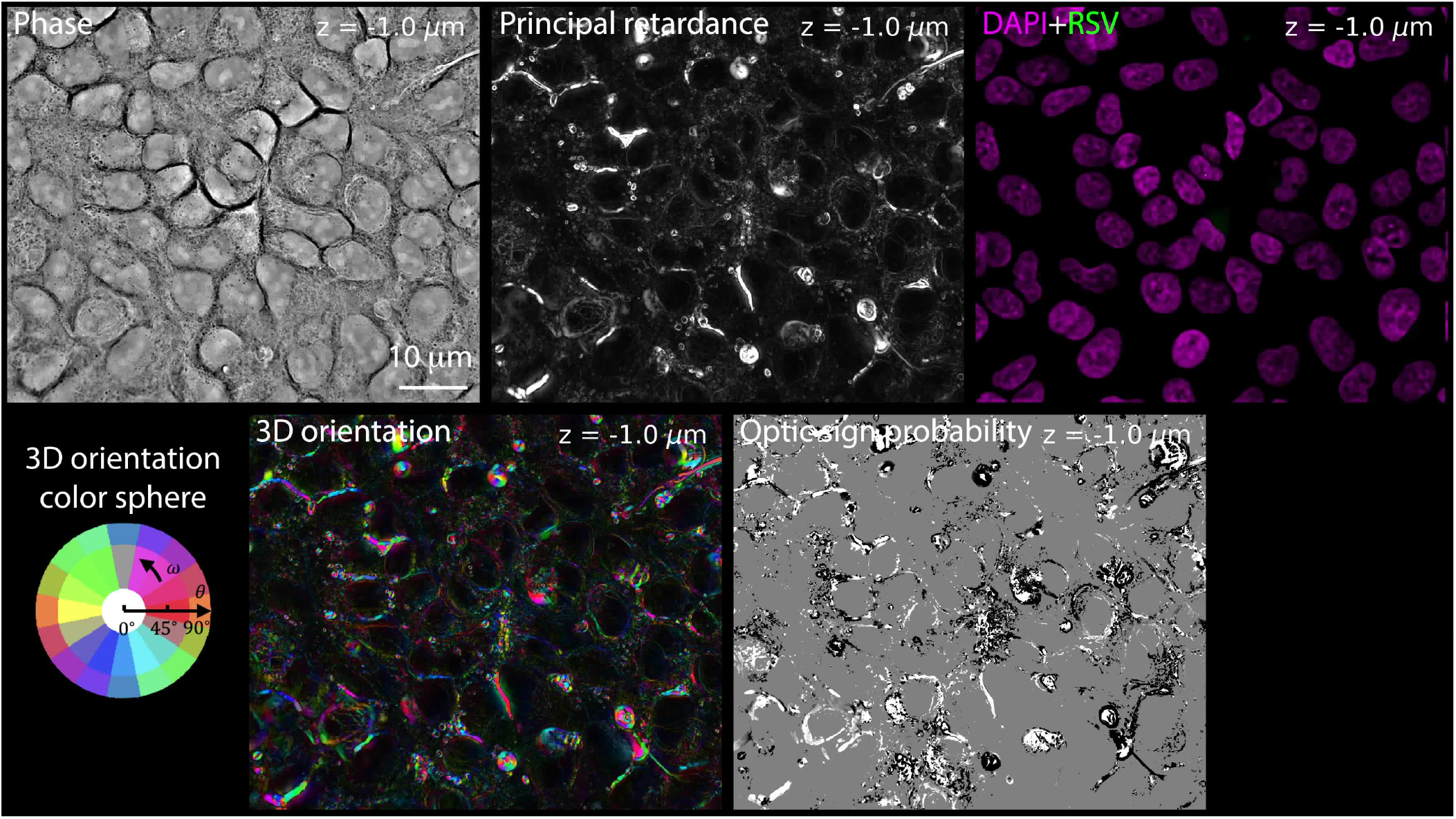
*z*-stacks of phase, principal retardance, 3D orientation, optic sign probability, and fluorescence images of fixed uninfected A549 cells shown in fig. 5A.

**Video 11.**
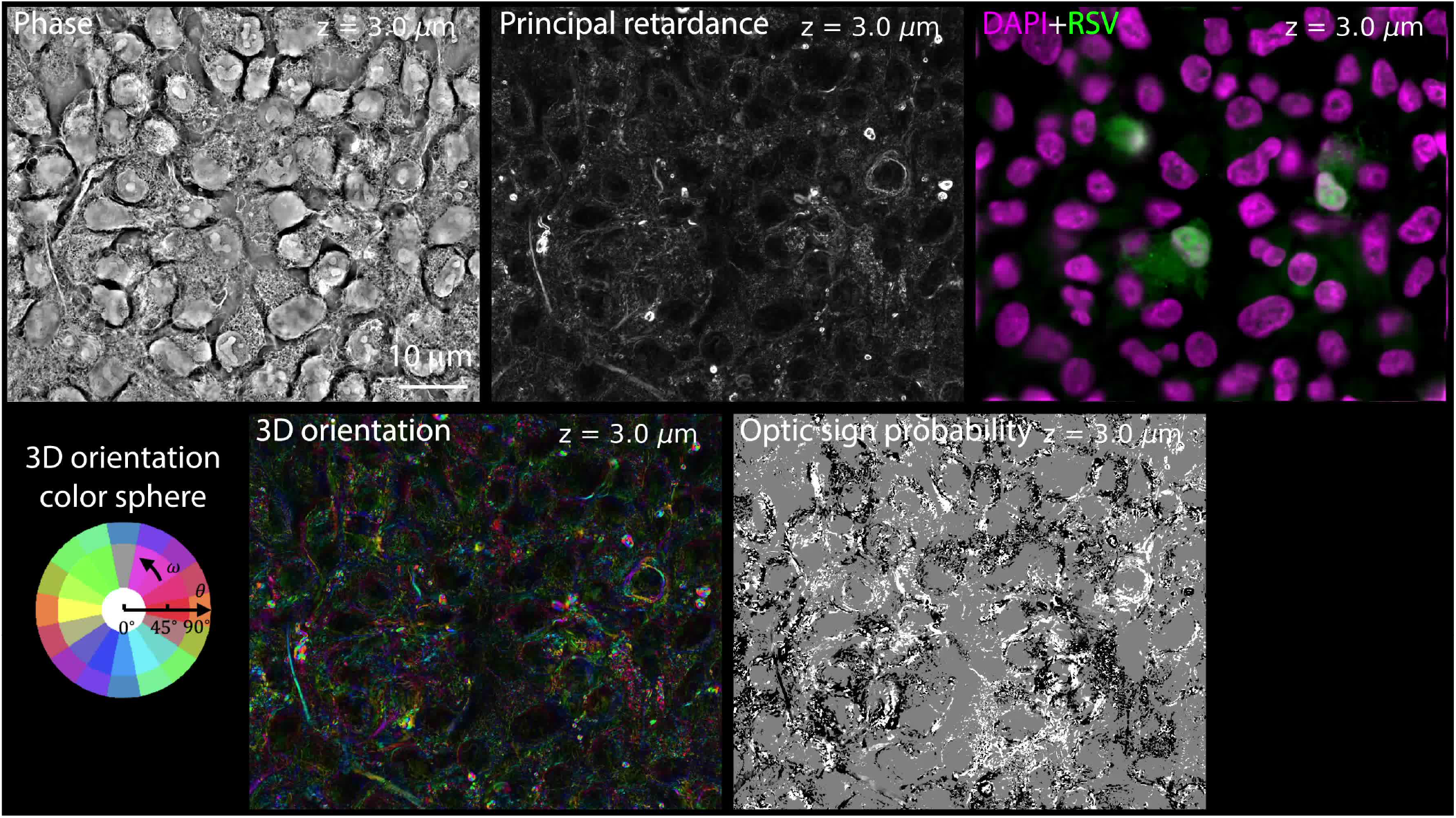
*z*-stacks of phase, principal retardance, 3D orientation, optic sign probability, and fluorescence images of RSV-infected A549 cells shown in fig. 5D.

**Video 12.**
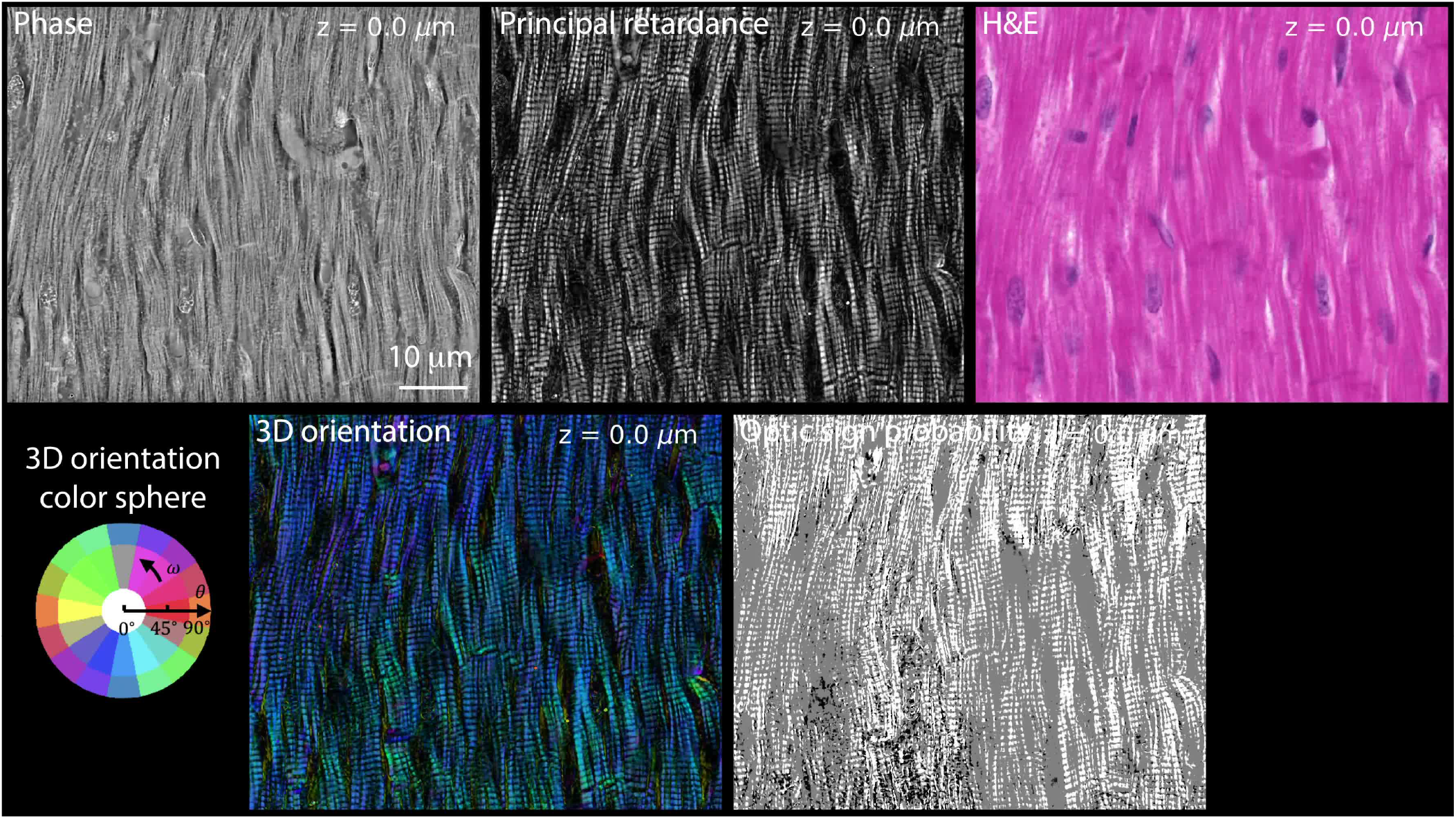
*z*-stacks of phase, principal retardance, 3D orientation, optic sign probability, and H&E images of the mammal cardiac tissue shown in fig. 6A (top).

**Video 13.**
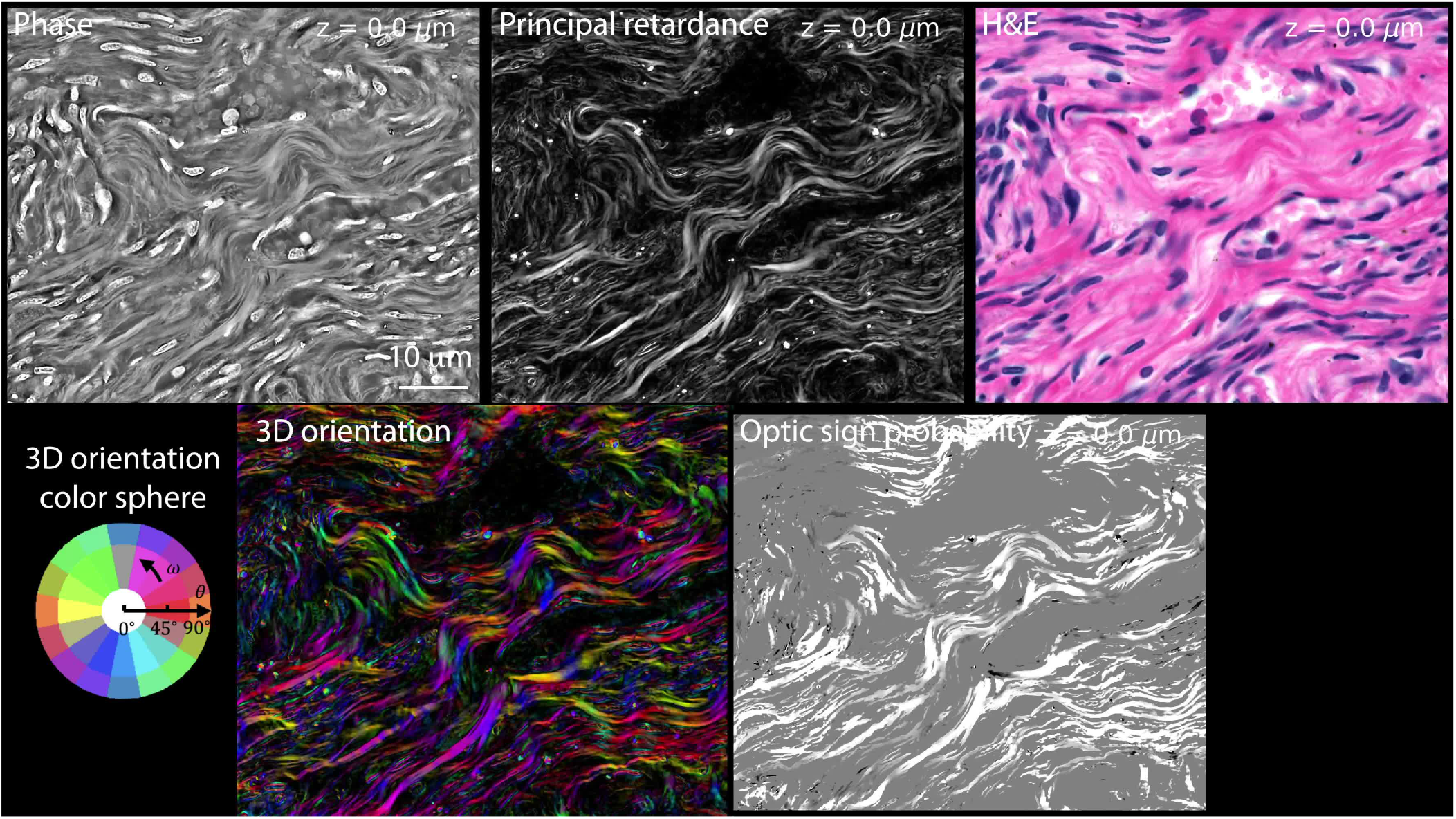
*z*-stacks of phase, principal retardance, 3D orientation, optic sign probability, and H&E images of the human uterus tissue shown in fig. 6A (bottom).

## Supplementary Note 1: Derivation of the forward model

3D distribution of optical phase and anisotropy of a specimen arise from the 3D distribution of the material’s relative permittivity. This permittivity determines the amount of average and differential optical phase shift and the characteristic axes responding to different input polarization modes, which corresponds to specimens’ density and anisotropy information. In the following subsections, we explain how the information of this permittivity is coupled to measurable quantities through vector wave equation (1, 2) and partially coherent imaging model (3–5).

### A: Permittivity tensor

Permittivity is a measure of how much a material is polarized by an applied electric field. The more easily a material is polarized, the slower a eletromagnetic wave is travelling in it, and more optically dense the material is. A polarized material radiates with a group of induced electric dipoles and creates measurable scattered fields. Understanding the permittivity of a material is key to extracting scattering-related optical quantities such as optical phase and anisotropy.

The permittivity is a scalar for an isotropic material such as water. The result of a scalar permittivity is that the induced electric dipole moment aligns and scales linearly with the applied electric field. In the case of an anisotropic material, the induced electric dipole moment still scales linearly with the applied electric field. However, its direction is dependent on the angle between the direction of the applied electric field and the orientation of the optic axis of the material. The permittivity of an anisotropic material hence takes a form of a tensor (6).

There are two types of anisotropic material depending on the number of optic axes they have, the uniaxial material and the biaxial material (6). Here we consider biological structure with at most one anisotropic axis (an uniaxial material). Assuming that the imaging system’s optical axis is aligned along the *z*-axis as shown in fig. 1. We express the relative permittivity tensor of an uniaxial material oriented in the *z*-axis as

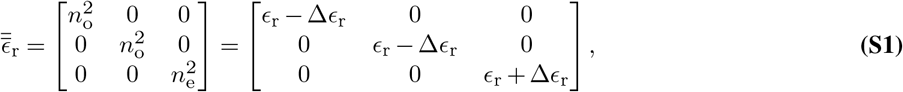

where *n_o_* and *n_e_* are refractive indices experienced by the ordinary and extraordinary wave, respectively. Refractive indices are generally complex numbers, whose real part causes phase delay and imaginary part introduces absorption. For the convenience of the following derivation, we express this permittivity tensor with the average permittivity (*ϵ*_r_) and the difference of permittivity (Δ*ϵ*_r_) from the ordinary and etraordanary wave. They are related to the refractive indices as follows:

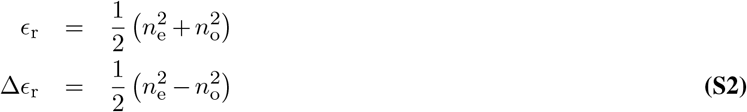

To generally understand the induced dipole moment of an arbitrarily oriented uniaxial material, we consider the material oriented with in-plane orientation of *ω* and inclination of *θ* as shown in fig. 1B. The permittivity tensor in the microscope’s frame of reference is obtained through applying rotational transformation on 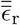 and is expressed as (7)

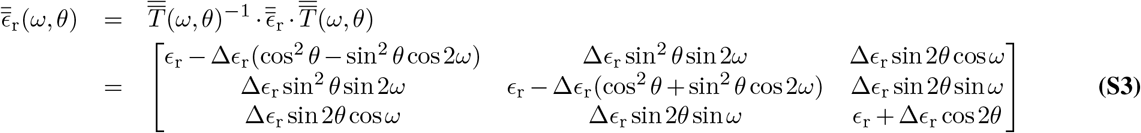

where 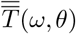 is the overall coordinate rotational matrix that is composed of two consecutive coordinate rotations around *z* and *y* axes as

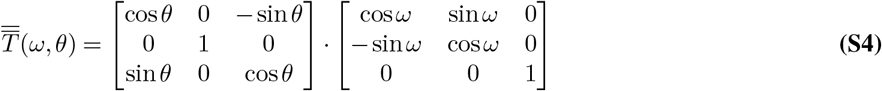

With this general form of permittivity, we will move to introduce how the induced dipole moment creates scattered electric fields that encode information of this permittivity tensor.

### B: Vector wave equation and Born approximation

Following the scalar wave scattering theory (6), the core understanding of wave scattering event lies in the combined solution of homogeneous and inhomogeneous wave equation. In order to account for a more accurate description of the interaction between a polarized light and an anisotropic material, we start by considering the vector wave equation derived from Maxwell’s equation as (1, 2)

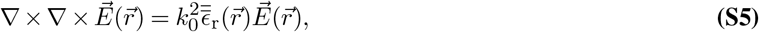

where 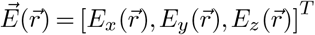 is the electric field in 3D space 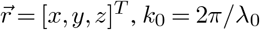 is the free-space wavenumber, and *λ*_0_ is the free-space wavelength of the light. Our goal here is to find a general solution of the electric field in this equation. To solve this equation, we rewrite this equation as

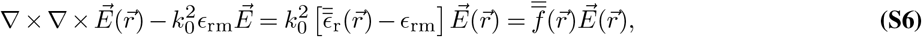

where *ϵ*_rm_ is the isotropic relative permittivity of the surrounding media and is a scalar value and 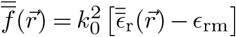 is the scattering potential tensor. Setting the right-hand side of the Eq. (S6) to zero, we get a homogeneous solution 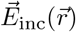, which can be simply an incident propagating plane wave. Treating the right-hand side of the Eq. (S6) as a source term, people have found its inhomogeneous solution as (1, 2)

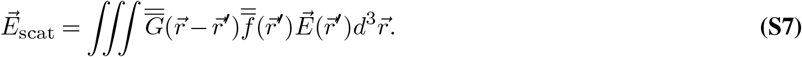

The 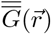 is the dyadic Green’s tensor expressed as (1, 2)

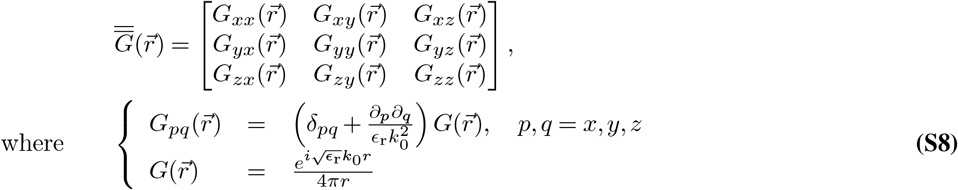

Combining the homogeneous and inhomogenous solution, we get a solution in the following form

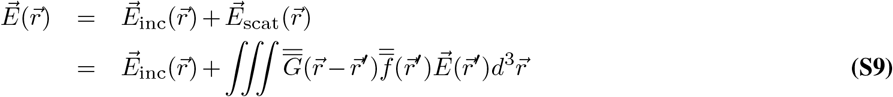

The interaction term of scattering potential tensor 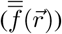 and the total electric field 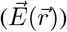 acts as an induced dipole moment radiating electric field to the space through the dyadic Green’s tensor. This radiating field is the scattered field that is triggered by the incident electric field. The total field is sum of the incident and the scattered fields. However, this is not yet a solution, because the scattered field depends on the knowledge of the total electric field, which is a sum of the incident and scattered fields. A recursive approach (8, 9) is needed for a complete solution.

Rigorous approaches to solve the total electric field can be computationally intensive (8, 9). Here we consider biological specimens that are relatively weakly scattering. Hence, we are able to replace 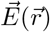 with 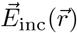 in Eq. (S7) by assuming the first Born approximation (6) and get

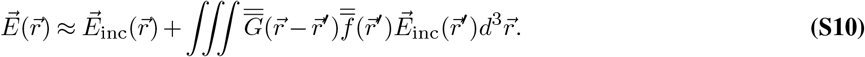

This equation explains how the material is interacting with the incident electric field under weakly scattering condition.

### C: 3D vectorial partially coherent imaging model

Here, we first establish the link from the scattered fields to measurable generalized Stokes parameters (3, 4) under partially coherent illumination. The relationship between the measurable Stokes parameters and the permittivity tensor (or scattering potential tensor) is generally nonlinear. In order to develop a more computationally efficient algorithm for the permittivity tensor retrieval, we sacrifice some accuracy by linearizing the model with the approach similar to (10–14). We will show this linearization leads to handy transfer functions for further reconstruction of permittivity tensor. In the end, we follow this model with a reduction of dimensionality to consider 2D imaging case.

#### C.1: Electric field of an incident plane wave

Our polarization microscope measures Stokes parameters of scattered light from the specimen under quasi-monochromatic spatially partially coherent illumination in a Köhler geometry. Each point on the source plane (aperture plane of a illumination condenser lens) generates a plane wave with 3D spatial frequency of 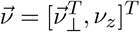, where 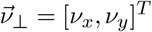 is the transverse spatial frequency and *ν_z_* is the axial spatial frequency satisfying 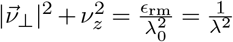. *λ* is the wavelength of light inside a media with permittivity of *ϵ*_rm_. This plane wave is expressed as

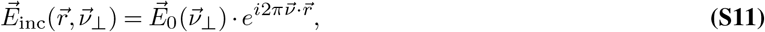

where 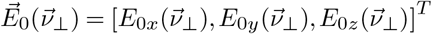 is the illumination-angle-dependent vector component of the incident plane wave from the lens-focusing effect.

Considering the polarization state of the light is known as 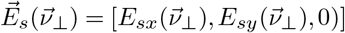 in the source plane, which is usually the case experimentally, we relate it to the illumination-angle-dependent electric field of the incident wave through the lens focusing effect (15) as

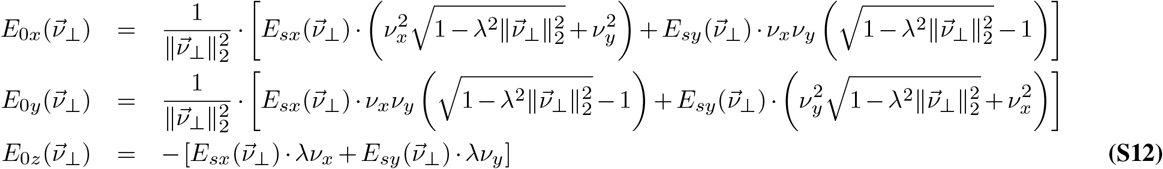

#### C.2: Scattered field to generalized Stokes parameters

For a Köhler geometry, each independent coherent plane wave coming from the source plane interacts with the specimen through Eq. (S10) to create scattered field and each *z*-plane of this 3D scattered field is convolved with a 2D coherent point spread function 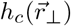 to form a 3D stack of output total field 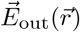. By expressing the scatter potential tensor as

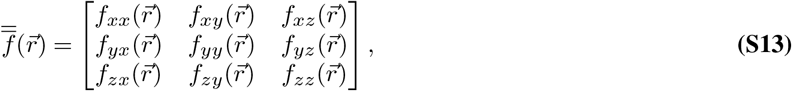

we express one arbitrary component of 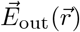 as

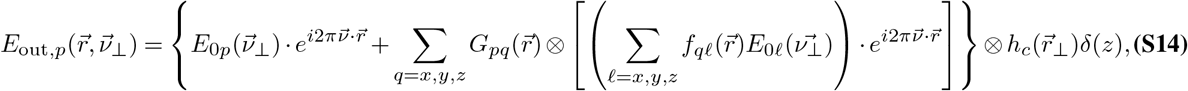

where *p* = *x,y,z,* ⊗ denotes 3D convolution, and 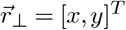 is the transverse coordinate of 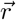.

We have so far derived the scattered electric field on our detector from a single plane wave in a partially coherent source pattern. Different partially coherent source patterns (*α*) are specified by different sets of 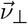. By integrating the contribution of all the coherent scattered modes from one illumination pattern, we obtain the general Stokes parameters (3, 4) as

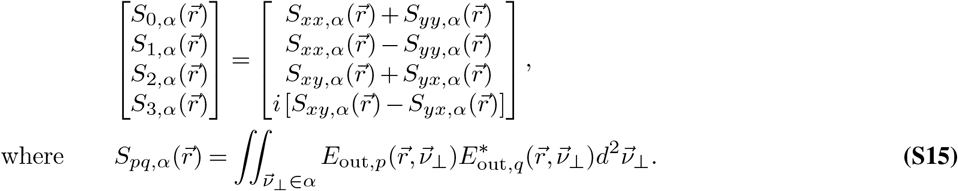

For ease of the following analysis, we take a closer look at the Fourier space of 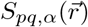 as

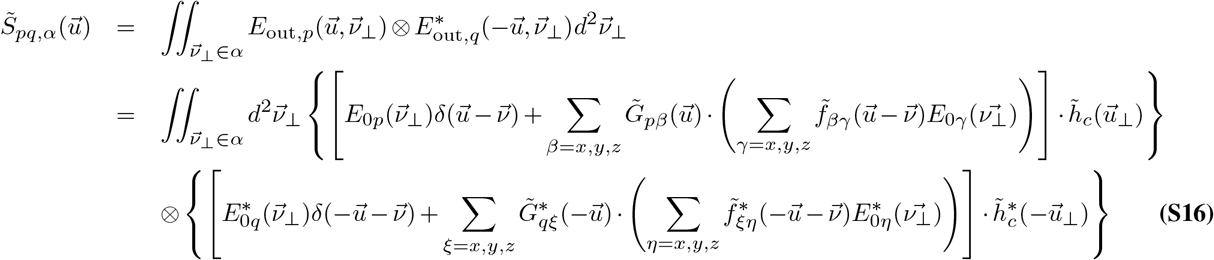

where 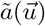 denotes the Fourier transform of a function 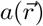 at the 3D spatial frequency 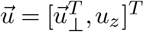. From this equation, we observe the measurable Stokes components, 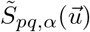, are nonlinearly related to our targeted information, 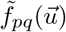. To solve this rigorously, it requires an iterative optimization algorithm as in the classic phase retrieval problem (16), which can be computationally heavy (time consuming) and subject to convergence instability (17). Instead, we adopt the linearization approach to simplify this equation for simpler, faster, and more robust solution.

#### C.3: Linearization of the model

Following previous section, we expand Eq. (S16) further into three major contribution terms

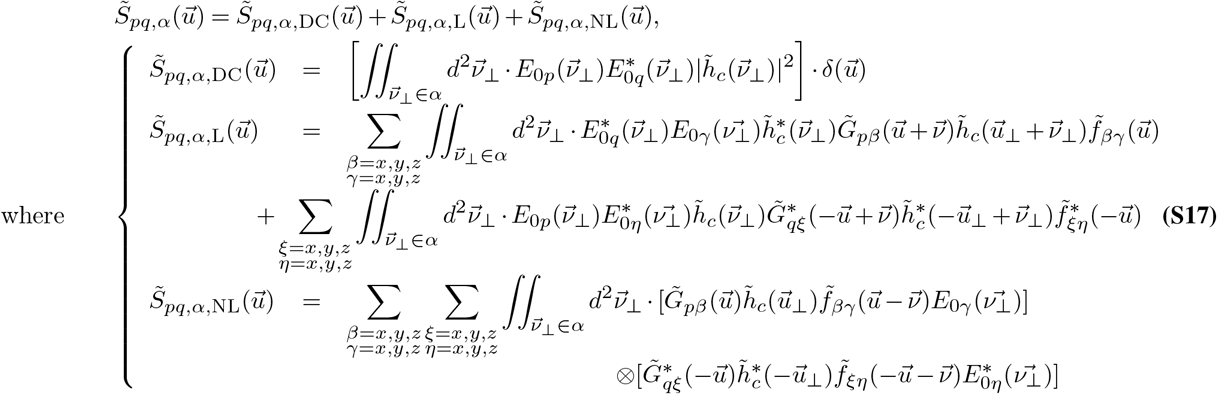

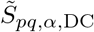 is the DC component of a Stokes component, 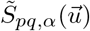, that has the strongest energy that comes from the incident electric field. 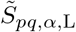 is the term that the scattering potential tensor are linearly related to the Stokes component. This is the second dominant term and it comes from the interference of the scattered electric field and the incident electric field. 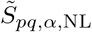 comes from interference between scattered fields, which is the weakest among three, and are nonlinearly related to the Stokes component.

In transparent biological specimen, most of the signals in the Stokes parameters come from the DC and the linear terms. We hence assume the contribution from the scattered field interference is negligible. Neglecting the nonlinear term and subtracting the DC term from 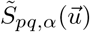, we deal only with the linear term as

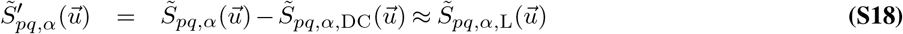

After this linearization, we see the DC-substracted Stokes component is a sum of filtered scattering potential tensor components, 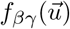 and 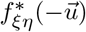, as seen in Eq. (S17). These corresponding filters are essentially weak object transfer functions presented in the scalar wave scattering theory (10–14). We introduce a set of notations to simplify their expressions as

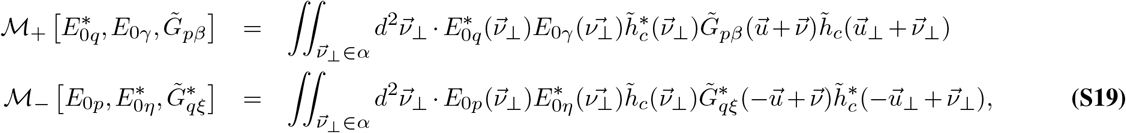

where 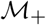 and 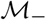 are mappings from the associated incident electric field components and dyadic Green functions to the actual transfer functions.

To relate the Stokes parameters to more physical interpretable quantities from the scattering potential tensor, we decompose the tensor components into the following forms

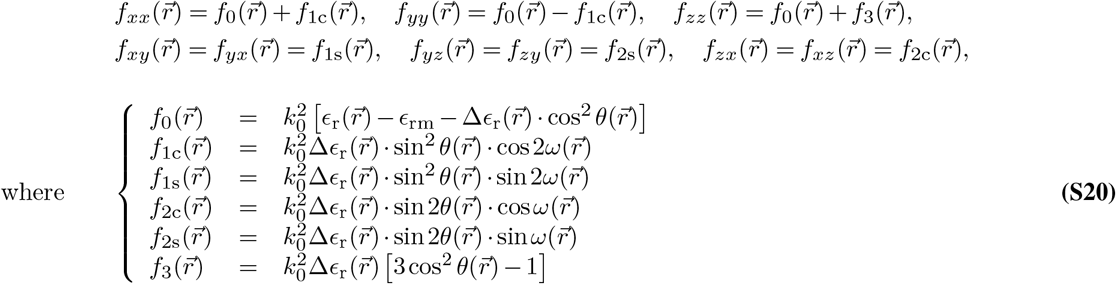

Here we assume 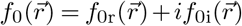 is a complex function carrying density (average refractive indices) information of the material 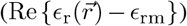 in the real part and the absorption information of the material 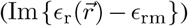 in the imaginary part. 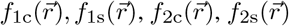, and 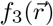 are assumed to be real functions (ignoring the effect of diattenuation), providing information of anisotropy 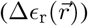, in-plane orientation 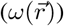, and inclination 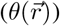.

By grouping all the linear contribution from the same tensor component in Eq. (S17) together, we arrive at this 7-unknown-variable linear equation to solve for

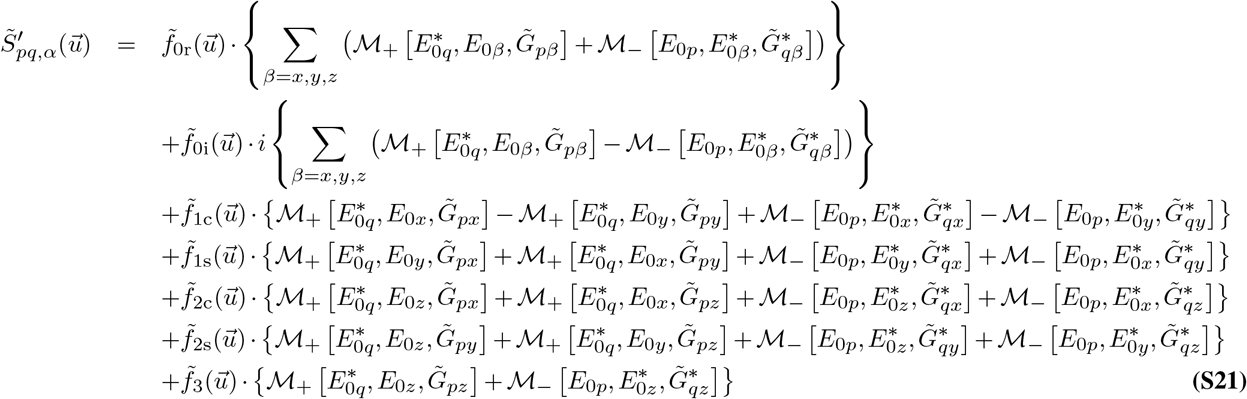

Each scattering potential tensor component has its transfer function to link to the Stokes component. However, the actual Stokes parameters in Eq. (S15) are a sum or a difference of the Stokes components. By merging all the contribution once again for *p,q* = *x, y,* we summarize our linearization of the Stokes parameters to scattering potential tensor in one equation as

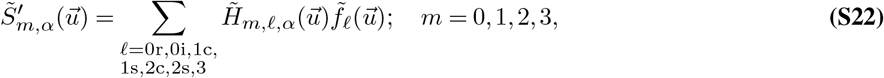

where 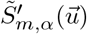 is the DC-subtracted Stokes parameter and 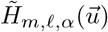 is the transfer function mapping from each tensor component *ℓ* to the *m*-th Stokes parameter under illumination pattern *α*. A detailed expression for these transfer functions are shown in the Appendix.

#### C.4: Reduction to 2D imaging model

When the thickness of a specimen is smaller than the depth of field of a certain imaging condition, acquiring data at one plane by changing the partially coherent illumination patterns enables extraction of 2D phase and anisotropic information at that plane. This 2D information extraction requires a slight modification of the transfer functions. In this 2D imaging case, we consider the scattering potential tensor to be concentrated in a single axial layer and is expressed as

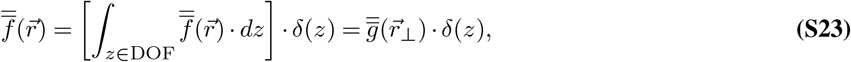

where 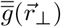 is the scattering potential tensor integrated in the *z*-direction over depth of field. We then substitute this expression for 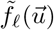 in Eq. (S22) and integrated in *u_z_*-direction to get a corresponding relationship in the 2D imaging case between 2D Stokes parameters and the integrated scattering potential tensor as

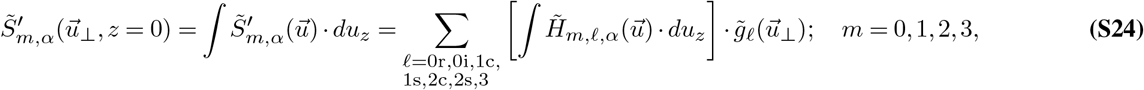

The transfer functions in 2D imaging case are simply a *u_z_*-integration of the previously established 3D transfer functions.

## Supplementary Note 2: Computing Stokes parameters from intensities

In fig. 1, we have demonstrated that the raw images from uPTI setup contain intensity modulation induced from specimen’s permittivity tensor. We have also established a forward model to describe the relationship between components of specimen’s permittivity tensor and the Stokes parameters of the output scattered light. Here we connect the missing link between the Stokes parameters of the scattered light to our measured intensity for further permittivity tensor retrieval.

### A: Calibration

Ideally, our polarization sensitive camera perfectly detects four channels of linearly polarized light oriented at 0°, 45°, 90°, and 135°. According to the definition of the Stokes vector (18), the measured intensity is linearly related to the Stokes parameters in the following form

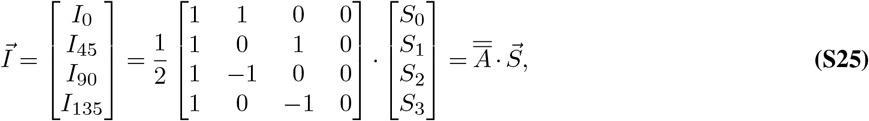

where 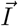 is the intensity vector of each polarization measurement, 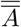 is the instrument matrix (19) of the optical system, and 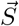 is Stokes vector of the light. Applying the inverse of the known instrument matrix, 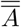, on the measured intensity vector, 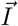, allows us to get the first three Stokes parameters for further deconvolution. However, the actual optical setup has multiple components (e.g. beam splitting prisms) with unexpected polarization effects. Such components provide non-ideal values in the instrument matrix. Thus, a calibration procedure to measure the instrument matrix is essential. To calibrate the unknown instrument matrix, we adopt the approach in (19). We place a linear polarizer mounted on a rotational stage at the specimen plane and rotate it to collect corresponding intensity variations in four polarization channel. According to Mueller matrix formulation, we model the intensities collected at angle *ω*_LP_ of the linear polarizer and retrieve the instrument matrix. Additional to the previous method, we demonstrate extra calibration of the polarization state of the incident light.

Here we show how we model our calibration through Mueller formulation. Assuming the incident light has the Stokes parameters of 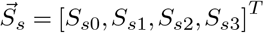, the generated polarization state of light after the linear polarizer aligned with angle *ω*_LP_ in Stokes parameters is calculated

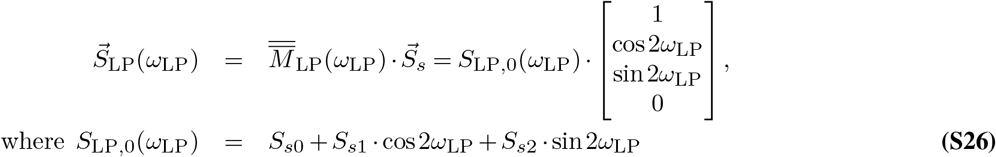

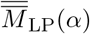 is the Mueller matrix of a linear polarizer aligned at angle *ω*_LP_. This formula shows that the total energy of the generated state, *S*_LP,0_(*ω*_LP_), will be a constant along *ω*_LP_ if the input light is unpolarized or circularly polarized. This total energy can vary if our input light has some linear polarization components. We estimate the total energy of the generated light by summing over intensities of four polarization channel. Through a Fourier series fitting, we obtain the Stokes parameters of the incident light and use it for later deconvolution.

Sending this generated polarization state into the detection optical system, we get the measured intensity through the multiplication of the instrument matrix on the generated Stokes parameters

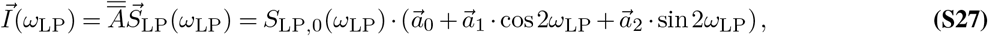

where 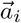 is the *i* + 1-th column of the instrument matrix 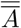. If we normalize the measured intensity with the total energy of the generated light, *S*_LP,0_(*ω*_LP_), we observe that the first three columns of instrument matrix are the Fourier series coefficients of the normalized intensities along *ω*_LP_. We conduct another fitting to estimate our instrument matrix. With the first three columns of the instrument matrix, we then used it to obtain the first three Stokes parameters of the light as

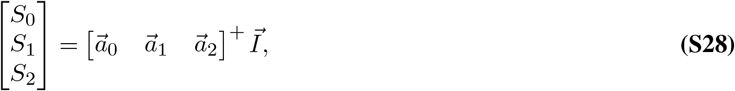

where *A*^+^ denotes a pseudo-inverse of the matrix *A*.

### B: Background correction

This Stokes parameters may still subject to leftover polarization effects that produce slowly varying background. In addition, the deconvolution works with the DC-subtracted Stokes parameters according to Eq. (6). With the derivation described next, we obtain the DC-subtracted Stokes parameters with the following operations

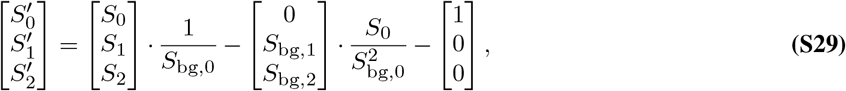

where *S*_bg,i_ is the *i*-th background Stokes parameters obtained when imaging at an empty field of view. When the transmission modulation of the specimen is relatively weak, *S*_0_/*S*_bg,0_ ≈ 1 and this is simply removing the DC components of the Stokes parameters. These operations are done to all sets of polarization acquisitions with different illumination patterns and then the deconvolution takes these DC-subtracted Stokes parameters for the retrieval of uniaxial permittivity tensor components.

### C: Derivation of the Stokes background correction

Since background variation is usually slowly varying, we adopt a non-diffraction-based Mueller calculus to develop our background removal scheme. An anisotropic sample with transmittance *t_s_* = |*e^μ+iϕ^*|^2^, in-plane orientation of *ω*, out-of-plane orientation of *θ*, apparent retardance *ρ*_ap_(*ω, θ*) can be modeled with an Mueller matrix as

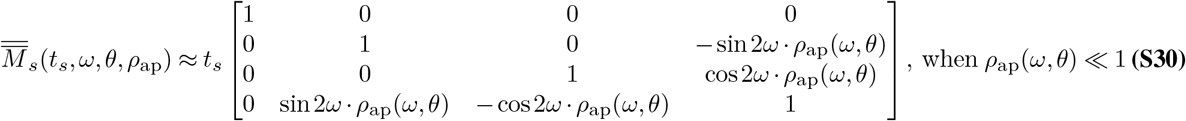

Sending in the circularly polarized light with the Stokes parameter of 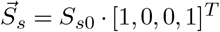, where *S*_*s*0_ is the energy of the incident light, the output Stokes parameters are obtained as

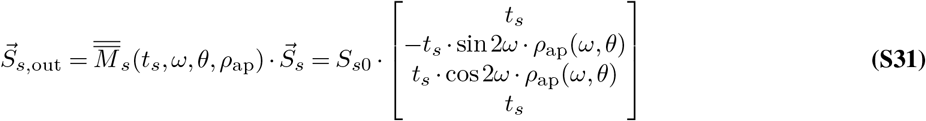

This is the Stokes parameters we get assuming we do not have extra background polarization effects from the imaging optics in the microscope. When the imaging optics are not perfect, slowly varying polarization modulation appears to mask the weak anisotropy signal out. To find a way to remove this effect, we model the background polarization effect to be an additional transparent retarder with the Mueller matrix of 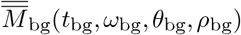. With this background Mueller matrix, the overall output Stokes parameters are obtained as

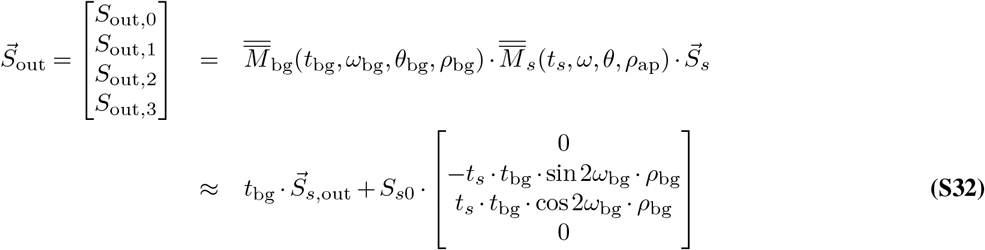

This overall Stokes parameters contain contributions from the specimen and the background effect. To separate these effects, we acquire the background Stokes parameters by imaging an area without the specimen and we get

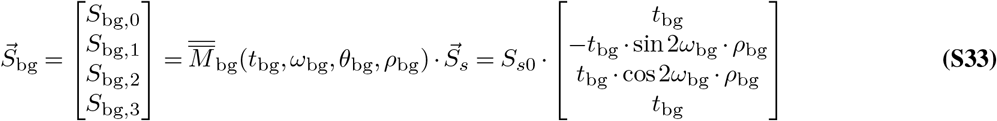

Comparing Eq. (S32) with Eq. (S33), we then come up with a correction to get the corrected Stokes parameters as

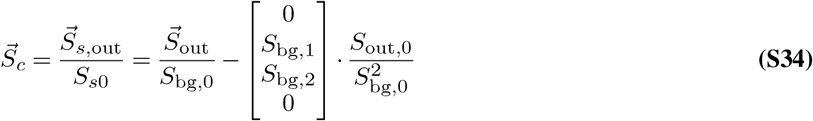

By doing this correction, we also remove the DC components of the Stokes parameters in *S*_out,1_ and *S*_out,2_. The only remaining Stokes parameters with the DC-component is the *S*_0_ component of 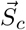. With the additional subtraction, we arrive at the final DC-subtracted Stokes parameters that we used for the deconvolution

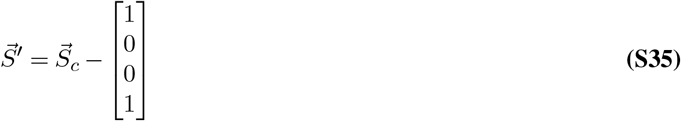

We apply these corrections on all sets of intensities from each illumination patterns before the deconvolution algorithm.

## Supplementary Note 3: Transfer functions and choice of illumination pattern

The choice of the illumination patterns for our experiment is modified from past literature of 3D differential phase contrast (3D DPC) microscopy (14) and 3D polarized light imaging (3D PLI) (20–22). To get 3D phase information, 3D DPC exploits illumination patterns of 4 rotating half-circles with NA matched in illumination and detection, which is an extension from 2D DPC (11). In 3D PLI, 3D orientation and the principal retardance of the optically anisotropic material (axons in human brain) are extracted with brightfield illumination, polarization-sensitive detection, and a tilting stage. In their acquisition, one untilted condition followed by four tilted conditions in 90°-separated orientations are used. The 4 tilted conditions with brightfield illumination is similar to untilted stage illuminating with 4 off-axis illuminations in rotating orientations as what it has been done in 3D DPC. The one untilted polarization measurement plays an important role for 3D orientation retrieval as shown in (20). From these rules of thumb, it is reasonable to use combination of brightfield illumination pattern plus several off-axis illumination patterns in a polarization-sensitive detection system for acquiring 3D phase, principal retardance, and 3D orientation information.

There are multiple combinations of patterns that could be used to get the information of permittivity tensor. Different illumination patterns result in different shapes of the transfer function in Eq. (6), which affects noise robustness over 3D spatial frequencies. By visualizing these transfer functions, we get a sense of how well specific information is transferred with the illumination patterns. Figure 1-supplement 2 compares the absolute sums of the transfer functions over illumination patterns for two possible combinations of illumination patterns that are deduced from past literature, which describe how individual scattering potential tensor components couple into the measured intensities as a function of their spatial frequencies. In summary, *S*_0_ carries phase (*ℓ* = 0r) and absorption information (*ℓ* = 0i), while *S*_1_ and *S*_2_ carry anisotropy information (*ℓ* = 1c, 1s, 2c, 2s, 3). For both sets of illumination patterns, phase, absorption, and projected anisotropy information (*ℓ* = 1c, 1s) transfer relatively well compared to the last three components (*ℓ* = 2c, 2s, 3). The last component of the scattering potential tensor, which contains information of optic sign, is the most poorly-transferred in both cases. This is why the optic sign estimation is noisy. Comparing the energy of the transfer function with these two sets of illumination, we find that the second set of illumination patterns generates more uniform energy of transfer function for all the scattering potential components. Off-axis illumination with more orientations tends to give better information transfer and better noise robustness across different spatial frequencies, but it takes more time for each experiment. To balance the trade-off between the noise performance and the acquisition time, we choose the second set of illumination patterns for our experiment, which is 8 off-axis max-NA sector patterns followed by 1 on-axis half-NA brightfield pattern. Note that these are not the only choices for proper permittivity tensor retrieval. Other combinations of patterns are possible, but it would require further investigation to determine the optimal one as that is done in phase imaging community (23, 24).

## Supplementary Note 4: The formation of laser-induced modifications in fused silica

When an intense femtosecond laser pulse is focused into transparent material, e. g. silica glass, high-order nonlinear absorption allows the energy to be deposited predominantly within the focal volume, producing a local permanent refractive-index modification. Although the process of energy absorption is now well understood, the actual formation of the directly written structures is still controversial.

Depending on the level of laser intensity, one can induce any of four qualitatively different types of material modification: positive refractive-index change (type 1) at relatively low intensity; birefringent regions with the evidence of nanograting formation (type 2) (25, 26) at intermediate intensity and voids (type 3) at high intensity. The transition threshold from one kind of structure to another depends on the laser writing parameters such as pulse duration, wavelength and irradiation time. The nanograting type (type 2) of modifications is observed under SEM as periodic subwavelength arrangement of nanoplanes with features as small 20 nm. Recently new type of birefringent modification has been observed as random arrangement of prolate-shape nanopores at transition from the type 1 to type 2 and coined as type X modification (27). The orientation of both nanogratings and elongated nanopores can be controlled by the polarization of the writing laser and results in form birefringence with slow axis oriented perpendicular to the polarization of writing laser and along the nanoplanes or long axis of prolate-shape nanopores. The induced birefringence is negative with the value for type 2 close to the birefringence in quartz crystal, and for type X of about 4 times lower value.

Birefringent structures produced with type X modification reveal ultra-low scattering loss and were observed when writing with infrared 1030 nm laser wavelength. Writing with shorter wavelength, such as 515 nm is advantageous for imprinting with high spatial resolution. However, only type 2 birefringent structures were revealed with the Abrio birefringence imaging system, which can only measure the average change of birefringence along axial direction. The structure of imprinted birefringent modification, e. g. the possible coexistence of different types of modification along the axial direction shown in fig. 2C, in principle can be characterized by SEM. However non-destructive method of 3D birefringence imaging, such as uPTI, is more convenient and resulted in the observation of type X structures in the tail of birefringent modification. This observation allows improving resolution of low loss birefringent structuring in silica glass.

